# MORPH Predicts the Single-Cell Outcome of Genetic Perturbations Across Conditions and Data Modalities

**DOI:** 10.1101/2025.06.27.661992

**Authors:** Chujun He, Jiaqi Zhang, Munther Dahleh, Caroline Uhler

## Abstract

Modeling cellular responses to genetic perturbations is a significant challenge in computational biology. Measuring all gene perturbations and their combinations across cell types and conditions is experimentally challenging, highlighting the need for predictive models that generalize across data types to support this task. Here we present MORPH, a MOdular framework for predicting Responses to Perturbational cHanges. MORPH combines a discrepancy-based variational autoencoder with an attention mechanism to predict cellular responses to unseen perturbations. It supports both single-cell transcriptomics and imaging outputs and can generalize to unseen perturbations, combinations of perturbations, and perturbations in new cellular contexts. The attention-based framework enables inference of gene interactions and regulatory networks, while the learned gene embeddings can guide the design of informative perturbations, as demonstrated in two applications. Overall, MORPH is a flexible tool for optimizing perturbation experiments, enabling efficient exploration of the perturbation space to advance understanding of cellular programs for fundamental research and therapeutic applications.

## 1 Introduction

Single-cell sequencing and imaging technologies have revolutionized our understanding of cellular heterogeneity, enabling detailed characterization of various cell states and functions at scale [1]. Coupled with CRISPR-based perturbation platforms and multiplexed barcoding strategies, these technologies now allow systematic, high-dimensional mapping of the heterogeneous cellular responses to individual genetic perturbations. Several methods combine genetic perturbation with single-cell RNA sequencing (scRNA-seq) as a readout for the gene expression changes [2–7], while optical pooled screens capture cellular phenotypes through imaging data [8–10]. Understanding these perturbation-induced effects not only provides insights into the structure of gene regulatory networks but also has significant implications for biomedical research, such as identifying potential therapeutic targets that could drive cells to a desired state [11–13] and optimizing combination therapies [14–16].

While the availability of single-cell perturbation data is growing, experimentally testing all individual gene perturbations – and their combinations – across all possible cell types, states, and diseases contexts remains unfeasible. Therefore, there is a need for computational methods that leverage the existing data to predict perturbation effects across different conditions and data types [17, 18]. Such models would be crucial for efficiently exploring the vast perturbational space and prioritizing the most informative experiments to perform. The urgency of this need is reflected in two recent community efforts: the Cell Perturbation Prediction Challenge launched in 2023 with a first competition focused on predicting gene perturbation effects on T-cell state distributions [19]; and the Virtual Cell Challenge, launched in 2025 to benchmark cross-context generalization at scale [20].

Current methods for this task [21, 22] perform well in approximating the state of perturbed cells on average but struggle to capture the full distribution of cellular responses, and often rely on randomly pairing control and perturbed cells to address the inherent unpaired nature of high-throughput sequencing data—a fundamental challenge arising from the destructive nature of these assays. Additionally, many existing methods assume that the effects of gene perturbations are additive [23, 24], limiting their ability to capture complex, non-additive interactions. More recently, methods developed concurrently with this work [25], have demonstrated that out-of-distribution prediction and transfer to unseen cell lines are achievable for transcriptomic data. However, the ability to predict the perturbation outcomes measured in modalities other than transcriptomics remains largely unexplored, as does the question of model interpretability: the extent to which learned representations reflect biologically meaningful gene relationships rather than opaque mappings.

To address these challenges, we developed MORPH, a MOdular framework for predicting Responses to Perturbational cHanges. Its modular design enables MORPH to adapt seamlessly to various data modalities, such as transcriptomic or imaging data. Given single-cell data on control cells and prior knowledge of gene perturbations, MORPH leverages a conditional variational autoencoder with an attention mechanism to predict the distribution of cellular responses to unseen perturbations either in transcriptomic or imaging modality. We demonstrate that MORPH generalizes to unseen single and combinatorial perturbations, transfers to new cell lines using only control cell data, and guides experimental design by identifying informative perturbation targets. Building on our previous theoretical work [26], which proved that causal effects can be reliably estimated even when datasets are not directly matched cell-by-cell, we show that MORPH’s attention mechanism captures functional—rather than merely co-expression—relationships among genes, enabling interpretable inference of gene regulatory structure. We validated MORPH on five Perturb-seq datasets [5, 7, 19] and an image-based optical pooled screen [10], outperforming available baselines throughout. In two applications, we show how MORPH’s different capabilities can be used synergistically for biological discovery. In the first, we leveraged the OPS dataset on HeLa cells infected with the Ebola virus [10] and in a simulated experiment, we showed that MORPH successfully nominated the top perturbation that shift the cell infection stage toward earlier stages. In the second, by pretraining on public data and fine-tuning on an in-house CD8 T-cell Perturb-seq dataset, MORPH successfully predicted a gene target (*Batf*) that could potentially enhance anti-tumor immunity by increasing the proportion of progenitor-like T cells, a result validated in an independent dataset.

Overall, our results highlight MORPH’s versatility and potential to guide the design of future perturbation experiments to enable more efficient and targeted perturbation screens.

## 2 Results

### MORPH enables the prediction of cellular responses to genetic perturbations at single-cell level

MORPH is designed to predict the effect of a genetic perturbation on an individual cell. We represent a cell as a vector *X*^·^ encoding any high-dimensional readout, such as gene expression profiles from RNA sequencing or morphological features extracted from imaging. We encode prior knowledge about a gene *g* of interest (such as pathway associations, gene interactions, etc.) and its perturbation effect as an embedding vector **v**^*g*^. This embedding allows MORPH to relate unseen perturbations to previously seen ones, based on the intuition that perturbations with similar biological properties tend to have similar effects. Crucially, these embeddings are derived from external data sources and do not use the target Perturb-seq data, ensuring no leakage of single-cell perturbation–response information from held-out perturbations. Since these priors are defined at the level of individual genes and do not encode multi-gene perturbations, MORPH models combinatorial perturbations by composing individual latent gene representations within its architecture (Figure 1a).

**Fig. 1.**
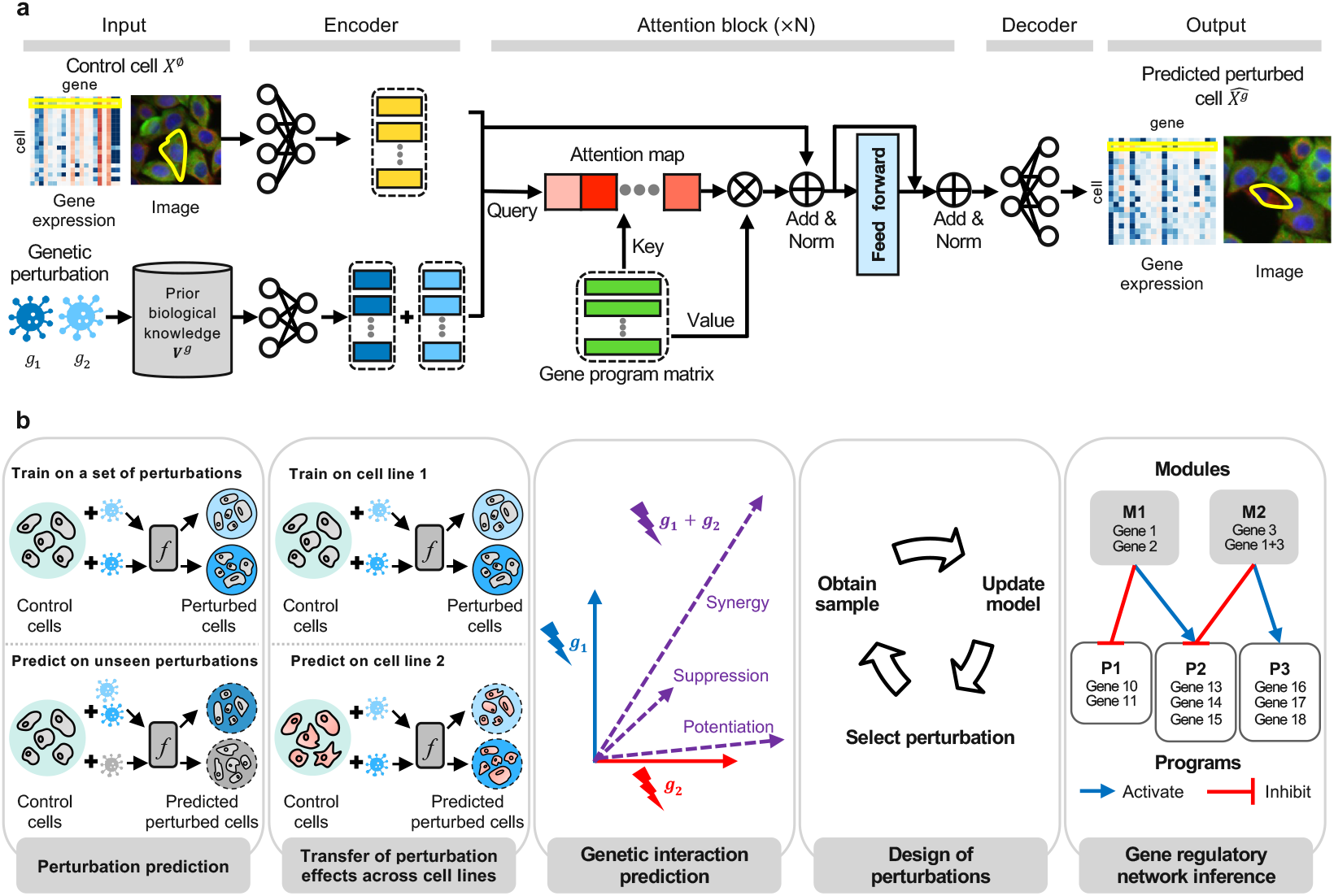
Overview of MORPH and its applications. **a**, Model architecture. For each pair of control cell and genetic perturbation, the model maps them into latent representations using separate encoders. In the latent space, attention modules dynamically identify the most relevant features for the given perturbation. The resulting attention output is then passed to a decoder to generate the predicted perturbed cell. We demonstrate MORPH on both transcriptomic and imaging modalities. **b**, Downstream applications. MORPH can be applied to a variety of tasks, including predicting unseen genetic effects, transferring perturbation effects across cell lines, predicting genetic interactions, designing perturbation experiments, and inferring gene regulatory networks.

Formally, given a control cell *X*^∅^, an embedding **v**^*g*^ for gene *g*, MORPH learns a function *f* that maps the pair (*X*^∅^, **v**^*g*^) to the predicted perturbed cell 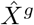, i.e., 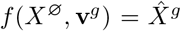. For each gene perturbation in the dataset, MORPH compares the predicted perturbed distribution of 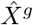 to the actual perturbed distribution of *X*^*g*^, enabling it to learn the mapping *f* from unpaired data. This is achieved using maximum mean discrepancy (MMD) between the predicted cell distribution 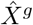 and the actual cell distribution of *X*^*g*^ (Methods). If combinatorial perturbation data is available (e.g. on genes *g*_1_ and *g*_2_), this is generalized by comparing 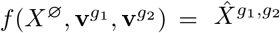 to the actual perturbed distribution of 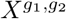.

To realize the mapping *f*, MORPH adopts a variational autoencoder (VAE)-based architecture. A control cell *X*^∅^ and gene perturbation embedding vector **v**^*g*^ are first embedded into latent spaces using two separate encoders. To enhance interpretability and reliability, these latent representations are passed through an attention mechanism that guides the model to focus on the most relevant aspects of the input data, before being decoded into the predicted perturbed cell 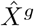. All encoders and decoders are implemented as multi-layer perceptrons (MLPs) (Figure 1a, Methods). The model is trained using a variational autoencoder loss on the control cells *X*^∅^ and MMD between the predicted control/perturbed cell distributions and the actual control/perturbed cell distributions (Methods). Using MMD also for the control cells promotes attention score changes that more faithfully reflect the magnitude of perturbation effects (Supplemental Fig. 1).

The design of our attention mechanism was motivated by a previous study [2] that had represented the effects of genetic perturbations using a bipartite regulatory network, where perturbations influence the expression of regulated genes. Clustering perturbations based on shared regulated genes reveals *perturbation modules* — groups of perturbations that produce similar cellular effects. Similarly, clustering genes based on their co-regulation by perturbations yields *gene programs* — sets of genes that respond in a coordinated way. These perturbation modules and gene programs reflect the underlying structure of gene regulation. MORPH integrates a cross-attention mechanism to capture the regulation from perturbations to gene programs (Methods). It constructs queries by concatenating the latent vectors obtained by encoding the control cell and the perturbation embedding. These queries attend to a learned gene-program matrix that serves as keys and values, producing attention scores that modulate the influence of each gene program on the final prediction through the network’s non-linear feed-forward layers (Figure 1a). To biologically interpret this non-linear process, we utilized an end-to-end probing procedure that maps these programs directly to gene expression (Methods). The gene-program matrix is shared across all cells of the same type and remains fixed at inference time, under the assumption that gene programs are relatively stable within a given cell type, while their activation depends on the specific perturbation and the initial state of a cell.

These design choices enable MORPH to address fundamental questions about genetic perturbations. We demonstrate that MORPH can predict cellular responses to unseen perturbations in both transcriptomic and imaging data, highlighting its versatility in handling various data modalities. Moreover, it can transfer perturbation effects across cell types, extrapolate genetic interaction types, optimize the design of genetic perturbations, and facilitate the investigation of genetic regulatory networks (Figure 1b). In the following sections, we evaluate each of these capabilities and illustrate how they can be combined for practical applications.

### MORPH generalizes to predict the effect of unseen single-gene Perturb-seq experiments with high accuracy

We evaluated the ability of MORPH to predict cellular responses to unseen single-gene perturbations using single-cell Perturb-seq data from the three screens in [7]: K562-EG, a screen of 2,033 essential-gene perturbations across over 310,000 K562 cells; RPE1-EG, a screen of 2,264 perturbations across over 240,000 RPE1 cells; and K562-GW, a genome-wide perturbation screen comprising 9,823 individual gene perturbations across over 1,900,000 K562 cells. The model was trained separately on each dataset, and predictions were compared against held-out perturbations not seen during training (Figure 2a).

**Fig. 2.**
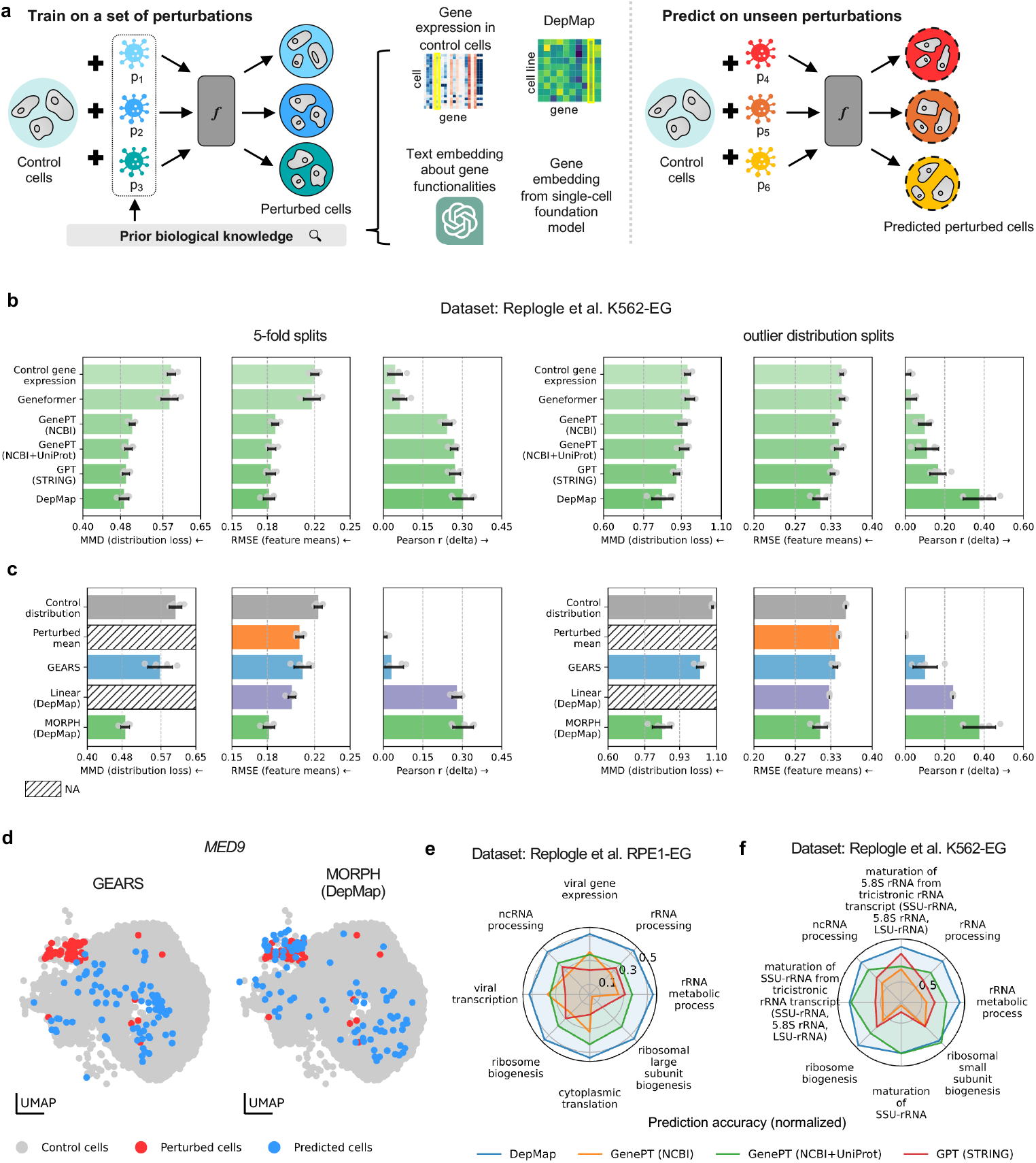
Predicting responses to unseen single-gene perturbations. **a**, Workflow of single-gene perturbation predictions. Different sources of prior knowledge were explored, such as gene embedding vectors derived from single-cell foundation models and perturbation-specific databases like DepMap. **b-d**, Evaluation results on the K562-EG dataset. Sources of prior knowledge (**b**) and models (**c**) were evaluated using a distributional distance (MMD), average RMSE between the mean predicted and observed gene expression profiles, and Pearson correlation between the mean predicted and observed perturbation gene expression changes relative to the training-set perturbed mean. Pearson correlation values were clipped at 0 for visualization purposes. Metrics were calculated using the top 50 differentially expressed (DE) genes for each perturbation. Evaluations were performed across 5-fold cross-validation splits and outlier distribution splits. Hatched bars indicate metrics that are not applicable. MMD is not reported for Perturbed mean and the linear model because they predict only mean expression profiles and do not generate single-cell distributions. Arrows indicate the direction of improvement (*→* higher is better; ← lower is better). **d**, UMAPs of observed perturbed cells and cells predicted by MORPH (on the right) and the current state-of-the-art model that predicts single-cell level response, GEARS (on the left). **e-f**, Spider plots summarizing the prediction accuracy using each prior knowledge base across the top 8 enriched gene sets. These gene sets were identified through gene set enrichment analysis on the union of genes where one prior outperformed the other. Prediction accuracy was computed as 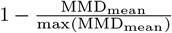, where MMD is the mean prediction loss (MMD) for each gene set under a given prior, and max(MMD_mean_) is the highest mean prediction loss across all gene sets.

We tested the model on two types of data splits (Figure 2b-c and Supplemental Figure 2). The first uses a standard 5-fold cross-validation, where the dataset was divided into five equal sets of perturbations. In each iteration, one set of perturbations served as the test set, and the remaining four were used for training. This process was repeated for all five folds, and the performance metrics were averaged to evaluate the model. The second, termed the “outlier distribution split”, specifically included perturbations producing phenotypes that were most distinct from the control distribution in the test split. This was achieved by performing Leiden clustering on the pseudo-bulk profiles of each perturbation and identifying the clusters whose centers were farthest from the cluster that contained control, measured by Euclidean distance between averaged bulk expressions. This split was designed to assess the ability of the model to generalize to unseen perturbations that induce significantly distinct profiles, providing a stricter stress test for generalization beyond standard cross-validation (Supplemental Figure 2).

To assess performance, we used three primary metrics (Figure 2b-c, Methods). Root mean squared error (RMSE) was used to quantify the mean difference between predictions and observations; Pearson correlation (delta) was used to measure the agreement between predicted and observed mean changes relative to the mean of perturbed training cells [27]; Spearman correlation was used as a complementary rank-based metric. In Supplemental Figures 3–6, we also reported the retrieval rank, defined as the fraction of other perturbations whose correlation with a predicted effect does not exceed its self-correlation (Methods), yielding scores between 0 and 1. In addition to these mean-based metrics, we calculated the maximum mean discrepancy (MMD) to evaluate *distributional* differences between predicted and true effects in gene expression observed in individual perturbed cells. Lower MMD values indicate closer alignment between the two cell distributions. This metric offers more fine-grained information than metrics that compare only mean gene expression. We calculated these metrics using the top 50 differentially expressed (DE) genes for each perturbation for a more focused evaluation (Figure 2 and Supplemental Figures 4-5) as well as using all genes (Supplemental Figure 6).

**Fig. 3.**
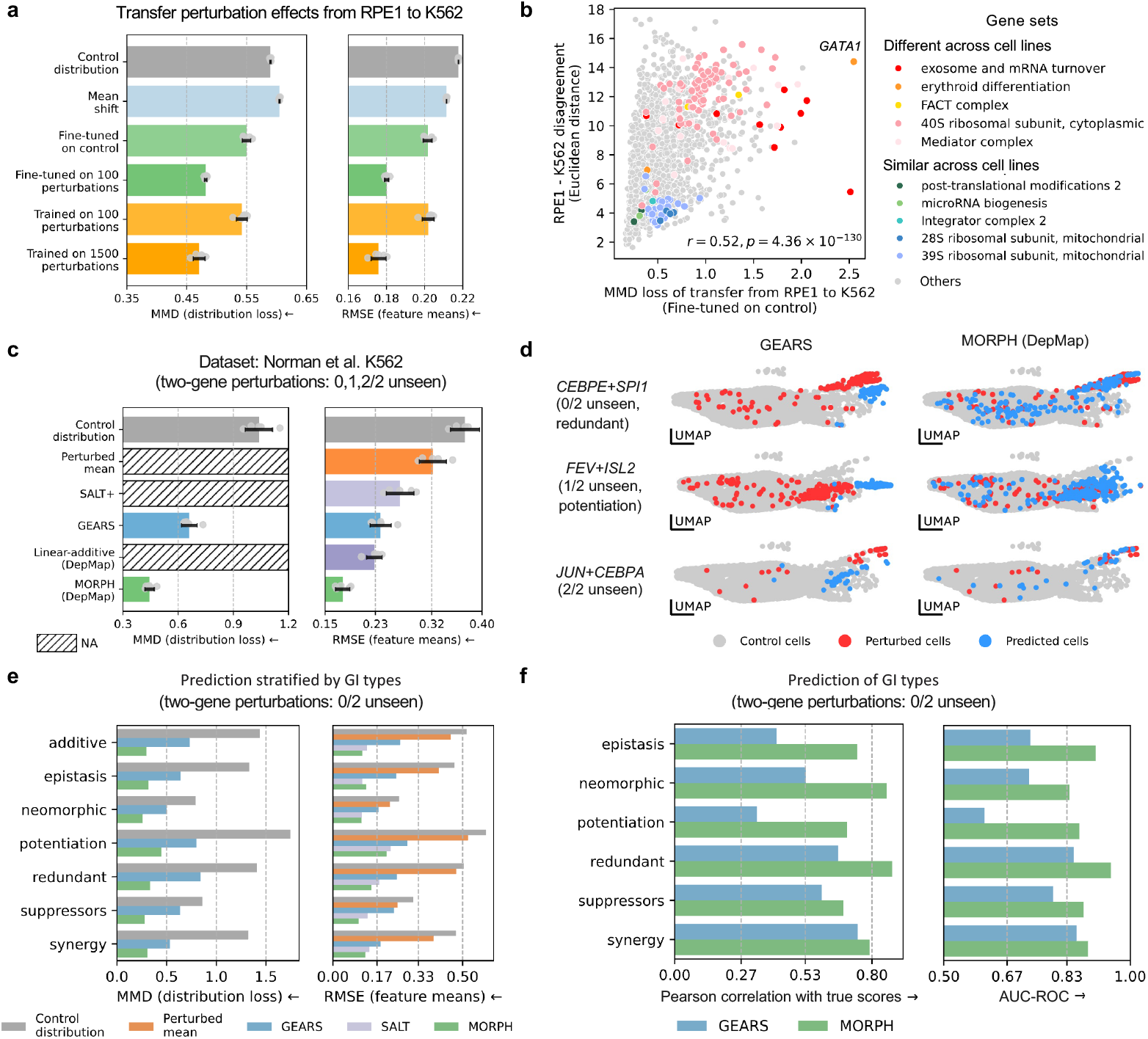
MORPH effectively transfers perturbation effects across cell lines and predicts double-gene perturbations. **a**, Evaluation of model performance in transferring perturbation effects from RPE1-EG to K562-EG [7], using MMD and RMSE metrics. Baseline models include the control distribution and mean shift, which applies the same mean change vector from control in the training cell line to the test cell line. The figure compared the performance of different training strategies of MORPH, including a model trained on the training cell line and fine-tuned using only control cells from the test cell line (“Fine-tuned on control”), a model fine-tuned on 100 perturbations from the test cell line (“Fine-tuned on 100 perturbations”), a model trained from scratch on 100 perturbations in the test cell line (“Trained on 100 perturbations”), and a reference model trained on 1,500 (75%) perturbations in the test cell line. **b**, Comparison of the performance of the model fine-tuned on control cells (K562), measured using MMD, and the cell line disagreement, calculated as the Euclidean distance between shift vectors for each perturbation in training (RPE1) and test (K562) cell lines. Pearson correlation = 0.52, p-value *<* 0.05. Genes are colored by gene sets representing varying levels of similarity across cell lines. **c**, Box plots showing model performance in predicting cellular responses to double-gene perturbations on the Norman dataset [5], averaged across scenarios where 0, 1, or 2 genes in the pair are unseen during training. Hatched bars indicate metrics that are not applicable. MMD is not reported for the Perturbed mean, SALT+ and linear-additive models, which predict only mean expression profiles. **d**, UMAP visualization of top perturbations with maximal effects in the test set. **e**, Analysis on double-gene perturbations where both single genes were observed during training, stratified by gene interaction types. Box plots comparing model performance across different interaction types. **f**, Box plots evaluating model performance in predicting genetic interaction types. Evaluation metrics include Pearson correlation and the area under the receiver operating characteristic curve (AUC-ROC). These metrics are computed based on gene interaction scores derived from model predictions and corresponding scores from observed data. AUC-ROC values indicate classification performance, where a random classifier would achieve an AUC-ROC of 0.5. Arrows indicate the direction of improvement (*→* higher is better; ← lower is better).

**Fig. 4.**
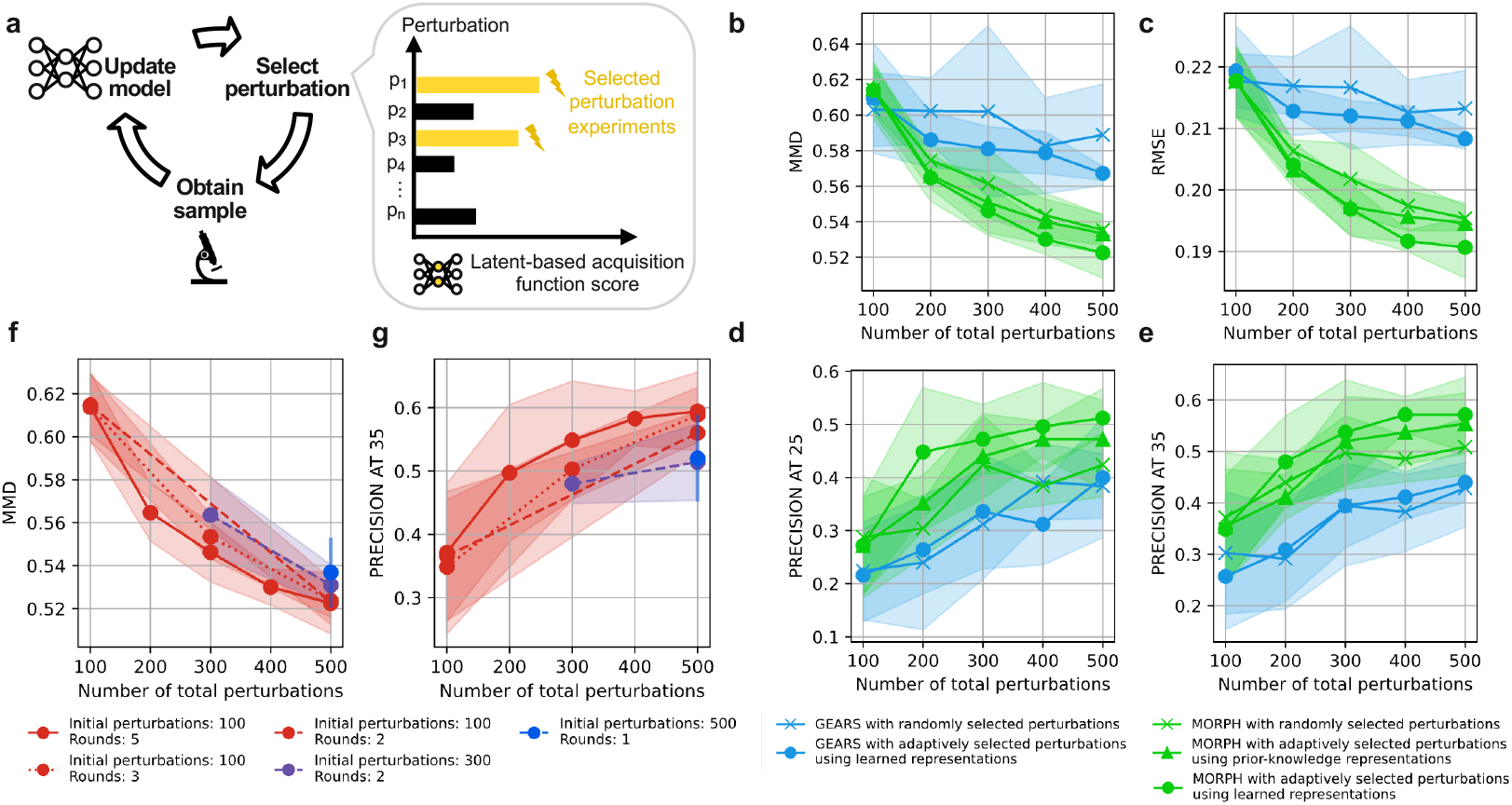
MORPH can be used for optimal design of perturbations. **a**, Iterative perturbation design framework. Perturbations are selected via an acquisition function, experimentally screened, and used to update the model with newly acquired data for the next round. **b-e**, Line plots comparing acquisition strategies and models for efficiently covering the perturbation space as measured by prediction over a held-out test set. The green line with ‘X’ markers represents MORPH with randomly selected perturbations. The green line with circle markers corresponds to MORPH using an adaptive strategy that selects perturbations with the highest predicted loss based on its learned latent representations. The green line with triangle markers shows a variant of MORPH that selects perturbations using a fixed, prior-based latent space. For comparison, the blue lines with ‘X’ and circle markers show GEARS with randomly selected and adaptively selected perturbations based on its learned latent representations, respectively. Lines indicate mean performance and shaded regions show *±* 1 standard deviations across 5 runs with different random seeds. Evaluation metrics include MMD (**b**) and RMSE (**c**) calculated on the top 50 DE genes on a set of perturbations withheld for testing. **d-e** shows the precision of using the predictions to identify the top 25 and 35 perturbations that shifted cells farthest from the control state. **f-g**, Line plots showing performance improvements of MORPH with adaptively selected perturbations using learned representations across different allocations of initial perturbations and rounds, under a fixed experimental budget. Shaded regions show *±*1 standard deviations across 5 runs using different random seeds.

**Fig. 5.**
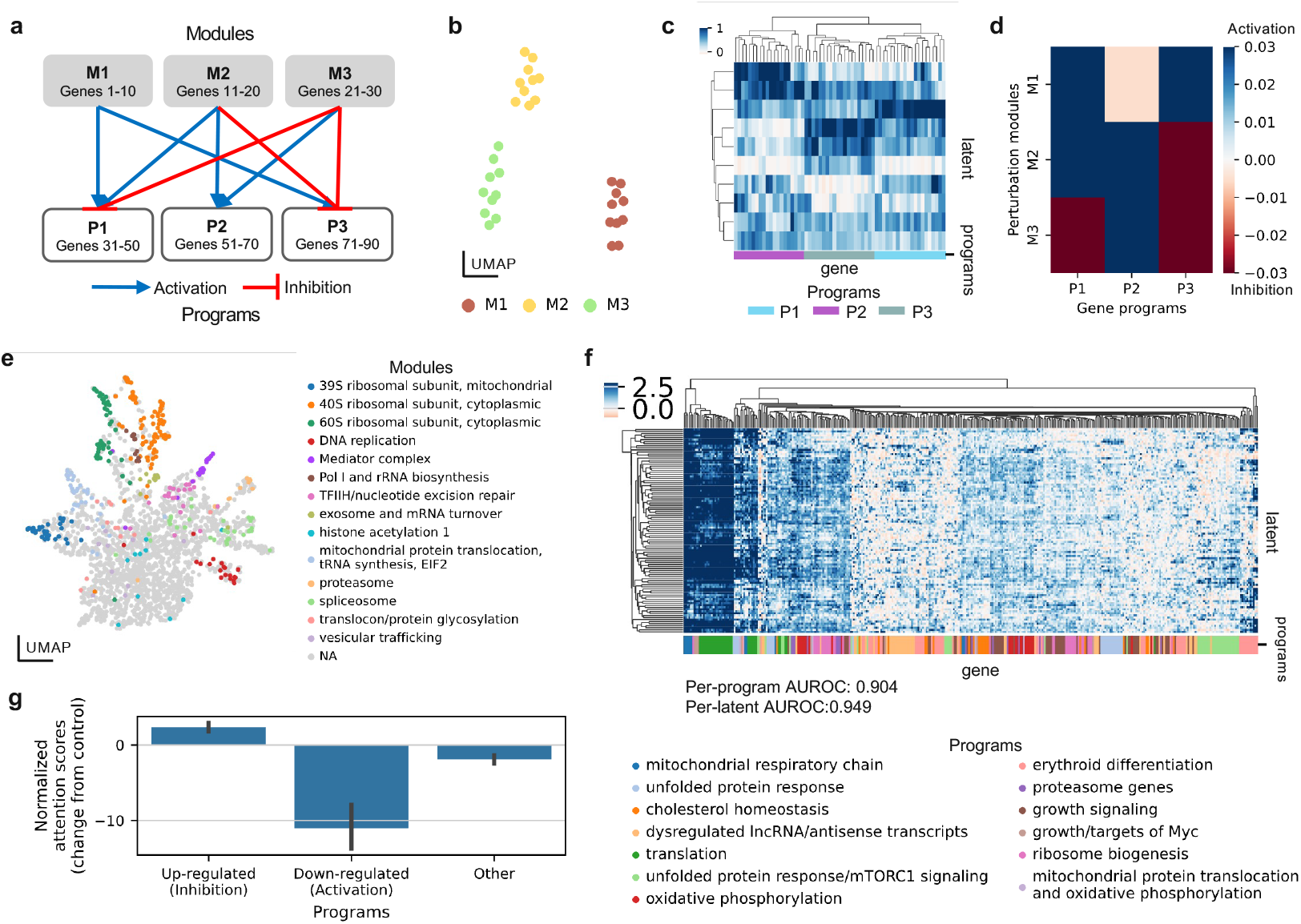
MORPH’s attention-based framework enables gene regulatory network inference. **a**, The gene regulatory network used to generate the simulated data, where perturbation modules (M) are clusters of genes that induce similar effects when perturbed and gene programs (P) are genes that exhibit similar response to perturbations. **b**, UMAP visualization of the learned perturbation latent space colored by perturbation modules on the simulated data. **c**, Hierarchically clustered genes into programs using the learned mapping from latent representations to genes, colored by true gene programs, on the simulated data. **d**, Heatmap representing the recovered gene regulatory structures inferred from the learned attention maps on the simulated data. **e-g**, Same analysis for MORPH trained on [7]: **e**, UMAP of the learned perturbation latent space colored by perturbation modules reported in [7]; **f** Hierarchically clustered heatmap of a subset of the gene–latent attribution matrix **M**, in which rows represent latent programs and columns represent genes. For visualization, we display only genes annotated in curated gene programs from [7]; annotated gene programs are indicated by the color bar. To quantify alignment between learned latent programs and annotated gene sets, we computed the area under the receiver operating characteristic curve (AUROC), treating attribution scores in **M** as continuous predictors and gene-set membership as binary labels. AUROC values were calculated using all genes. Because latent programs are learned without predefined correspondence to annotated gene sets, we applied a best-match strategy to compute per-latent and per-program AUROC scores (see Methods). Numerical AUROC values are indicated. **g**, box plots of rank-normalized attention score changes for reported up- and down-regulated programs post-perturbation from [7].

**Fig. 6.**
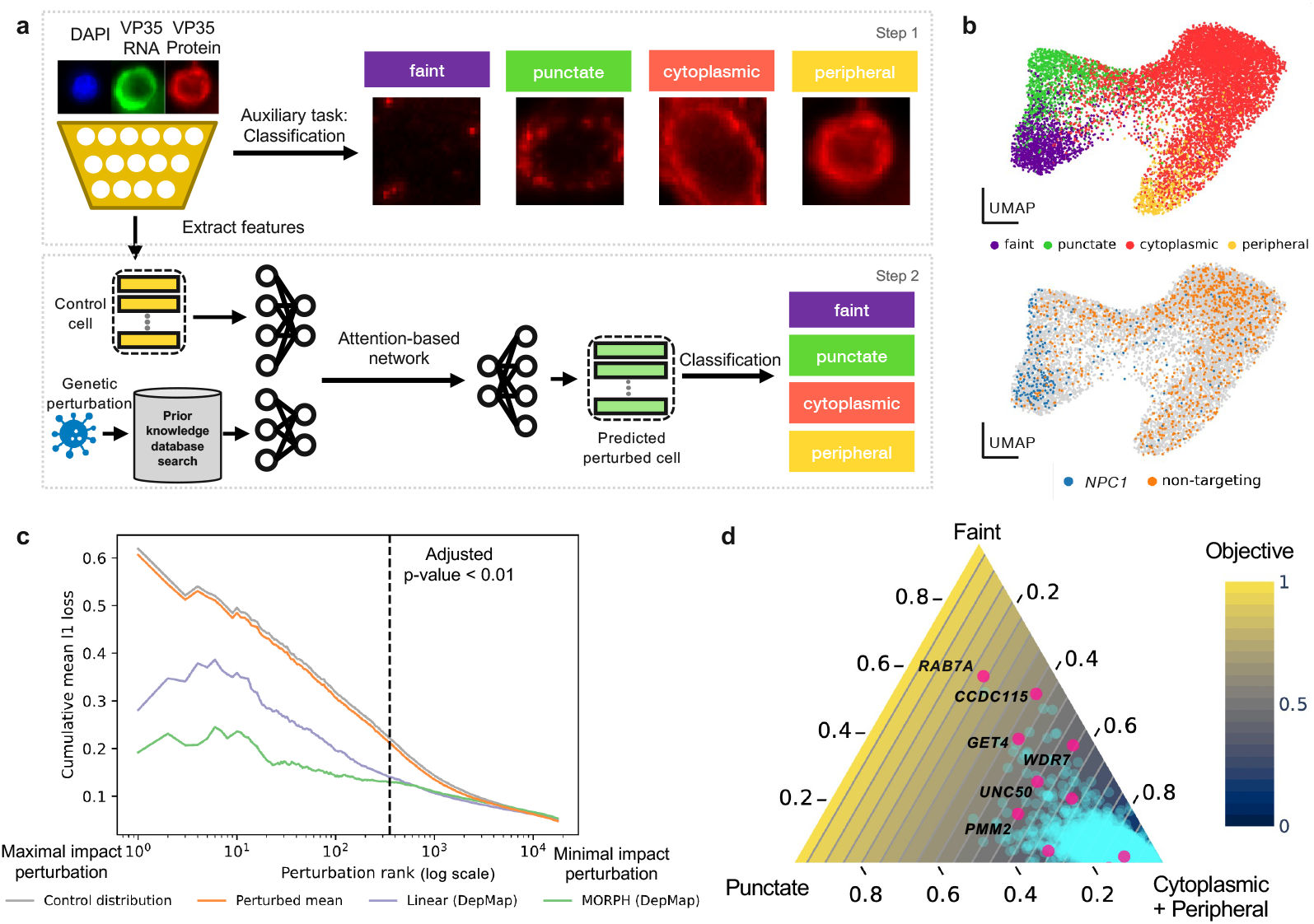
Application to imaging modality for perturbation outcome prediction using optical pooled screens. **a**, Workflow of predicting perturbation outcomes using an optical pooled screen of cells infected with the Ebola virus and subjected to genome-wide perturbations. In step 1, a pre-trained Vision Transformer was fine-tuned on an auxiliary task to classify segmented cell images into four infection stages. In step 2, image-based features were extracted and used as inputs to MORPH, which was trained to predict imaging features after perturbations. Finally, a logistic regression model classified the predicted perturbed cell features into four infection stages. **b**, UMAP visualizations of extracted image features colored by infection state (top), or to highlight the image features extracted from *NPC1* knockout cells, a perturbation that significantly alters cell infection states, versus those extracted from cells receiving a non-targeting guide RNA (bottom). **c**, Model performance evaluation using the normalized *ℓ*_1_ loss between predicted and true infection state distribution vectors. Perturbations on the x-axis are ranked by their impact on infection states based on chi-squared test, with genes having maximal impact on the left and minimal impact on the right. The y-axis shows the cumulative mean *ℓ*_1_ loss for each method. The dotted line indicates perturbations with a Bonferroni-corrected p-value *<* 0.01. **d**, Ternary plot showing the ground-truth cell state distributions for genetic perturbations. The three axes represent the proportion of cells in the “Faint”, “Punctate”, and combined “Cytoplasmic” and “Peripheral” stages of infection. The background gradient visualizes the optimization objective, defined as the early-state fraction (the sum of “Faint” and “Punctate” states). Blue points represent the 14,973 held-out perturbations (test set). Pink points indicate the top 10 candidate genes nominated by MORPH from this held-out set.

We experimented with four types of prior knowledge on genes for the perturbation embedding **v**^*g*^: control gene expression from unperturbed cells, gene embeddings derived from the Geneformer single-cell foundation model [28], language-based embeddings obtained by GenePT [29], and embeddings derived from the DepMap database [12] containing cell viability scores across cell lines (Figure 2a). Among these priors, DepMap embeddings consistently yielded the best performance, particularly in the outlier distribution split, a trend observed across different screens (Figure 2b and Supplemental Figures 4-5). This result is notable because DepMap is the only prior based directly on experimental perturbation data, highlighting the value of using functionally grounded information to improve generalization. To confirm that informative priors are necessary for generalization, we performed ablations using non-informative perturbation embeddings (random, identity, and permuted DepMap), which markedly reduced performance under 5-fold cross-validation on K562-EG (Supplemental Figure 3a). We additionally evaluated a *prior-only* variant of MORPH in which the control-cell gene expression input was fully masked (set to zero), such that predictions relied exclusively on the DepMap embedding (MORPH (DepMap, prior-only) in Supplemental Figure 3a). Although this model underperformed the full version, the drop was smaller than when informative gene priors were replaced with non-informative embeddings. This indicates that MORPH leverages both inputs, but that performance is more sensitive to degradation of the gene prior than to masking the control-cell input. Furthermore, to ensure that performance was not driven by matched cell-line correlations between CRISPR viability screens and Perturb-seq experiments, we performed targeted ablations in which the corresponding cell-line data (K562 or RPE1 lineage variants) were removed for constructing the DepMap prior. MORPH maintained strong predictive performance on both K562-EG and RPE1-EG datasets [7] and continued to outperform the perturbed-mean baseline (Supplemental Figure 3b–c). These results indicate that MORPH learns transferable representations that generalize across cancer cell lines, rather than relying on cell-line–specific memorization.

We compared our method using the DepMap prior to a range of existing methods, including a simple baseline that assumes no perturbation effects (Control distribution), a perturbed-mean baseline that predicts the average gene expression across all perturbed training cells (Perturbed mean), as well as two state-of-the-art models: GEARS [21], a graph-based deep learning model, and a linear model [24], which we adapted to incorporate the DepMap embeddings as prior knowledge.^1^ When evaluated on the prediction of top 50 differentially expressed genes per perturbation (Fig. 2c, Supplemental Fig. 4-5), MORPH outperformed all baselines across all metrics in the K562-EG and RPE1-EG datasets, in both types of data splits (5-fold cross validation and outlier distribution split). On the K562-GW dataset, the simpler linear baseline performed competitively with MORPH. This is expected because the genome-wide screen includes many perturbations for which the top 50 DE genes show little change in expression compared to control, making near-average predictions competitive. Similarly, while MORPH is competitive on distribution-level metrics when evaluating predictions across all expressed genes for each perturbation (Supplemental Fig. 6), the simpler perturbation-mean baseline performed better for average-based metrics such as RMSE, likely reflecting that in this setting the average transcriptomic effect of perturbations is diluted by the large number of genes that are only minimally affected. These results indicate that MORPH’s improvements over baselines are most pronounced at the distribution level as well as for perturbations with strong transcriptional effects. Consistent with this observation, UMAP visualizations of the MORPH predictions for perturbations with the largest effects in the test set align closely with the observed perturbed cell states (Figure 2d and Supplemental Figure 7).

**Fig. 7.**
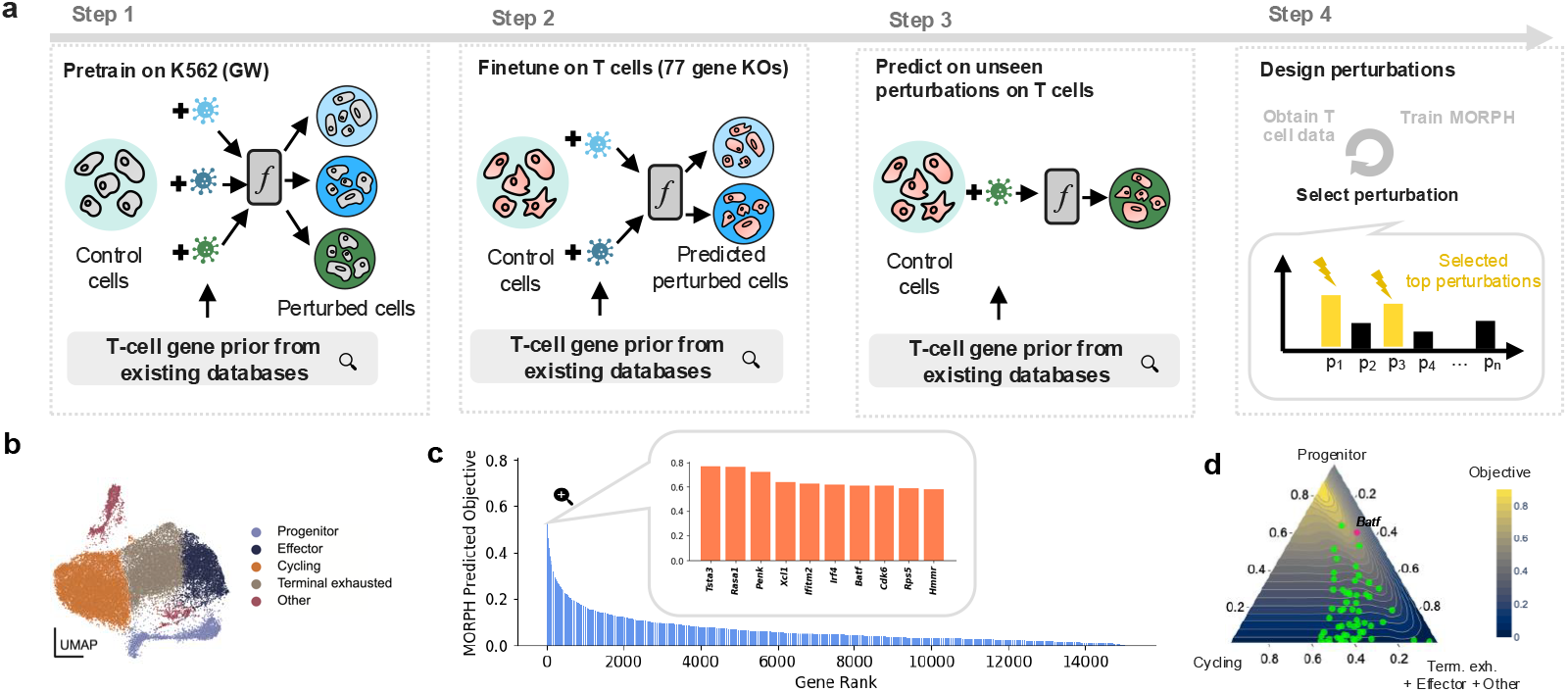
Application of MORPH to T cell differentiation. **a**, Utilizing MORPH’s capabilities to transfer across cell lines, perform perturbation prediction, and perform experiment design to identify optimal perturbations, we applied MORPH in four steps: pretraining on a genome-wide Perturbseq dataset in K562, finetuning on an in-house generated 73-gene-knockout Perturb-seq dataset in CD8 T cells, predicting the effect of 15,006 unseen perturbations in CD8 T cells, and designing perturbations with predicted improved objective function values. Across all steps, MORPH uses the same T-cell gene prior that we collected from existing datasets. **b**, Cells in the in-house generated 73-gene-knockout Perturb-seq experiment are annotated by five possible states. **c**, MORPH computes an objective function value from the predicted perturbed cell states, for each unseen perturbation. The top 10 genes predicted to achieve the highest objective function value are highlighted. **d**, MORPH identifies 1 novel gene, *Batf*, with overall better objective function value than the targets tested in the expert-curated 73-gene-knockout Perturb-seq screen (indicated by the green dots in the ternary plot).

As mentioned above, the choice of prior knowledge influenced prediction accuracy. To better understand the impact of prior knowledge on the prediction accuracy, we analyzed gene-specific performance differences (Figure 2e-f and Supplemental Figure 8). For each dataset, we identified target genes for which a specific prior outperformed others and performed gene set enrichment analysis on these subsets of perturbations (Methods). We then evaluated the performance of different priors among the significantly enriched gene sets. In the RPE1 cells, DepMap embeddings achieved the best performance across all significantly enriched gene sets (Figure 2e). GenePT language-based embeddings, particularly those incorporating information from both NCBI and UniProt, outperformed versions based solely on NCBI or STRING data (Figure 2e). In the K562 cells, DepMap embeddings also exhibited consistently strong performances, with 2 exceptions: the “ribosomal small subunit biogenesis” and the “maturation of SSU-rRNA” for which GenePT embeddings (NCBI+UniProt) led to better performance (Figure 2f). This variation may reflect context-dependent differences in how prior knowledge informs predictions. To test whether combining multiple strong priors could improve performance, we implemented a mixture-of-experts (MoE) model that integrates the language-based and DepMap priors. While the MoE model did not show substantial improvement over individual priors when evaluated on this dataset (Supplemental Table 1), it may offer more balanced performance across a broader range of genes, which could be advantageous in settings where consistency across various targets is important.

Finally, we found that MORPH remains robust to statistical perturbations of the input data. We retrained MORPH on K562-EG under gene masking (randomly zero-masking 10% or 90% of genes in control cells) and library-size rescaling (total counts of 20,000 or 100,000). Across all settings, predictive performance remained strong (Supplemental Table 2). Retrieval rank was evaluated on both the top 50 DE genes and the full gene set, as this metric is directly comparable across perturbation settings that alter expression scale. To further assess architectural robustness, we evaluated MORPH across a range of latent dimensions and decoder depths. Performance remained stable across configurations, with no single model size consistently outperforming others (Supplemental Figure 9). Together, these results indicate that MORPH is robust to relevant variations in both input preprocessing and model capacity.

In summary, we demonstrated that MORPH outperforms prior methods in predicting cellular responses to unseen single-gene perturbations, achieving robust performance across all evaluations and particularly excelling at capturing single-cell response distributions (MMD). DepMap embeddings consistently provided the most accurate predictions, particularly in challenging scenarios that require generalizing to outlier distributions.

### MORPH enables transferring perturbation effects across cell lines

With limited perturbation screens and the large number of potential cellular contexts of interest, it is crucial to determine whether perturbation effects can be transferred across distinct contexts. Specifically, we aimed to assess whether a model trained on a particular perturbation in one context—such as a particular cell line or disease state—could accurately predict the effects of the same perturbation in a different cellular context.

To evaluate MORPH’s ability to transfer perturbation effects across contexts, we trained it on RPE1-EG perturbation data and assessed its transferability to K562-EG (both datasets were obtained from Replogle et al. [7]). The model was first trained on all RPE1 perturbations, then fine-tuned using only K562 control cells by minimizing reconstruction loss (Methods). Fine-tuning on the control cells helps the model adapt to the new cell line’s basal state. For comparison, we included two baseline models: (1) the control distribution, which predicts the control cells in K562 without perturbation effects, and (2) the mean shift model, which applies the same perturbation-induced pseudobulk changes from RPE1 controls to K562 controls to predict perturbed cells in K562. Our results demonstrate that MORPH trained on RPE1 and fine-tuned on the control cells from K562 substantially outperforms both baseline models (Figure 3a). In fact, this model achieves performance comparable to MORPH trained directly on a subset of random 100 perturbations from K562. Furthermore, MORPH pretrained on RPE1 and then fine-tuned on 100 perturbations from K562 improves the performance to a level similar to MORPH trained on 1,500 perturbations from K562, suggesting that this transfer learning scheme enables MORPH to reach similar performance using only 7% (100/1,500) of the data.

By default, all model parameters were updated during this fine-tuning process. To identify which components drive successful transfer, we conducted an ablation study by fine-tuning specific modules independently. We found that fine-tuning only the decoder achieved performance comparable to updating the full model, whereas finetuning only the encoder or the gene-program matrix was insufficient (Supplemental Figure 10). This indicates that adapting the mapping from latent representations to gene expression in the target cell line (i.e., the decoder) is critical for effective transfer.

We hypothesized that perturbations transferring well across cell lines are those involving pathways or gene sets that are more similar between the cell lines. To test this hypothesis, we defined a cell line disagreement metric by calculating the Euclidean distance between the mean shift vectors for each perturbation in K562 and RPE1. We then compared the prediction performance of the model fine-tuned only on control cells with this cell line disagreement metric. We observed a positive correlation between cell line disagreement and prediction loss (Pearson correlation = 0.52, p-value *<* 0.05), indicating that perturbations affecting pathways with dissimilar effects between the two cell lines are harder to predict (Figure 3b).

By analyzing the gene sets with higher cell line disagreement, we identified pathways such as exosome and mRNA turnover, erythroid differentiation, FACT complex, 40S ribosomal subunit, cytoplasmic processes, and the mediator complex (Figure 3b). Among these, erythroid differentiation appears to be particularly cell-line-specific. K562 cells are derived from chronic myeloid leukemia [30] and retain erythroid differentiation potential [31, 32]. In contrast, RPE1 cells are non-cancerous immortalized retinal pigment epithelial cells that lack these erythroid features, likely contributing to the higher divergence in pathways related to erythroid differentiation between the two cell lines.

Pathways with higher similarity between K562 and RPE1 include ubiquitous and essential cellular processes, such as post-translational modifications and microRNA biogenesis, which regulate gene expression and protein function. Likewise, the 28S/39S ribosomal subunit and mitochondrial, critical for metabolism and protein synthesis, are more preserved across these cell lines (Figure 3b).

Overall, these analyses show that perturbation effects learned on one cell line can be transferred to a different cell line by fine-tuning the model using only control data from the new cell line. This approach outperforms baseline methods and reduces data requirements while maintaining high prediction accuracy. Transferability depends on pathway similarity between the two cell lines: conserved processes like mitochondrial functions generalize well, while cell-type-specific pathways, like erythroid differentiation, are harder to predict.

### MORPH generalizes to predict the effect of unseen combinatorial gene perturbations

With ongoing technological advances, it is plausible that experimental data for single-gene perturbations across a broad range of genes and cell lines will become increasingly accessible over time. However, generating experimental data for a comprehensive set of combinatorial perturbations will remain a significant challenge; for example, even considering perturbations involving only up to 5 genes out of around 20,000 in the human genome in one cellular context yields over 2.97 × 10^19^ combinations. Computational methods that can efficiently generate potential responses to combinatorial perturbations could offer a scalable alternative that could reduce the burden of exhaustive experimentation.

To evaluate MORPH’s ability to predict combinatorial perturbations, we investigated multi-gene perturbations from Norman et al. [5], which includes 234 single-gene and double-gene perturbations across 110,000 cells in the K562 cell line. We assessed performance under varying levels of difficulty: In the simplest case, both genes in a double perturbation had been individually perturbed and observed during training (referred to as 0/2 unseen). More challenging settings included cases where only one (1/2 unseen) or neither (2/2 unseen) of the single-gene perturbations had been seen during training.

As baselines, we compared MORPH to the control distribution, the perturbed mean, GEARS, and a family of simple additive models for double-gene perturbations. SALT [23] predicts double-gene perturbations by summing the single-gene shifts from the control mean and adding the result back to the control state, assuming additive effects, and thus requires both constituent single-gene perturbations to be observed during training. To handle settings where one or both single-gene perturbations are unseen, we used SALT+, our extension of SALT, which applies the same additive rule but falls back to the perturbed-mean baseline whenever a required single-gene profile is unavailable. Finally, we evaluated a linear-additive model, which also assumes additivity, but uses the previously described linear regression model (trained to map DepMap gene embeddings to single-gene expression profiles) to supply missing single-gene effects, enabling predictions when one or both single-gene perturbations are unseen.

Comparing against the control baseline, perturbed mean, SALT+, GEARS, and the linear-additive model, we found that MORPH consistently outperforms the current state-of-the-art approaches across all key metrics, improving performance by 33% in MMD and 22% in RMSE (Figure 3c, Supplemental Figure 5d and Supplemental Figure 11a). SALT was not included in this evaluation, since it requires that both single-gene perturbations be observed during training. Visualizing the UMAP projections of the perturbations with maximal effects in the test set, we also observed that MORPH’s predictions align more closely with the observed perturbed cells (Figure 3d and Supplemental Figure 11b).

As highlighted by Norman et al. [5], gene interaction (GI) types vary, and the effects of double-gene perturbations are not always simple additive combinations of single-gene effects. To assess how GI types influence prediction performance, we conducted a 5-fold cross-validation experiment. For each fold, MORPH was trained on all single-gene perturbations and 4/5 of the double-gene perturbations, with the remaining 1/5 of the double-gene perturbations reserved for testing. After gathering predictions across all folds, we reported the performance stratified by GI type. In this setting, all single-gene perturbations are observed during training, which implies that the additive baselines (SALT, SALT+, and the linear-additive model) become equivalent; we therefore reported SALT only. We found that the GI types, potentiation and redundancy, posed the greatest challenges for prediction across all methods (Figure 3e). Notably, current SOTA methods showed a substantial drop in performance for these GI types, while MORPH remained more robust. For instance, in redundant interactions, MORPH improved prediction performance by approximately 40% compared to the best SOTA methods.

Finally, we tested whether MORPH’s predictions could accurately classify gene pairs into their correct GI types. By comparing gene interaction scores derived from model predictions to those based on observed data (Methods and Supplemental Note 2), we found that MORPH achieved the highest Pearson correlation between predicted and true scores (Figure 3f). Additionally, when using the predictions to classify gene pairs into GI types with a threshold derived from the given labels in [5], MORPH achieved the highest area under the receiver operating characteristic curve (AUC-ROC) (Figure 3f). These results highlight the utility of our approach in not only predicting perturbation effects but also understanding and classifying the nature of complex genetic interactions.

### MORPH provides an informative gene embedding for optimal design of perturbations

Perturbation screens are costly and time-intensive. This highlights the need for strategies to optimize experimental design. We assessed MORPH’s performance on the problem of selecting perturbations to accelerate learning across the entire perturbation space. Specifically, we aimed to achieve accurate predictions across all perturbations from screening only a limited subset. Towards this, we adopted an iterative lab-in-the-loop framework [33, 34]; namely, we selected perturbations based on the score of an acquisition function computed using MORPH trained on existing data, these perturbations were then added to the dataset, and MORPH was updated using both new and existing data for the next round (Figure 4a).

The key principle of this approach is to identify the most informative perturbations that help the model generalize more efficiently. The acquisition function facilitates this selection by evaluating and ranking the unscreened perturbations based on their potential to improve model performance. We first considered two acquisition strategies commonly used in the active learning literature: prioritizing perturbations with high uncertainty [35–38] and prioritizing perturbations with dissimilar effects [39, 40]. However, these approaches performed similar to random selection (Supplemental Figure 12a-d, Methods). To overcome the limitation of these approaches that they may not capture the information in deep generative models, we adopted a learning-based approach, originally developed in the context of computer vision [41] (Methods). Specifically, to estimate the challenging perturbations for the model, we trained a light-weight auxiliary model to predict the *prediction loss* for each unscreened perturbation. We evaluated two ways to train the auxiliary model for loss prediction: (1) using representations learned by MORPH (referred to as “adaptively selected perturbations using learned representations”), and (2) using fixed, prior-knowledge representations (referred to as “adaptively selected perturbations using prior-knowledge representations”).

Using the K562-EG dataset from Replogle et al. [7], we reported the prediction loss on the held-out test set of 204 perturbations, as a measure of how well the model generalizes. The strategy of “adaptively selected perturbations using learned representations” outperformed both the random baseline and “adaptively selected perturbations using prior-knowledge representations” in test-set prediction performance, as measured by MMD (Figure 4b, Supplemental Figure 12e) and RMSE (Figure 4c, Supplemental Figure 12f). To further assess the contribution of the learned representations and model architecture, we also evaluated GEARS under both random and adaptive selection. Across both prediction metrics - MMD and RMSE (Figure 4b–c) - MORPH consistently outperformed GEARS, suggesting that MORPH more effectively captured the overall perturbation space. Overall, the performance gap between MORPH and GEARS was larger than the differences among active learning strategies within each model.

We further evaluated the selection strategies on a second downstream task: identifying perturbations with the most significant effects on cells. Since most single-gene perturbations have minimal impact on cells, identifying those with strong effects could highlight valuable experimental targets. To assess MORPH’s performance on this task, we ranked perturbations in the test set based on their predicted impact, measured by the distributional distance (MMD) between predicted perturbed and control distributions. We then calculated the precision of detecting the top 25 and 35 most impactful perturbations for each selection strategy. Also in this task, MORPH consistently outperformed GEARS, and the adaptive method using learned representations still outperformed the baseline and prior-only method (Figure 4d-e, Supplemental Figure 12g–h).

Finally, we investigated the impact of different allocations of a fixed experimental budget (here we considered 500 perturbations) on MORPH’s performance across the previous two tasks: predicting cellular responses and identifying perturbations with significant effects. Using the strategy of adaptively selecting perturbations using learned representations, we observed that allocating more perturbations to later active learning rounds generally improved predictive performance compared to front-loading them in the initial round. For instance, randomly selecting 100 perturbations in the first round (red lines) outperformed randomly selecting 300 perturbations in the first round (purple line) in terms of predictive performance measured by MMD on the test set (Figure 4f). This is likely because more strategically chosen perturbations lead to better overall performance. While going beyond two active learning rounds, performance differences among allocation strategies became negligible (different types of red lines), performing only one round without active learning (blue dot) yielded significantly worse results (Figure 4f). We observed similar trends when evaluating the model’s ability to identify high-impact perturbations: more strategically selected perturbations led to better identification performance. Having three rounds (red dotted line) outperformed two rounds (red dashed line). And five rounds (red solid line) achieved the best performance, though the difference between three and five rounds was marginal (Figure 4g). Overall, active learning with few rounds of experiments consistently enhanced the predictive performance within the fixed budget.

### MORPH’s attention-based framework allows the inference of gene regulatory programs

To enhance the interpretability and reliability of MORPH, we aimed to extract the mechanisms that it learned through training. Specifically, we sought to determine whether analyzing the weights of the different components learned by MORPH from the Perturb-seq data (Figure 1a) could reveal the underlying gene regulatory network. As in prior work [2], we modeled gene regulation via a bipartite network, where perturbations act as parent nodes and regulated genes as child nodes. Clustering the parent nodes gives rise to “perturbation modules” such that perturbations within the same module induce similar effects. Clustering the child nodes gives rise to “gene programs,” where genes within the same program exhibit similar responses to shared perturbations. The directed edges between modules (parent nodes) and programs (children nodes) represent regulatory effects, providing insights into how perturbations influence gene expression patterns.

Notably, MORPH is based on a formal causal framework: The proposed regulatory bipartite network model can be formalized as a causal structural equation model, and we proved identifiability of the causal regulatory effects in this model (Supplemental Note 4). In addition, we demonstrated on simulated data, where the bipartite network structure is known, how MORPH’s attention-based framework can recover it; more precisely, we simulated control and perturbed gene expression data based on a given bipartite network, trained MORPH on the simulated data, and analyzed the learned weights (Figure 5a; Methods). To identify perturbation modules, we clustered perturbations in the latent space generated by the perturbation encoder (Figure 1a, bottom panel of the encoder). This analysis showed that perturbations within different modules formed distinct clusters in the learned embedding space (Figure 5b). Next, we mapped gene programs to expressed genes by perturbing the attention maps to determine which genes had the highest scores within each learned program (Figure 1a, attention block and decoder; Methods). Genes corresponding to different ground-truth programs were grouped into distinct clusters (Figure 5c). Finally, we reconstructed the bipartite graph by connecting perturbation modules to gene programs using the learned attention maps (Figure 1a, attention block; Methods). This analysis showed that the reconstructed bipartite network closely resembled the data-generating network and accurately captured both positive and negative regulation (Figure 5d). These results together demonstrate that MORPH can successfully recover the true underlying regulatory network on simulated data by analyzing the weights of each component of MORPH.^2^

We then applied the same analysis to Perturb-seq data from Replogle et al. [7] and validated our findings against the structures reported in the original study. Clustering perturbations in the latent space consistently revealed perturbation modules that aligned with those identified in [7] (Figure 5e). Similarly, clustering the program-to-gene mappings uncovered gene clusters that corresponded to the provided labels in [7] (Figure 5f). Finally, we compared the model’s learned regulatory edges between modules and programs, represented as attention scores, to the bipartite structure reported in [7]. On average, the model assigned greater attention to programs that were up-regulated after perturbation when predicting perturbation outcomes, whereas down-regulated programs received less attention compared to control reconstruction. (Figure 5g). These results suggest that MORPH effectively captured biologically meaningful regulatory patterns.

As a robustness check, we retrained MORPH under gene masking in control cells and library-size rescaling and re-ran the interpretability readouts. While masking the control anchor weakened the alignment between attention differences and regulation effects defined from the unperturbed data, the inferred gene programs remained largely stable across perturbations, with a drop only under extreme (90%) masking (Supplemental Figure 14a-d). As an additional sanity check, we compared the trained model to a randomized-weight model with identical architecture. Under randomization, the structured relationship between attention and gene regulation disappeared, and alignment with annotated gene programs decreased substantially (per-latent AUROC: 0.949 vs. 0.590; Methods, Supplemental Fig. 14e), indicating that these patterns reflect learned biological structure rather than architectural artifacts. To further assess whether attention contributes beyond interpretability, we performed an explicit ablation in which the attention blocks were removed and the control-cell latent and perturbation latent embeddings were concatenated and fused using an MLP. To ensure a fair comparison, we matched model capacity by increasing the decoder depth and latent dimensionality of the non-attention model such that the total number of parameters was comparable (attention-based MORPH: 3,485,876 parameters; concatenation+MLP: 3,509,192 parameters). Using 5-fold cross-validation on three datasets (Norman K562, Replogle K562-EG, and Replogle RPE1-EG), we found that incorporating attention provides a consistent benefit, improving several key evaluation metrics across datasets while maintaining comparable performance on others (Supplemental Table 6). Together, these results indicate that the attention mechanism serves both as an interpretable structural component and as a beneficial inductive bias for predictive performance.

### MORPH is generally applicable to single-cell data including imaging-based read-outs

To evaluate the modularity of MORPH, we applied it to imaging data from an optical pooled screen [10]. To minimize batch effects, we used data from a single plate; this included 5,230,322 HeLa-TetR-Cas9 cells transduced with guide RNAs targeting 20,336 genes. These cells were infected with the Ebola virus and underwent genome-wide perturbations. Measurements were taken across six different imaging channels post-perturbation. The authors classified the cells into four infection states—”Faint”, “Punctate”, “Cytoplasmic”, and “Peripheral”—representing progression from the earliest to the latest stages of infection. The primary objective was to identify perturbations that significantly alter the distribution of infection states.

To extract features from the images, we fine-tuned a pretrained Vision Transformer (ViT) [42] (originally trained on ImageNet-21k, containing 14 million images [43]) using an auxiliary task of classifying the segmented single-cell images from the optical pooled screen into the four infection stages (Methods). The extracted image features were then used as input to MORPH (Figure 6a). To assess sensitivity to the choice of image representation, we also evaluated MORPH using unsupervised image embeddings, including DINO [44], scDINO [45], and CNN (Methods). Relative to the supervised ViT features, the unsupervised embeddings showed reduced separability of infection states and led to decreased performance for predicting perturbations with large effects (Supplemental Figure 15). By visualizing the extracted ViT features, we observed that most cells were in the cytoplasmic and peripheral states, indicative of later infection stages. Only a small subset of perturbations significantly shifted cells toward earlier infection states, such as faint or punctate (Figure 6b).

To evaluate model performance, we employed 5-fold cross-validation at the perturbation level, in which genetic perturbations were randomly partitioned into five folds. In each run, MORPH was trained on four folds and evaluated on the heldout fold, ensuring that test perturbations were not observed during training. For the held-out predictions, we classified the predicted perturbed cell features into the four infection states using logistic regression and computed the normalized L1 loss between the predicted and observed infection state distribution vectors. For comparison, we benchmarked MORPH against three baseline models: a control-distribution baseline that predicts the infection-state distribution of control cells; a perturbed-mean baseline that predicts the average distribution across perturbed cells in the training set; and a linear model that predicts infection-state distributions using prior-knowledge-based embeddings. Since GEARS cannot be applied to imaging data, we excluded it from this analysis. Both the linear model and MORPH utilized the same prior knowledge (DepMap), which we identified as the strongest prior knowledge source based on the analyses in the previous sections.

Since most perturbations did not significantly affect infection states, the control distribution baseline performed well when considering genome-wide perturbations. For evaluation purpose, we performed a chi-squared test comparing the perturbed state distribution to the control state distribution for each perturbation and ranked the perturbations based on their impact on infection states. Notably, MORPH outperformed the baselines by 33% for the top-ranked genes, i.e., perturbations that most significantly affect infection state. Furthermore, MORPH remained competitive when considering all genes (Figure 6c-d and Supplemental Figure 16).

To assess generalization to phenotypically distinct conditions, we evaluated MORPH on a challenging outlier-distribution split where perturbations causing substantial phenotypic shifts were concentrated in the test set (Supplemental Figure 17a). All models showed reduced performance in this difficult regime, while MORPH and the linear model still outperformed the control-distribution and perturbed-mean baselines (Supplemental Figure 17b). This regime naturally motivates active experimental design. As demonstrated in Figure 4d-e, MORPH can actively propose perturbations that drive the largest phenotypic shifts away from the control state, enabling targeted data acquisition to populate these outlier regions.

Finally, we evaluated whether MORPH can support a target-nomination workflow. Our goal was to identify perturbations that shift the state distribution toward earlier infection, quantified as the early-state fraction (the sum of “Faint” and “Punctate” states). To mimic a perturbation nomination experiment, we restricted the training data to a small, 3,000-perturbation expert-guided pilot screen and treated the remaining 14,973 perturbations with available DepMap embeddings as held out (Methods). By ranking the held-out perturbations using their predicted early-state fraction, we obtained the following top 10 candidates: *ARPC4, RAB7A, GET4, PMM2, ATP6V0E1, NRF1, CCDC115, WDR7, UNC50*, and *AIFM1*. We evaluated MORPH’s nominations using ground-truth measurements from the genome-wide screen. The top 10 nominated genes outperformed most genes in both the held-out test set (Fig. 6d) and the pilot set (Supplemental Figure 18), with *RAB7A* ranking 2^*nd*^ in MORPH predictions and 1^*st*^ in the test-set ground truth (across 14,973 perturbations). MORPH also identified *CCDC115* and *GET4* among the held-out genes, ranking them 7^*th*^ and 3^*rd*^ by prediction compared with ground-truth ranks of 4^*th*^ and 5^*th*^, respectively.

The original study validated some of the identified genes in a targeted secondary screen, noting that *GET4* had not previously been reported to regulate EBOV infection [10]. Together, these results demonstrate MORPH’s applicability across different data modalities and its capability to guide large-scale target discovery.

The application of MORPH to imaging data extends beyond single-gene predictions and target nomination demonstrated here. Following the same framework, MORPH can also be used for cross-context transfer and active experimental design to efficiently explore the entire morphological phenotype space. In addition, the approach naturally extends to predicting unseen double-gene perturbations to guide combinatorial imaging screens as these become available. However, interpreting the inferred regulatory network is less straightforward since the inferred graph describes a mapping from genetic perturbations to phenotypic features (rather than genes).

### MORPH identifies candidate genetic targets to mitigate T-cell exhaustion

To further demonstrate MORPH’s capacity for biological discovery, we applied it to a challenging biomedical problem, which was the core of our recent Cancer Immunotherapy Machine Learning Challenge ([46], manuscript in preparation): identifying new genetic targets to prevent T-cell exhaustion. Cytotoxic CD8+ T cells play a key role in removing abnormal/cancerous cells. However, chronic exposure to weakly immunogenic tumor-associated self-antigens along with other immunosuppressive signals progressively differentiates naive CD8+ T cells into a spectrum of exhausted T cell states. Transiently increasing the proportion of progenitor-like cells has been shown to improve outcomes under immune checkpoint blockade [47]. Our goal was to predict gene perturbation targets that could shift T cell states to increase the progenitor-like cell ratio. To this end, we synergistically employed three of MORPH’s capabilities: transfer perturbation effects across cell lines, perform perturbation prediction, and perform experiment design to identify optimal perturbations (Figure 7a).

First, we pretrained MORPH using as input the K562-GW dataset from [7], and a T-cell gene prior that we assembled by extracting data from prior literature. This included data on perturbations in T cells from proliferation [48–50], production [51– 53], and bulk RNA-seq screens [54], but no Perturb-seq data (Methods). Next, we finetuned MORPH to predict perturbation effects using the in-house generated single-cell Perturb-seq dataset on CD8 T cells generated for our Challenge. This dataset measures the transcriptional effect of 73 gene knockouts on T cells *in vivo* after injection into a mouse harboring melanoma for 14 days. By annotating each single cell as one of five T cell states: “Progenitor”, “Effector”, “Cycling”, “Terminal exhausted”, and “Other” (Figure 7b), we summarized the Perturb-seq data for each of the 73 perturbed genes by a five-dimensional proportion vector of perturbed cells in each state. We then utilized the fine-tuned MORPH model to predict unseen perturbation effects. For all 15,006 genes with expression measured in the 73-gene-knockout Perturb-seq data but not perturbed among the 73 genes, we generated predicted perturbed cells under the corresponding perturbations. Each predicted perturbed cell was mapped to one of the five T cell states using an MLP classifier trained on the 73-gene-knockout data, yielding a predicted cell state proportion for each untested gene. Finally, we designed targets with potentially improved objectives by ranking the untested genes. The objective function was designed to capture the increased proportion of progenitor cells. Specifically, the objective function for a given perturbation was the fraction of progenitor cells it produces, provided that the proportion of cycling cells was predicted to exceed 5%; otherwise, the objective was set to zero. The constraint on cycling cells ensured that the perturbed cells retained differentiation capabilities.

Applying this four-step approach, we nominated untested targets with potentially improved objectives compared to the 73 tested genes. This task was challenging because the 73 genes in our Perturb-seq panel were expert-curated based on domain knowledge of genes that strongly affect T cell differentiation, representing the best-known targets to date. As shown in Figure 7c, the top 10 nominated genes (out of 15,006) were: *Tsta3* (0.7688), *Rasa1* (0.7656), *Penk* (0.7250), *Xcl1* (0.6406), *Ifitm2* (0.6281), *Irf4* (0.6188), *Batf* (0.6094), *Cdk6* (0.6094), *Rps5* (0.5875), and *Hmmr* (0.5813). Notably, *Batf* (ranked 7/15,006 by MORPH) is a transcription factor that was validated by an independent Perturb-seq screen on CD8 T cells [55]. By annotating cells using the same cell state definitions as in the 73-Perturb-seq dataset used for training, we obtained scores for all genes in the independent screen. In this independent screen, *Batf* achieved higher objective values (0.6061) than 72 of the 73 genes in the 73-Perturb-seq dataset used for training (top two scores: 0.6429 and 0.6) (Figure 7d). Importantly, the independent screen was not used during MORPH training, which relied solely on our T-cell gene prior, the K562-GW dataset, and the in-house generated 73-Perturb-seq dataset. Therefore, the identification of *Batf* provides independent validation of MORPH’s ability to nominate targets with the potential to improve cancer immunotherapy.

## 3 Discussion

In this study, we presented MORPH, a deep-learning framework designed to predict the single-cell distribution of transcriptional or phenotypic outcomes resulting from unseen genetic perturbations. By combining prior biological knowledge on genetic perturbations with transcriptomic or phenotypic information from control cells, MORPH effectively extrapolates perturbation effects beyond observed data in both sequencing and imaging modalities. Concurrently and independently, TxPert [25] advanced the state of the art for transcriptomic perturbation prediction by leveraging multiple biological knowledge graphs, demonstrating that cross-context transfer is achievable for sequencing-based data. Because this work was developed in parallel with ours, a direct benchmarking comparison was not included; MORPH’s contributions are best understood as complementary: where TxPert focuses on transcriptomic prediction accuracy, MORPH uniquely addresses cross-modality prediction—extending to imaging phenotypes—and the interpretability of learned gene relationships, enabling inference of the underlying gene regulatory structure from perturbation responses. Through simulation and empirical validation, we found that MORPH learns perturbation-response patterns by attending to groups of genes that are regulated similarly in response to a perturbation. This capacity enables MORPH to extrapolate perturbation responses to unseen single or combinatorial perturbations, leveraging similarities with perturbations seen in training or captured in prior biological knowledge. Furthermore, MORPH can transfer the learned perturbation-response patterns to unseen cell lines using only control cell data in the new context. We demonstrated that the gene relationships captured by MORPH’s attention framework are biologically meaningful and offer insights into the cell line-specific underlying gene regulatory networks—a feature not available in previous methods for predicting perturbation effects.

MORPH’s modular design allows for easy integration of different prior knowledge sources, making the framework highly adaptable and extensible. Among all sources of prior knowledge, we identified DepMap [12] as the most informative database on the perturbation screens considered in this paper, possibly because it is the only database built directly from perturbation experiments supporting the notion that perturbational relevance matters more than data quantity — a finding that is consistent with results from the first Competition within our Cell Perturbation Prediction Challenge [19], where features derived from perturbational data systematically outperformed those from observational databases. As an example of adapting MORPH using a context-relevant dataset, in our T cell application, MORPH pretrained on the human K562 genome-wide perturbation data was fine-tuned on an in-house mouse CD8 T-cell Perturb-seq dataset of 73 selected perturbations. Despite the limited amount of context-relevant perturbation data, MORPH was able to transfer the perturbation-response structure learned on K562 cells to mouse T cells and effectively predict the effects of unseen perturbations, which we then ranked using an objective function to quantify the increased ratio of progenitor cells. A target identified by MORPH was able to shift T-cell states towards the desired ratio more effectively than majority of the expert-designed gene perturbation panel, as confirmed on an independent dataset.

In a separate application on OPS data, we demonstrated MORPH’s cross-modality robustness. When trained on a fraction of the available OPS data, MORPH was able to effectively predict genes that could impair the Ebola virus infection progression in HeLa cells, as confirmed using the held-out data. Another key advantage of MORPH over previous approaches is its ability to predict perturbation effects at the distributional level, rather than only estimating average responses. This enables the model to capture heterogeneous cellular responses that are often obscured by mean-based predictions. Notably, MORPH shows the largest performance gains for genes with the strongest perturbation effects. As a result, MORPH predictions are particularly well suited to identify perturbations capable of driving cells away from baseline states. This ability can be leveraged to efficiently guide the targeted acquisition of context-relevant perturbational data to rapidly expand coverage of the perturbation space. At the same time, MORPH further benefits from carefully selected additional perturbations, with performance improving at a faster rate than the state-of-the-art method GEARS [21]. In summary, these two applications (T-cell and OPS) demonstrate how MORPH’s capacities can be used synergistically to identify targets predicted to move the cells to a desired state, supporting both biological discovery and potential target identification.

Looking ahead, MORPH is expected to improve as more perturbation screening data becomes available. Its iterative framework could be used to efficiently generalize to new cellular contexts by leveraging the perturbation embeddings learned in one context to iteratively identify and prioritize the most informative perturbations in another context. Furthermore, MORPH could directly be extended beyond genetic perturbations to model the effects of chemical perturbations, enabling applications in drug discovery [56]. Overall, MORPH provides a versatile framework for modeling perturbation effects across different modalities, with broad applications to interrogating biological mechanisms and aiding in the development of more precise and effective treatments.

## 4 Resource availability

### Lead contact

Further information and requests for resources should be directed to and will be fulfilled by the lead contact, Caroline Uhler.

### Data and code availability

#### Data

The datasets analyzed in this paper are publicly available and include Perturbseq datasets from Norman et al. [5], Replogle et al. [7], and Dixit et al. [2], as well as the optical pooled screen dataset from Carlson et al. [10]. In addition, we used gene embeddings derived from Geneformer [28], GenePT [29], and DepMap [12]. The corresponding accession numbers of the single-cell datasets are listed in the attached Key Resources Table. A processed version of the Norman [5] and DepMap [12] datasets can be retrieved from: https://drive.google.com/drive/folders/1TQJE281q4xH7HcNHMg1v0urD99EDj5bO?usp=drive_link.

#### Code

All original code is publicly available and has been deposited at https://github.com/uhlerlab/MORPH. The repository also contains the open-source software packages used, a detailed description on how to reproduce all our results, as well as a demo application and instructions on how to apply our pipeline to user-provided datasets.

## 5 Acknowledgments

We would like to thank E. Forte for valuable discussions and feedback on the manuscript, and J. Weissman for helpful discussions. C.H, J.Z. and M.D. were supported by the Eric and Wendy Schmidt Center at the Broad Institute. J.Z. was partially supported by an Apple AI/ML PhD Fellowship. C.U. was partially supported by NCCIH/NIH (1DP2AT012345), NIDDK/NIH (5RC2DK135492-02), ONR (N00014-24-1-2687), AstraZeneca, the United States Department of Energy (DE-SC0023187), and MIT J-Clinic for Machine Learning and Health.

## 6 Author contributions

All authors designed the research. C.H. and J.Z. developed and implemented the algorithms. C.H., J.Z., performed model and data analysis. M.D. and C.U. supervised the study. All authors wrote the paper.

## 9 Methods

### Learning framework of MORPH

Single-cell RNA sequencing and other sequencing-based assays are inherently destructive, where each cell is usually only measured either before or after a perturbation, but not both. Consequently, we only have access to distribution-level data rather than paired individual measurements from before and after perturbation. Thus, MORPH considers a setting where unpaired samples are drawn from control and perturbed distributions: ℙ(*X*^∅^) and ℙ(*X*^*g*^), where ℙ(*X*^∅^) represents the control distribution and ℙ(*X*^*g*^) represents the distribution under perturbation on gene *g*. Each sample *X*^·^ ∈ ℝ^*D*^ from these distributions represents features of a cell, such as its gene expression levels across *D* genes or its image features of dimension *D*.

Given a control sample *X*^∅^ and a perturbation embedding 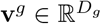 for a gene *g* (see the following section for details on how to obtain the perturbation embedding), the goal is to learn a function

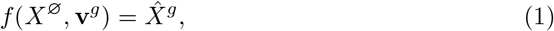

that predicts the perturbed outcome 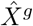, which can be applied to both “seen” gene perturbations (i.e., perturbations that have been tested and are in the dataset) and “unseen” gene perturbations. Building on [26], we learn the function *f* via a discrepancy-based variational autoencoder (VAE), which minimizes a distributional loss between the predicted perturbed samples 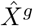 and the real perturbed samples *X*^*g*^.

To be more precise, in this framework, the control distribution P(*X*^∅^) is modeled using a VAE. Let *Z* denote the latent variables that generate the cell features *X*^∅^ and p(*Z*) be their prior distribution, realized using a standard Gaussian. The VAE consists of two components: an encoder q_*ϕ*_(*Z* | *X*^∅^), parameterized by *ϕ*, which approximates the posterior distribution of the latent variables given *X*^∅^; and a decoder p_*θ*_(*X*^∅^ | *Z*), parameterized by *θ*, which approximates the conditional distribution of *X*^∅^ given *Z* under no perturbation. The Evidence Lower Bound (ELBO) for the likelihood of observing a sample *X*^∅^ is:

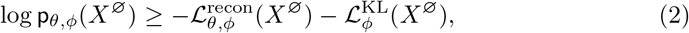

where

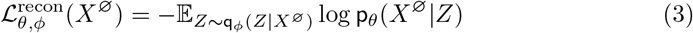

and

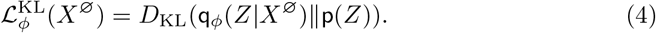

Here, the encoder q_*ϕ*_(*Z* | *X*^∅^) is realized via a Gaussian distribution, where the mean and standard deviation are parameterized by *ϕ*. The sampling of *Z* from q_*ϕ*_(*Z* | *X*^∅^) is implemented via the reparamterization trick for back-propagation [57]. The term *D*_KL_(·∥·) represents the Kullback–Leibler (KL) divergence, which regularizes the latent space by encouraging the marginal approximate posterior q_*ϕ*_(*Z X*^∅^) to be close to the prior p(*Z*). We use a deterministic decoder, where denoting the output as 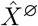 after decoding *Z* sampled from q_*ϕ*_(*Z* | *X*^∅^), and it reduces to an *l*_2_ loss for sample reconstruction of the control cells:

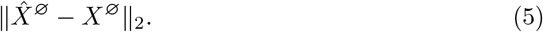

To model the perturbation effects, MORPH uses a discrepancy-based component. Specifically, MORPH encodes *X*^∅^ into *Z* and **v**^*g*^ into a latent representation, and concatenates them into a joint latent representation *Z*. The decoder then generates samples 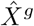 using *Z*^*g*^, simulating the effect of perturbing gene *g* on the control cells *X*^∅^. We measure the discrepency between the distribution of generated perturbed samples 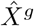 and real perturbed samples *X*^*g*^ using maximum mean discrepancy (MMD) [58], which is defined as

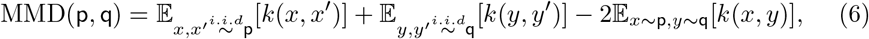

for two arbitrary distributions p and q, where *k* denotes a kernel function. Denoting by *G* the set of all genes that were pertrubed in a dataset, MORPH uses the following discrepency loss:

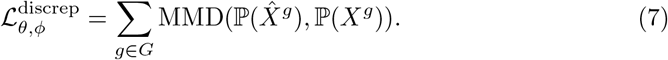

We also calculated an MMD loss for the control cell distributions, since we observed that this made attention score shifts more reflective of the perturbation effects magnitude (Supplemental Figure 1). Therefore, the total loss for reconstructing the control cells is:

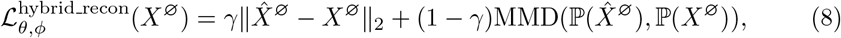

where *γ* is a tunable hyperparameter.

The full objective function minimized by MORPH for model training is:

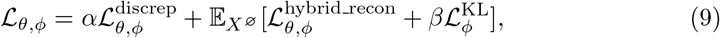

where *α, β, γ* are hyperparameters controlling the balance among alignment between predicted and observed perturbation distributions, regularization of the latent space, and accurate reconstruction of control states. The selection of hyparameters is discussed in Supplemental Note 2.

### Incorporating biological knowledge to represent perturbations

To generalize to unseen perturbations, MORPH represents each gene perturbation using a feature vector, i.e., 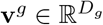, which represents prior knowledge of gene *g* and its perturbation effect. The feature vector allows MORPH to relate unseen perturbations to previously seen ones, based on the intuition that perturbations with similar biological properties tend to have similar effects. We explored multiple sources of prior knowledge to construct these vectors.

#### Control gene expression

The simplest form of prior knowledge leverages gene expression profiles from control cells. Let 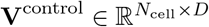 denote the gene expression matrix of control cells, where *N*_cell_ denotes the number of control cells and *D* is the number of genes measured in each cell as defined in the previous section. To obtain a feature vector for perturbing gene *g*, the *g*^*th*^ column of **V**^control^ is extracted, i.e., 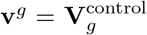. This approach is based on the hypothesis that genes with similar expression patterns in the control cell may exhibit similar effects when being perturbed. In our experiments, we randomly sampled *N*_cell_ = 1, 500 from the control cells to obtain a 1, 500-dimensional feature vector for each gene.

#### Geneformer

The emergence of single-cell foundation models derived from large-scale single-cell datasets provide new approaches to obtain gene feature vectors. Specifically, we utilized the gene embeddings from Geneformer [28], which offer the advantage of context-dependent representations. In particular, we fed control cell data into the pretrained foundation model and extracted the second-to-last layer’s^3^ output for each gene per control cell. We then averaged these vectors across all control cells to obtain a 256-dimensional feature vector for each gene.

#### GenePT and GPT

As proposed in [29], we defined another prior by leveraging OpenAI’s ChatGPT text embedding [59]. We obtained the gene embeddings shared by the authors, which were derived from gene summaries in NCBI [60] and UniProt [61]. Beyond these sources, we also explored generating embeddings using the same approach but incorporating information from STRING [62]. This resulted in three different text-based priors, namely, a 1, 536-dimensional gene feature for NCBI, a 3, 072-dimensional gene feature for NCBI and UniProt, and a 1, 536-dimensional gene feature for STRING.

#### DepMap

Finally, we explored gene embeddings derived from the DepMap database. DepMap provides a matrix 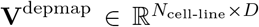, where *N*_cell-line_ = 1, 100 is the number of cell lines in the dataset. Each entry 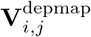 represents the viability of cell line *i* under perturbation *j*. To represent a specific perturbation on gene *g*, we extracted the *g*^*th*^ column of the matrix **V**^depmap^, i.e., 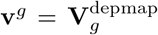, resulting in a 1, 100-dimensional feature vector **v**^*g*^.

### MORPH’s attention mechanism

To refine the joint latent embedding *Z*^*g*^ of a control cell and a gene feature vector, MORPH employs a cross-attention mechanism [63] in the latent space. Toward this, the *D*_*Z*_-dimensional joint embedding 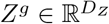 serves as the query, and to construct keys and values, MORPH learns a latent gene program matrix 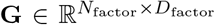, where we chose *N*_factor_ = 50^4^ gene factors each of dimension *D*_factor_ = 50. The attention mechanism integrates information from the matrix **G** into *Z*^*g*^ dynamically, allowing the model to capture complex dependencies between perturbations and gene programs. The model has 2 attention layers, with the output 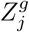 of the *j*^*th*^ layer for *j* ∈ {1, 2} computed using:

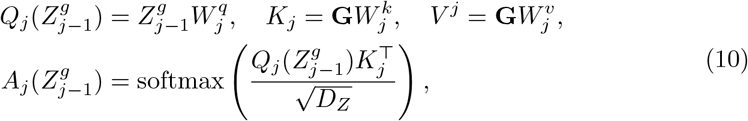

where 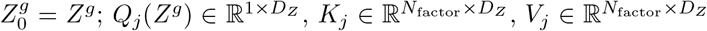 represent the query, key and value matrices; 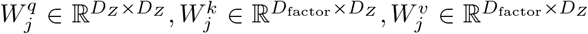 are the learnable weight matrices.

The attended output 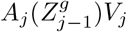 is added to 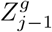 using a residual connection:

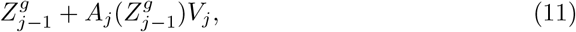

and is then passed into an MLP to obtain 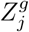, similar to a standard transformer block; see e.g. [63]. Finally, the refined latent representation 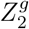 after 2 attention layers is decoded using an MLP to predict the post-perturbational cellular response 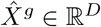.

### Transfer of perturbation effects across cell lines

To transfer perturbation effects from cell line 1 to cell line 2, we first trained MORPH on all perturbation data from cell line 1. This yielded a mapping *f*_1_, which is trained on and models the perturbation response in cell line 1.

Next, we fine-tuned *f*_1_ by minimizing the reconstruction loss in equation (9) using only the control cells from cell line 2. This step ensures that the model aligns its latent space to the new cell line’s distribution. Finally, we used the finetuned model, denoted by *f*_1→2_, to predict the effect of perturbing gene *g* in cell line 2; i.e., 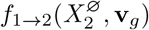, where 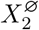 denotes the control cell from cell line 2 and **v**_*g*_ is the feature vector for gene *g*.

In the evaluations, we also considered fine-tuning *f*_1_ using a subset of perturbations from cell line 2. This alters the procedure described above by obtaining *f*_1→2_ using additionally this subset of perturbations by minimizing the full objective in equation (9).

### Genetic interaction prediction

For model evaluation, we performed 5-fold cross-validation by splitting the double-gene perturbations into five equal folds. In each iteration, the model was trained on all single-gene perturbations and four of the five double-gene perturbation folds, while the remaining fold was used for testing. This process was repeated across all five folds. The same procedure was also applied to evaluate the baseline models.

After training, we computed gene interaction scores for the test set perturbations following the approach described in [5] and then classified each gene pair into a specific GI subtype (Supplemental Note 2). This was performed on the predicted and true observed gene expression data. Model performance was assessed using two metrics: (1) Pearson correlation between predicted and observed GI scores, and (2) the area under the receiver operating characteristic curve (AUC-ROC) for classifying gene pairs into GI subtypes.

### Adaptive design of perturbation experiments

To investigate whether MORPH can help with designing an efficient strategy for adaptively selecting perturbations, we considered the problem of minimizing the model’s prediction error on unseen perturbations given a fixed budged of perturbations that could be performed. Towards this, we adapted three prominent experimental design strategies described below and compared these to a random baseline, which consists of choosing random perturbations in every experimental round.

#### Selection strategy based on prediction error maximization [41]

The active learning process iteratively updates the learned perturbation prediction model over *R* rounds. In round 1, a random subset of genes is selected for which perturbation experiments are performed. These experiments, together with control cells, are used as warm-up to obtain *f* ^(1)^. In round *i*+1, denote the perturbations performed so far as 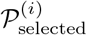 and the current model as *f* ^(*i*)^. The perturbations 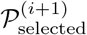 in the next batch of experiments are chosen, which maximize the prediction error; prediction error is estimated as follows, adapted from [41].

Let 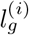 be the ground-truth prediction error of the current model *f* ^(*i*)^ on gene 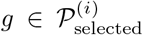, evaluated using the MMD between the observed and predicted perturbed cells. In order to extrapolate the prediction error to unseen genes, a light-weight prediction model *h*^(*i*)^ is trained to minimize the mean squared error (MSE) loss:

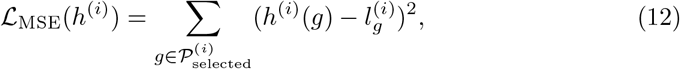

as described in the following paragraph. All perturbations are ranked using *h*^(*i*)^ and the perturbations with highest prediction error as measured by *h*^(*i*)^ are selected for the next batch of experiments 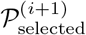. The model is then updated from *f* ^(*i*)^ to *f* ^(*i*+1)^ using the additional perturbation data.

We considered two approaches for obtaining *h*^(*i*)^: (1) using the learned perturbation representation, which is obtained by passing **v**^*g*^ through the perturbation encoder of *f* ^(*i*)^, as input to *h*^(*i*)^, and (2) using the prior-knowledge perturbation representation, i.e., **v**^*g*^, as input to *h*^(*i*)^.^5^ This comparison allowed us to evaluate whether incorporating a dynamically learned latent space improves the selection of perturbations over a static prior-based representation.

#### Selection strategy based on coverage maximization [39, 40]

We also explored a coverage-based strategy [39, 40] (Supplemental Figure 12). To estimate coverage, we applied *k*-means clustering to the learned perturbation representation, which is obtained by passing **v**^*g*^ through the perturbation encoder of *f* ^(*i*)^, at each round *i*, for all genes *g*. The number of clusters was set equal to the number of perturbations we can select in round *i* + 1, and the perturbations closest to the cluster centers were selected for the next batch of experiments 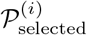. Similarly, we also benchmarked this strategy by applying the same *k*-means clustering method directly to the prior-knowledge perturbation representation **v**^*g*^, which remains fixed during training.

#### Selection strategy based on uncertainty maximization

The last strategy we explored involves selecting the most uncertain samples by estimating uncertainty within the learned perturbation representation, which is obtained by passing **v**^*g*^ through the perturbation encoder of *f* ^(*i*)^ (Supplemental Figure 12). To assess uncertainty, we measured the sensitivity of the latent representation to perturbation inputs. Specifically, for each *g*, we quantified uncertainty as the magnitude of the rate of change in the latent space, evaluated by the difference of the encoded perturbation representation at two consecutive training epochs. To obtain a scalar measure, we computed its *l*_2_-norm and averaged across epochs. This approach is based on the intuition that if the model has effectively learned the effect of a perturbation, its corresponding latent representation should remain relatively stable.

#### Evaluation

To assess performance of each approach, we considered two types of evaluations. First, we evaluated the model prediction on a held-out set of perturbations, *P*_test_, across all rounds using MMD and RMSE between the predicted and observed perturbed cell distributions; see the evaluation section below for more details.

Second, we considered the downstream task of identifying perturbations with the most significant effect. Let *Ŝ*^*N*^ be the set of perturbations predicted to have the top *N* largest MMD distance from the control distribution and *S*^*N*^ be the corresponding set based on the observed data. We compared these sets using

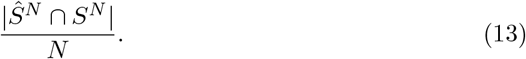

### Data simulation for gene regulatory network inference

To generate simulated data based on a bipartite structure, we first constructed a colored directed bipartite graph *G*, as shown in Figure 5a. This graph consists of two disjoint sets of nodes: parent nodes, partitioned into clusters *M*_1_, *M*_2_, *M*_3_, and child nodes, partitioned into clusters *P*_1_, *P*_2_, *P*_3_. Regulatory relationships between parent and child nodes are represented by colored directed edges, where blue edges denote activation effects and red edges denote inhibitory effects. If a cluster *M*_*k*_ is connected to a cluster *P*_*ℓ*_, then every node in *M*_*k*_ regulates every node in *P*_*ℓ*_.

We obtained the regulation matrix 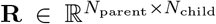, where each entry **R**_*i,j*_ represents the regulatory effect of parent node *i* on child node *j*, as follows:

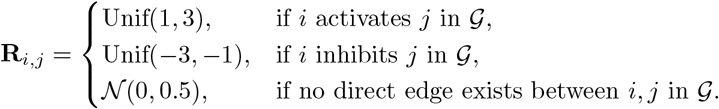

Let **X**^parent^ denote the vector of parent nodes. The value of each parent node *i* was sampled independently from a normal distribution:

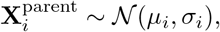

where *µ*_*i*_ ~ Unif(0, 3) and *σ*_*i*_ ~ Unif(0, 1). The values of the child nodes were computed as a weighted sum of the parent nodes:

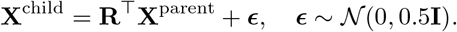

To simulate single-gene perturbations, we applied soft interventions on individual parent nodes as follows. For each intervention *I* applied to a parent node *i*, the value was modified as:

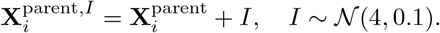

The child node values were then recalculated based on the perturbed parent nodes:

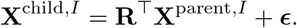

We sampled 6, 500 cells, trained MORPH on this simulated data, and evaluated the inferred bipartite regulatory network by comparing it to the ground-truth network G.

### Interpreting model predictions through gene programs

To identify the gene programs, we manipulated the attention scores and observed their resulting changes in the output space. Let 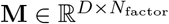 denote a mapping matrix, where each entry **M**_*ij*_ quantifies the attribution of gene *i* to factor *j*. **M** is obtained as follows and is used to estimate the gene programs from the learned gene program matrix **G**.

We manipulated the first attention map 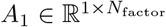 defined in Equation (10) by setting the attention weight corresponding to factor *j* in **G** to 1 and all other attention weights to 0. This operation simulates a scenario where the model focuses entirely on factor *j* when generating the predicted gene expression profile. We set column *j* in **M** to the model’s predicted output under this manipulation. To obtain the gene programs, we performed hierarchical clustering on the rows of **M**.

To identify the bipartite regulatory network connecting perturbation modules and gene programs, we leveraged the learned attention weights. As before, we used the first attention layer *A*_1_. For each control cell *X*^∅^ and perturbation *g*, the trained model produces an attention map *A*_1_(*Z*^*g*^) that guides the prediction of a perturbed cell, and a control attention map *A*_1_(*Z*^∅^) that guides the prediction of a control cell. Averaging across *N*_*g*_ samples yields the mean attention map 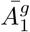 for a perturbation *g*, where *N*_*g*_ denotes the number of cells with perturbation *g*. To emphasize relative reweighting across factors, the mean attention scores were rank-normalized across factors within each perturbation. We then quantified the effect of the perturbation by computing the difference between rank-normalized attention scores: 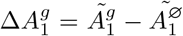, where 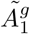 denotes the rank-normalized attention profile. This map reveals how the model shifts its focus across gene programs when predicting the perturbed state. To propagate perturbation effects from gene programs to downstream genes, we multiplied the perturbation-induced attention shift with the program–to-gene mapping matrix. Specifically, for a perturbation *g*, we computed 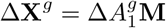, where 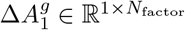 captures the perturbation-induced change in factor attention and 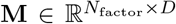 maps factors to gene expression outputs. The resulting vector Δ**X**^*g*^ ∈ ℝ^1×*D*^ quantifies predicted up- and down-regulation of individual genes induced by perturbation *g*. To obtain module-to-program regulation, we first averaged 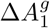 across perturbations within the same module to compute 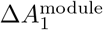, propagated this through the factor–gene mapping via 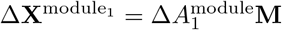, and finally averaged the gene-level scores across genes belonging to each program.

#### Quantification of gene program alignment

To quantify alignment between latent gene programs and annotated gene sets, we computed the area under the receiver operating characteristic (AUROC) scores. For each latent program *j*, we used the corresponding column of the mapping matrix **M**, where **M**_*ij*_ denotes the attribution score of gene *i* to latent program *j*, as continuous gene-level scores. For each pair of latent program and annotated gene set, AUROC was calculated by treating these attribution scores as predictors and membership in the annotated set as binary ground-truth labels.

Because the latent programs are learned in an unsupervised manner and have no predefined correspondence to annotated programs, we adopted a best-match strategy. Each latent program was assigned to the annotated gene set with which it achieved the highest AUROC. Averaging these best-match AUROC values across latent programs yielded the per-latent AUROC. Conversely, each annotated gene set was assigned to the latent program achieving the highest AUROC. Averaging these values across annotated gene sets yielded the per-program AUROC.

### Application to imaging data

To demonstrate the applicability of MORPH on imaging data, we applied it to predict the infection state distribution of Ebola-virus-infected cells under different perturbations based on an optical pooled screening dataset [10].

To extract image features that are then inputted into MORPH, we selected the three most relevant imaging channels: DAPI, VP35 RNA, and VP35 protein. We inputted these channels into a Vision Transformer (ViT) [42], which was pretrained on ImageNet-21k with 14 million images. We finetuned the resulting embeddings using an auxiliary classification task, where the images of control cells were classified into four infection states—”Faint”, “Punctate”, “Cytoplasmic”, and “Peripheral”—as labeled in [10]. This auxiliary task was trained by minimizing the following supervised contrastive loss [64]:

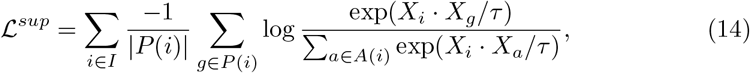

where *I* is the collection of indices *i* for all images in the batch, *P* (*i*) is the set of indices of images that belong to the same infection state as the *i*-th image, *A*(*i*) = *I \* {*i*} is the set of indices of all images in the batch except the *i*-th image, *τ >* 0 is a temperature parameter, and *X*_*i*_ is the feature representation of image *i*, which serves as input to MORPH.

To assess the impact of the representation on downstream perturbation prediction, we additionally evaluated alternative feature extractors based on fully unsupervised learning. Specifically, we considered representations from DINO [44], scDINO [45], and a convolutional autoencoder (CNN). For DINO and scDINO, pretrained models were applied directly to the optical pooled screening images to extract fixed feature embeddings. For the CNN baseline, we trained a convolutional autoencoder to reconstruct the input images and used the latent representations from the encoder as features. For consistency across methods, all representations were computed using the same three imaging channels (DAPI, VP35 RNA, and VP35 Protein). These features were used as inputs to MORPH.

To evaluate the capability of MORPH to predict infection states under unseen perturbations, we fit a logistic regression on the extracted image features of the training cells to classify them into the four infection states, and we applied this logistic regression model to the image features predicted by MORPH for unseen perturbations. Since most perturbations resulted in a cell state distribution nearly identical to the control distribution, we applied a chi-squared test, as described in the following paragraph, on the predicted state distribution to de-noise the predictions: if the test statistic was below a threshold, the prediction was set to the control state distribution vector. These de-noised predicted state distribution vectors were then compared with the ground-truth state distribution vector.

For each perturbation, the chi-squared test compared its (predicted) state distribution to that of the control cells. Given four possible infection states—”Faint”, “Punctate”, “Cytoplasmic”, and “Peripheral”—the chi-squared statistic was computed with three degrees of freedom as: 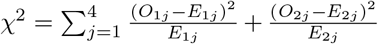 where *O*_*ij*_ represents the count of cells in state *j* for control (*i* = 1) and perturbation (*i* = 2). The expected counts, *E*_*ij*_, were calculated as 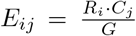, where 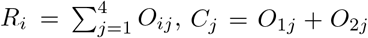 *C*_*j*_ = *O*_1*j*_ + *O*_2*j*_, and *G* = *R*_1_ + *R*_2_. To account for multiple hypothesis testing, we applied a Bonferroni correction to adjust the resulting p-values.

#### Simulation of target nomination

To evaluate MORPH in a perturbation nomination setting, we implemented the following steps. We assembled a training set starting with 18 known Ebola regulators, including *NPC1, DHODH, CAD, UMPS, VPS11, VPS16, VPS18, VPS33A, VPS39, VPS41, CTSB, CTSL, SPNS1, GNPTAB, UVRAG, PIKFYVE, FIG4*, and *EXT1* [10]. We then supplemented these with 1,491 perturbations randomly sampled from the top 30% of all 20,336 genome-wide perturbations ranked by early-state fraction (the sum of “Faint” and “Punctate”), and 1,491 perturbations randomly sampled from the remaining pool. This resulted in a training set of 3,000 perturbations in total. We then trained a Vision Transformer (ViT) on the pilot screen and used it to extract image embeddings, which were then used to train MORPH following the same pipeline as previously described. For the remaining 14,973 held-out genes with available DepMap embeddings, we used MORPH to generate predicted perturbed cell features. We mapped each predicted cell to one of the four infection states using a softmax regression model trained on the pilot data, yielding a predicted cell state proportion for each heldout gene. Finally, we ranked the held-out perturbations by their predicted early-state fraction to identify the top candidates for validation against the ground truth.

### Application to T cell differentiation

For pretraining, we used the K562-GW dataset [7], containing 9,823 perturbations measuring the expression of 8,248 genes across over 1,900,000 cells (after quality control). To ensure that the pretrained model can be transferred to the target cell line (i.e., the in-house 73-gene-knockout Perturb-seq data in CD8 T cells in mouse), we subsampled the measured genes to the top 5000 highly variable genes that are present in both datasets after standard library size normalization and log transformation. Note that K562 is a human cell line, whereas the target cell line is in the mouse; therefore, a gene is considered shared if its human ortholog is measured in K562 and its mouse ortholog is measured in the target cell line. After preprocessing the datasets, we pretrained MORPH using the same hyperparameters and setup as used for perturbation prediction on K562 (Supplemental Note 1), except that the held-out test set is set to none to ensure all perturbations are used either in the training set or in the validation set. After pretraining, we finetuned MORPH on the target cell line using the same setup as in the transferring across cell line experiments, using randomly selected 85% of the 73 perturbations for training and the remaining 15% for validation.

The T-cell gene prior used in both the pretraining and finetuning steps is constructed by concatenating after minimax normalization each individual feature from various sources: proliferation scores under various conditions [48–50], production scores of various markers [51–53], and time-coarse bulk differential expression [54].

The independent Perturb-seq dataset from [55] measures the effect of 180 transcription factor knockouts, amongst which 30 are overlapping with the in-house generated 73-gene-knockout Perturb-seq dataset. To compute the objective function on this dataset, we first used Harmony [65] to remove batch effects between this dataset and the in-house generated dataset, we then assigned each cell in this dataset to the majority cell state of its neighboring cells in our annotated in-house generated dataset. The neighboring cells were defined using Leiden clustering, where a cell’s neighboring cells are defined to be the other cells in its Leiden cluster. Cells with no neighboring cells in our in-house generated dataset were excluded from the analysis.

### Datasets and preprocessing

#### Single-cell RNA sequencing data

For the Replogle [7], Dixit [2] and in-house Perturb-seq datasets, the Scanpy toolbox [66] was used to perform cell and gene filtering, library size normalization, and log1p transformation. For all datasets, the 5,000 most highly variable genes, identified using the `highly_variable_genes` function [66], were selected to reduce the complexity of the prediction problem, following preprocessing approaches similar to [21, 67]. For cell-line transfer experiments, datasets were first restricted to the intersection of commonly measured genes across cell lines, after which the top 5,000 highly variable genes were selected using the source dataset together with control cells from the target dataset. Preprocessing for the Norman [5] dataset was inherited from [67]. To ensure sufficient data for learning a distribution, perturbations with fewer than 16 cells were filtered out from the datasets. Perturbations with between 16 and 32 cells were upsampled (with replacement) to 32 cells.

#### Optical pooled screening data

The processed imaging data, single-cell masks, and metadata were obtained from the Google Cloud Storage provided by [10]. For each train-test split, the ViT model was fine-tuned on the training set using an auxiliary task of classifying images into different infection states, and the fine-tuned ViT model was then applied to the entire dataset to extract image features; see the previous section for more details. Since the dataset is highly imbalanced—most perturbations exhibit phenotypes similar to the control distribution—the training set features were balanced by up-sampling perturbations with significant effects. Perturbations with significant effects were identified using a a chi-squared test, comparing infection states with and without perturbations, based solely on the training set.

### Evaluation

#### Single-cell RNA-seq data

We evaluated MORPH by comparing the distribution of real and predicted gene expression of perturbed cells. We used three metrics for this: *root mean squared error* (*RMSE*) of feature means, *Pearson correlation* between the predicted and true mean post-perturbation gene expression changes relative to the control, and *maximum mean discrepancy* (*MMD*) between the real and predicted gene expression distributions of the perturbed cells.

##### RMSE

It measures the root mean squared error between the means of the observed and predicted cell distributions as

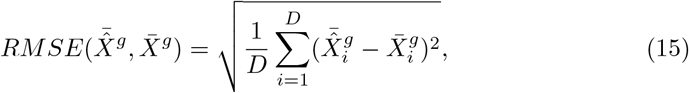

where 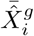 and 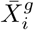 denote the predicted and observed mean of feature *i* in the perturbed cells (where gene *g* was perturbed), and *D* is the number of total features in a cell.

##### Pearson correlation (delta)

It quantifies the relationship between the observed and predicted mean changes from the training-set perturbed mean as

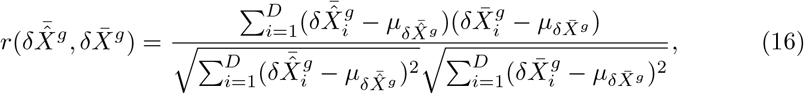

where 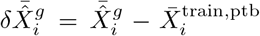 and 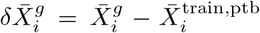 represent the predicted and observed mean changes in feature *i* following perturbation *g*, relative to its mean value across all *perturbed* cells in the training set, 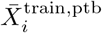. Here, 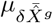 and 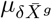 the averages of 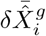 and 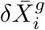 across features *i*, respectively. This metric follows the evaluation protocol of [27] and reduces the influence of global transcriptional shifts shared across perturbations (e.g., stress or cell-cycle effects), thereby emphasizing perturbation-specific structure.

##### Spearman correlation (delta)

We additionally computed the Spearman rank correlation to evaluate the agreement between predictions and observations. It is calculated identically to Equation 16, but replacing the raw predicted and observed mean changes, 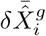 and 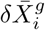, with their respective ranks across features.

##### Retrieval rank

For each perturbation *g*, let 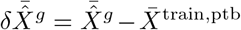 and 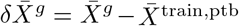 represent the predicted and observed mean changes following perturbation *g*, relative to the mean across all *perturbed* cells in the training set, 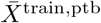. The retrieval rank of *g* is defined as

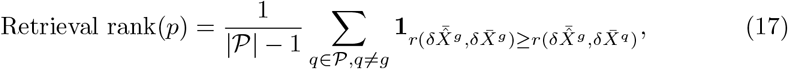

where *P* denotes the set of evaluated perturbations and *r* denotes the Pearson correlation. The score therefore represents the fraction of other perturbations whose correlation with the predicted effect of *g* does not exceed the self-correlation. Values range from 0 (worst) to 1 (best).

##### MMD

Metrics based solely on feature means can be insensitive to distributional heterogeneity, potentially favoring predictions that capture only the average rather than the full complexity of the underlying distribution. To address this limitation, we incorporated a distributional distance measure—maximum mean discrepancy (MMD) [58] —which captures differences beyond the mean by considering higher-order moments. We used an unbiased estimate of MMD by averaging kernel similarities over the cells in each set (Equation (6)). The Gaussian kernel was used, and the MMD is reported as an average over multiple bandwidths, similar to [56].

We reported all metrics using the top 50 marker genes as well as using all genes. Marker genes are computed for each perturbation with the scanpy [66] function `rank_genes_groups`, using the untreated control cells as reference.

#### Imaging data

We evaluated the model based on its ability to distinguish between biologically relevant cell states. Specifically, we computed the L1 loss between the predicted and observed cell state vectors for each perturbation, normalizing it by 2 to ensure a range between 0 to 1:

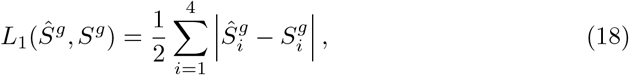

where *Ŝ*^*g*^ and *S*^*g*^ represent the predicted and observed cell state distribution vectors for cells subjected to perturbation *g*, with 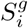 showing the proportion of perturbed cells in state *i*.

### Implementation details

We used PyTorch [68] to implement the MORPH neural network model. Hyperparameter search is discussed in Supplemental Note 1. Detailed computational budgets are provided in Supplemental Table 7. Pathway enrichment analysis was implemented using the GSEApy package [69].

## Footnotes

1 [24] proposed to use PCA-derived embeddings of target genes from the training data to predict the effect of unseen perturbations. However, we observed that DepMap embeddings capture more relevant information about gene-gene similarity with respect to their perturbation effects. To ensure that we compare our model to the strongest version of this baseline, we adapted the linear model to use DepMap embeddings instead.

2 We found that residual connections in the attention layers, while beneficial for prediction performance, diluted the signal when interpreting the gene regulatory network (GRN). Specifically, removing the residual connections in the first attention layer led to a clearer readout of program-to-gene mappings, improving interpretability at the cost of slightly reduced prediction accuracy (Methods, Supplemental Table 5 and Supplemental Note 3). Given that our primary interest here is GRN inference, we focus on this variant of MORPH for the remainder of the interpretability analysis. Results using the full model can be found in Supplemental Figure 13.

3 We used the second-to-last layer since the final layer tends to encode task-specific features, whereas the second-to-last layer captures more generalizable representations [28].

4 We set *N*_factor_ = 100 for the genome-wide Perturb-seq dataset. See Supplemental Note 1 for details.

5 We did not apply this to GEARS, since the input gene embeddings for GEARS are initialized randomly.

