## Supplemental PDF for "MORPH Predicts the Single-Cell Outcome of Genetic Perturbations Across Conditions and Data Modalities"

**This PDF file includes:**

Supplemental Notes 1, 2, 3, and 4,
Supplemental Tables 1 to 7,
Supplemental Figures 1 to 18.

### Supplemental Notes

#### 1 Hyperparameters

Hyperparameters were determined by searching on a validation set using a fixed split of the Norman dataset. The selected hyperparameters were applied consistently across all single-cell RNA sequencing datasets and different data splits. The search space for hyperparameters included the number of attention layers $-\{1, 2, 3\}$, latent dimension of the control encoder $-\{50, 200\}$, latent dimension of the perturbation encoder $-\{50, 200\}$, latent matrix dimension $-\{[20, 50], [50, 50], [100, 50]\}$, $\alpha - \{1, 2\}$, $\beta - \{0.0001, 2\}$, $\gamma - \{0, 1\}$, a learning rate of 0.0001, and a batch size of 32. The final selected hyperparameters for single-cell RNA-sequencing experiments were as follows: two attention layers, a latent dimension of 50 for both the control and perturbation encoders, a latent matrix dimension of $[50, 50]$, $\alpha = 2$, $\beta = 2$, $\gamma = 0$, a learning rate of 0.0001, and a batch size of 32. In the configuration file, this is equivalent to setting `latdim_ctrl = 50, latdim_ptb = 50`, `geneset_num = 50, geneset_dim = 50`, `mxAlpha=2`, `mxBeta=2`, and `Gamma1=0`. Early stopping was implemented, with training halted if the validation loss did not decrease for more than 20 epochs (i.e., `tolerance_epoch=20`). For the K562-GWPS dataset, hyperparameters were re-optimized on a validation set due to the increased complexity of modeling the massive perturbation space. The final values were a latent dimension of 200 for both the control and perturbation encoders, a latent matrix dimension of $[100, 50]$, $\alpha = 1$, $\beta = 2$ , and $\gamma = 1$. In the configuration file, this is equivalent to setting `latdim_ctrl = 200, latdim_ptb = 200`, `geneset_num = 100, geneset_dim = 50`, `mxAlpha=1`, `mxBeta=2`, and `Gamma1=1`. This adjustment was made to better capture the full perturbation space, as a larger model was required for effective representation.

For the imaging dataset, hyperparameters were similarly optimized using a validation set and applied across all data splits. The final selection included two attention layers, a latent dimension of 50 for both the control and perturbation encoders, a latent matrix dimension of $[20, 50]$, $\alpha = 2$, $\beta = 0.0001$, and $\gamma = 0$, a learning rate of 0.0001, and a batch size of 32. In the configuration file, this is equivalent to setting `latdim_ctrl = 50, latdim_ptb = 50`, `geneset_num = 20, geneset_dim = 50`, `mxAlpha=2`, `mxBeta=0.0001`, and `Gamma1=0`.

#### 2 Determining the gene interaction (GI) subtypes

Let $\bar{X}^g \in \mathbb{R}^D$ represent the mean feature vector when perturbing gene $g$, where $D$ is the dimension of the perturbation readout (e.g. RNA-seq or imaging). To quantify the change relative to the unperturbed control state ($\bar{X}^{\varnothing}$), we computed the difference:

$$\delta\bar{X}^g = \bar{X}^g - \bar{X}^{\varnothing}$$

For cells subjected to two perturbations $(a + b)$, we modeled their cell state changes as a linear combination of the individual perturbation effects:

$$\delta\bar{X}^{a+b} = c_a\delta\bar{X}^a + c_b\delta\bar{X}^b + \epsilon$$

where $c_a$ and $c_b$ are constants fitted to the data, and $\epsilon$ is an error term capturing deviations from the linear approximation. Following [1], we fitted this model using the Theil-Sen estimator (`TheilSenRegressor` with parameters: fit_intercept=False, max_subpopulation=1e5, n_subsamples=1000, max_iter=1000, and random_state=1000).

Using the fitted models, we then computed key metrics (Table 3). Here, $[\delta\bar{X}^a, \delta\bar{X}^b]$ denotes the concatenated matrix with the two single profiles as columns, and *dcor* refers to distance correlation [2]. Each metric was normalized to the same range by min-max scaling across all of the perturbations considered.

While [1] categorized different types of genetic interactions by applying UMAP to the computed metrics, using UMAP to group perturbations for benchmarking model predictions introduces variability due to its stochastic nature, sensitivity to hyperparameters, and lack of direct correspondence with GI metrics. Therefore, following [3], we computed GI scores based on the description of each GI type in [1] and determined clustering thresholds using the labeled GI subtypes provided in [1]. This allowed us to classify perturbations into distinct GI types (Table 4). Here, we incorporated 'equality of contribution' alongside 'magnitude' to define the synergy score, distinguishing it from potentiation. We used the top 2,500 highly variable genes to calculate these GI metrics.

### 3 Trade-off between prediction accuracy and interpretability

To analyze clusters of perturbations with similar effects (i.e., perturbation modules) and groups of genes that respond similarly to perturbations (i.e., gene programs), we compared the model with the best predictive performance ($f$) to a variant without the first residual connection in the initial attention layer ($f^{\mathrm{nr}}$). We did not observe any significant difference between the two in identifying modules and gene programs. However, when examining regulatory effects from perturbations to genes, $f^{\mathrm{nr}}$ produced clearer results. Two key factors contributed to the challenge of interpreting the attention maps.

First, in the full model $f$, we found that some regulatory effects were encoded in the perturbation embeddings ($Z^g$). Due to the residual connections, the attention maps could only capture residual information rather than direct regulatory effects, making it difficult to interpret perturbation effects on gene expression. This could even lead to a reversal of effect signs. Therefore, when our goal was to infer the bipartite gene regulatory network, we focused on a simplified model $f^{\mathrm{nr}}$ that removes the residucal connection in the first attention layer. In contrast, we used the full model $f$ when the goal was accurate prediction.

Second, MORPH has two attention layers, which means regulatory effects could be distributed across both layers. We observed that most regulatory effects were captured in the first layer, and thus focused our analysis on the first attention layer when inferring the gene regulatory network. Although this approximation can lead to mismatches between the scale of the first attention maps and the magnitude of changes in gene expression, the non-linearity between the two layers makes it difficult to combine their effects. The most interpretable model would use a single attention layer without residual connections - effectively removing the second attention layer in $f^{\mathrm{nr}}$. But this configuration resulted in lower predictive performance - another trade-off between MORPH's predictive accuracy and interpretability.

### 4 Identifiability of bipartite gene regulatory network

In this section, we show that our framework provably identifies the bipartite gene regulatory network between perturbation modules and gene programs, under mild assumptions. We formulate it as the causal disentanglement problem of identifying latent causal variables and the structure relating them to observed data. We then establish identifiability results under a set of mild assumptions, and finally discuss how this setup aligns with the framework proposed in the main paper.

Let $X \in \mathbb{R}^D$ denote observed variables generated from unobserved latent variables $U = (U_1, \ldots, U_p) \in \mathbb{R}^p$ through an unknown deterministic mixing function $f$, such that

$$X = f(U), \tag{1}$$

where the latent variables $U_1, \ldots, U_p$ are mutually independent. We define a bipartite causal graph $\mathcal{G} \subseteq U \times X$, where an edge $U_i \to X_j$ denotes a regulatory effect from the latent variable $U_i$ to the observed variable $X_j$. We assume that the mixing function $f$ is differentiable, and it defines regulatory effects through the Jacobian matrix $J_f(U)$, where a non-zero partial derivative $\partial f_j / \partial U_i \neq 0$ indicates an edge $U_i \to X_j$ in $\mathcal{G}$. In the context of Perturb-seq, each latent variable $U_j$ corresponds to a perturbation module (e.g., a group of perturbed genes with similar induced effects) and an edge $U_j \to X_i$ in the bipartite graph $\mathcal{G}$ encodes the regulatory effect from the perturbation module $U_j$ on gene $X_i$. The gene programs can be obtained by grouping genes $X_i$ that share similar mixing functions $f_i$.

We consider single-node soft interventions on latent variables; i.e., an intervention $g$ with target $i \in [p] := \{1, \ldots, p\}$ modifies the joint distribution $\mathbb{P}(U)$ into $\mathbb{P}^g(U)$ by changing the distribution $\mathbb{P}(U_i)$. Because of the destructive nature of sequencing, a cell can only be observed either before or after the perturbation and thus we focus on the setting where we have unpaired data from the observational distribution $\mathbb{P}(X^{\varnothing})$ and interventional distributions $\mathbb{P}(X^{g_1}), ..., \mathbb{P}(X^{g_K})$, where each $\mathbb{P}(X^{g_k})$ contains samples $X = f(U)$ under an intervention $g_k$.

With samples $X$ from $\mathbb{P}(X^{\varnothing}), \mathbb{P}(X^{g_1}), ..., \mathbb{P}(X^{g_K})$, the goal is to identify the latent variables $U$, the intervention targets $i_1, \ldots, i_K$, and the bipartite DAG $\mathcal{G}$, up to a suitable equivalence classes. Following [4], we consider the setting where we have at least one intervention per latent node, and make the following mild assumptions:

**Assumption 1.** The interior of the support of $\mathbb{P}(U)$ is non-empty in $\mathbb{R}^p$, and $f$ is a full row rank polynomial.

**Assumption 2** (Linear interventional faithfulness [4])**.** Let $g$ be an intervention with target $i \in [p]$. It holds that $\mathbb{P}(U_i + U_{[p]\setminus\{i\}}C^\top) \neq \mathbb{P}^g(U_i + U_{[p]\setminus\{i\}}C^\top)$ for all constant vectors $C \in \mathbb{R}^{|[p]\setminus\{i\}|}$.

Assumption 1 ensures that the latent variables have sufficient variability and that the mixing function $f$ preserves identifiability by being full row rank, preventing any latent variable from being completely masked in the observed data. In practice, we approximate $f$ using a stack of attention layers and MLPs. Furthermore, since $\mathbb{P}^g(U_i + U_{[p]\setminus\{i\}}C^\top = u) = \int_x \mathbb{P}^g(U_i = x)\mathbb{P}^g(U_{[p]\setminus\{i\}}C^\top = u - x)dx = \int_x \mathbb{P}^g(U_i = x)\mathbb{P}(U_{[p]\setminus\{i\}}C^\top = u - x)dx$, Assumption 2 ensures that the change after intervening on gene $g$ introduced in $\mathbb{P}^g(U_i = x)$ does not get erased after averaging by the distribution of $U_{[p]\setminus\{i\}}C^\top$. Thus Assumption 2 is a mild condition on the interventional changes that are allowed.

Since the latent variables $U$ are mutually independent, they can be re-indexed without introducing changes in the distribution of the observed the variables. This results in a trivial form of non-identifiability, which is formalized using the following definitions:

**Definition 1** (Causal disentanglement equivalence class [4]). The tuples $\langle U, i_1, ..., i_K\rangle$ and $\langle \hat{U}, \hat{i}_1, ..., \hat{i}_K\rangle$ are causal disentanglement (CD)-equivalent if and only if there exists a permutation $\pi$ of $[p]$, non-zero constants $\lambda_1, ..., \lambda_p \neq 0$, and $b_1, ..., b_p$ such that

$$\hat{U}_i = \lambda_{\pi(i)} U_{\pi(i)} + b_{\pi(i)}, \quad \forall i \in [p], \quad \text{and } \hat{i}_k = (i_k)_\pi, \quad \forall k \in [K].$$

**Definition 2** (Latent-index equivalence class of bipartite graphs). Two bipartite graphs $\mathcal{G}, \hat{\mathcal{G}} \subseteq U \times X$ are equivalent under a latent index permutation if there exists a permutation $\pi \in [p]$ such that

$$(U_i \to X_j) \in \mathcal{G} \quad \Leftrightarrow \quad (U_{\pi(i)} \to X_j) \in \hat{\mathcal{G}}.$$

**Theorem 1.** Under Assumptions 1 and 2, if each latent variable appears once in $(i_1, \ldots, i_K)$, then the tuple $\langle U, i_1, .., i_K\rangle$ is identifiable up to its CD-equivalence class, and the bipartite graph $\mathcal{G}$ is identifiable up to its latent-index equivalence class.

In the context of single-cell Perturb-seq data, Theorem 1 shows that the perturbation modules and the bipartite regulatory network between perturbation modules and gene programs are identifiable. We end this section with the proof of Theorem 1.

*Proof of Theorem 1.* First, we establish the identifiability of the intervention targets. With Assumptions 1 and 2, Theorem 1 in [4] shows that if each latent variable $i_k$ is the target of at least one soft intervention $g_k$, then the intervention target mapping $g_k \mapsto i_k$ is identifiable up to a permutation of the latent indices.

Since $U_i$ are mutually independent, Lemma 8 in [4] directly implies that $U$ is identifiable up to permutation and scaling. Thus, $\langle U, i_1, \ldots, i_K\rangle$ is identifiable up to its CD-equivalence class.

This means that there exists an invertible scaled permutation matrix $\Lambda \in \mathbb{R}^{p\times p}$ and a vector $b \in \mathbb{R}^p$ such that $\hat{U} = \Lambda U + b$. Denote $\Lambda = DP$, where $P$ is a permutation matrix corresponding to some permutation $\pi$ and $D = \text{diag}(\lambda_{\pi(1)}, .., \lambda_{\pi(p)})$ is a diagonal matrix with non-zero diagonal entries.

Let $\hat{f}$ be the function such that $X = \hat{f}(\hat{U}) = f(U) = \hat{f}(\Lambda U + b)$. By the chain rule, $J_f(U) = J_{\hat{f}}(\hat{U}) \cdot \Lambda$, where $J_f$ is the Jacobian matrix of $f$. Since $\Lambda = DP$, we have $J_f(U) = J_{\hat{f}}(\hat{U}) \cdot DP$. Concretely, for any $i \in [G], j \in [p]$, this gives $(J_f(U))_{i,\pi(j)} = \lambda_j (J_{\hat{f}}(\hat{U}))_{i,j}$. Because $\lambda_j \neq 0$, we have that $(J_f(U))_{i,\pi(j)} = 0$ if and only if $(J_{\hat{f}}(\hat{U}))_{i,j} = 0$. This means that $\mathcal{G}$ is identifiable up to a permutation of the latent indices.

Therefore, under Assumptions 1 and 2, we can identify the tuple $\langle U, i_1, \ldots, i_K\rangle$ up to its CD-equivalence class and the bipartite $\mathcal{G}$ up to its latent-index equivalence class. □

### Supplemental Tables

| Dataset | Methods | MMD ↓ | RMSE ↓ | Pearson correlation, delta ↑ | Retrieval rank, delta ↑ |
|---|---|---|---|---|---|
| Replogle et al. K562-EG | Control gene expression | 0.588 (0.009) | 0.220 (0.003) | 0.045 (0.028) | 0.533 (0.032) |
| | Geneformer | 0.584 (0.018) | 0.218 (0.007) | 0.063 (0.027) | 0.552 (0.034) |
| | GenePT (NCBI) | 0.505 (0.007) | 0.187 (0.003) | 0.242 (0.019) | 0.692 (0.018) |
| | GenePT (NCBI+UniProt) | 0.497 (0.008) | 0.184 (0.003) | 0.269 (0.014) | 0.711 (0.012) |
| | GPT (STRING) | 0.492 (0.007) | 0.183 (0.004) | 0.272 (0.021) | 0.712 (0.019) |
| | DepMap | 0.487 (0.01) | 0.181 (0.005) | 0.303 (0.040) | 0.748 (0.034) |
| | Mixture-of-experts | **0.486 (0.011)** | **0.179 (0.004)** | **0.311 (0.006)** | **0.749 (0.007)** |
| Replogle et al. RPE1-EG | Control gene expression | 0.834 (0.034) | 0.261 (0.009) | 0.118 (0.045) | 0.571 (0.038) |
| | Geneformer | 0.817 (0.025) | 0.257 (0.005) | 0.172 (0.04) | 0.622 (0.037) |
| | GenePT (NCBI) | 0.718 (0.04) | 0.227 (0.012) | 0.275 (0.031) | 0.694 (0.021) |
| | GenePT (NCBI+UniProt) | 0.689 (0.03) | 0.219 (0.008) | 0.312 (0.039) | 0.715 (0.027) |
| | GPT (STRING) | 0.712 (0.017) | 0.225 (0.005) | 0.275 (0.031) | 0.685 (0.029) |
| | DepMap | **0.653 (0.034)** | **0.211 (0.01)** | **0.367 (0.018)** | 0.750 (0.012) |
| | Mixture-of-experts | 0.673 (0.032) | 0.214 (0.009) | 0.355 (0.011) | **0.752 (0.011)** |

Table 1: 5-fold cross-validation results comparing different sources of prior knowledge (evaluated on top 50 DEGs). The Mixture-of-experts models combine priors from NCBI, STRING, and DepMap. Specifically, each prior is embedded into a latent space via a separate encoder, and another multilayer perceptron is used to merge them in the latent space. Values are reported as the mean with standard deviation in parentheses: Mean (SD).

| | Retrieval rank (DE) ↑ | Retrieval rank (whole genome) ↑ |
|---|---|---|
| Control distribution | 0.394 | 0.503 |
| Perturbed mean | 0.489 | 0.464 |
| Control gene masking (10%) | 0.786 | 0.785 |
| Control gene masking (90%) | 0.760 | 0.753 |
| Library-size rescaling (total = 20,000) | 0.776 | 0.773 |
| Library-size rescaling (total = 100,000) | 0.744 | 0.732 |

Table 2: Evaluations of model predictions under statistical perturbations of input data in the K562-EG dataset. Retrieval rank is reported for predictions evaluated on the top 50 differentially expressed (DE) genes and on the full gene set. Baselines of control distribution and perturbed mean are included for reference.

| Metric | Definition |
|---|---|
| Model fit | $dcor(c_a\delta\bar{X}^a + c_b\delta\bar{X}^b, \delta\bar{X}^{a+b})$ |
| Dominance (i.e., how much larger one coefficient is than the other) | $\lvert\log_{10}(\lvert c_a\rvert/\lvert c_b\rvert)\rvert$ |
| Magnitude, measuring the "strength" of interaction | $\sqrt{c_a^2 + c_b^2}$ |
| Similarity of single transcriptional profile to double transcriptional profile | $dcor([\delta\bar{X}^a, \delta\bar{X}^b], \delta\bar{X}^{a+b})$ |
| Equality of contribution | $\frac{\min(dcor(\delta\bar{X}^a,\delta\bar{X}^{a+b}),dcor(\delta\bar{X}^b,\delta\bar{X}^{a+b}))}{\max(dcor(\delta\bar{X}^a,\delta\bar{X}^{a+b}),dcor(\delta\bar{X}^b,\delta\bar{X}^{a+b}))}$ |

Table 3: Metrics used to define genetic interaction sub-types.

| GI type | Description in [1] | GI scores | Thresholds |
|---|---|---|---|
| Synergy | magnitude >> 1 | magnitude + equality of contribution | $> 1.128$ |
| Suppressors | $c_a, c_b$ both small | − magnitude | $> -0.383$ |
| Neomorphic | large $\epsilon$ | − model fit | $> -0.873$ |
| Redundant | $\delta\bar{X}^a, \delta\bar{X}^b, \delta\bar{X}^{a+b}$ all similar | similarity + equality of contribution | $> 1.685$ |
| Epistasis | $c_a >> c_b$ | − equality of contribution + dominance | $> -0.372$ |
| Potentiation | magnitude >> 1, $\delta\bar{X}^{a+b}$ dominated by one phenotype | magnitude − equality of contribution | $> 0.252$ |

Table 4: Genetic interaction scores and thresholds.

| Model (Metric) | Dataset (Mean ± SD) | | |
|---|---|---|---|
| | K562-EG from [5] (single-gene perturbation) | RPE1-EG from [5] (single-gene perturbation) | K562 from [1] (single- and double-gene perturbation) |
| $f$ (MMD, distribution loss) | **0.487 ± 0.010** | **0.653 ± 0.034** | **0.445 ± 0.025** |
| $f^{\mathrm{nr}}$ (MMD, distribution loss) | 0.495 ± 0.008 | 0.686 ± 0.020 | 0.479 ± 0.026 |
| $f$ (RMSE, feature means) | **0.181 ± 0.005** | **0.211 ± 0.010** | **0.178 ± 0.011** |
| $f^{\mathrm{nr}}$ (RMSE, feature means) | 0.185 ± 0.003 | 0.219 ± 0.005 | 0.195 ± 0.013 |
| $f$ (Pearson correlation, delta) | **0.303 ± 0.040** | **0.367 ± 0.018** | **0.629 ± 0.053** |
| $f^{\mathrm{nr}}$ (Pearson correlation, delta) | 0.286 ± 0.024 | 0.336 ± 0.029 | 0.609 ± 0.048 |

Table 5: Comparison of prediction performance with residual connections ($f$) and without residual connections ($f^{\mathrm{nr}}$) in the first attention layer, evaluated on top 50 DEGs.

| | Norman K562 | | Replogle K562-EG | | Replogle RPE1-EG | |
|---|---|---|---|---|---|---|
| Metric | No attn | Attn | No attn | Attn | No attn | Attn |
| MMD (DE) ↓ | 0.499 (0.038) | **0.445 (0.025)** | 0.489 (0.014) | **0.487 (0.010)** | 0.678 (0.021) | **0.653 (0.034)** |
| MMD (whole) ↓ | 0.336 (0.011) | **0.325 (0.009)** | **0.400 (0.001)** | 0.401 (0.001) | 0.447 (0.003) | **0.445 (0.002)** |
| RMSE (DE) ↓ | 0.186 (0.014) | **0.178 (0.011)** | 0.186 (0.006) | **0.181 (0.005)** | 0.214 (0.007) | **0.211 (0.010)** |
| RMSE (whole) ↓ | 0.040 (0.003) | **0.038 (0.002)** | 0.063 (0.001) | 0.063 (0.001) | 0.092 (0.002) | **0.091 (0.001)** |
| Pearson $r$ (DE) ↑ | 0.622 (0.048) | **0.629 (0.053)** | 0.297 (0.024) | **0.303 (0.040)** | 0.356 (0.034) | **0.367 (0.018)** |
| Pearson $r$ (whole) ↑ | 0.598 (0.034) | **0.629 (0.028)** | **0.225 (0.009)** | 0.219 (0.006) | 0.301 (0.022) | **0.310 (0.017)** |
| Retrieval rank (DE) ↑ | 0.844 (0.031) | **0.846 (0.033)** | **0.755 (0.013)** | 0.748 (0.034) | **0.750 (0.020)** | **0.750 (0.012)** |
| Retrieval rank (whole) ↑ | 0.889 (0.017) | **0.896 (0.011)** | 0.758 (0.020) | **0.760 (0.015)** | 0.726 (0.007) | **0.740 (0.016)** |

Table 6: Ablation of attention in MORPH. Comparison between MORPH with attention blocks and a capacity-matched variant in which attention blocks are removed and control-cell and perturbation latent embeddings are concatenated and fused via an MLP. Results are reported on the K562 cell line from Norman et al., the K562-EG dataset from Replogle et al., and the RPE1-EG dataset from Replogle et al., using 5-fold cross-validation. Values are reported as mean (standard deviation) across folds, with better values bolded.

| Dataset | Wall-clock time (hours) | Peak GPU memory (GB) |
|---|---|---|
| K562-EG [5] | 2.945 | 1.105 |
| RPE1-EG [5] | 2.406 | 1.105 |
| K562-GW [5] | 28.661 | 1.299 |
| Norman K562 [1] | 1.065 | 1.105 |

Table 7: Wall-clock time and peak memory consumption for training MORPH across evaluated datasets. All models were trained on a single NVIDIA RTX A6000 GPU, with peak memory tracked via `torch.cuda.max_memory_reserved`.

### Supplemental Figures

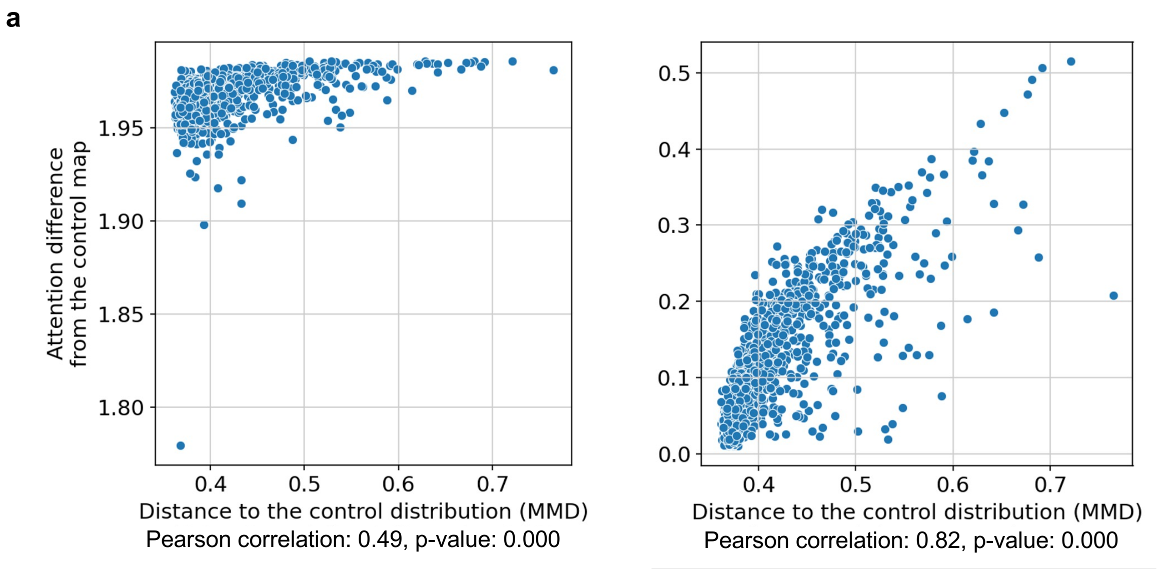


**Supplemental Figure 1**: **Effect of the control-distribution Maximum Mean Discrepancy (MMD) regularizer on attention interpretability. a,** Comparison of two distinct models trained on the RPE1-EG dataset [5] to evaluate different control cell reconstruction losses. Each point corresponds to a single perturbation. The x-axis shows the MMD between the perturbed and control cell distributions, quantifying the magnitude of each perturbation effect. The y-axis shows the difference in attention scores between the perturbed condition and the control reconstruction, computed as the sum of absolute differences in the first attention layer in MORPH. Results are shown for a model using Mean Squared Error (MSE) as the reconstruction loss for control cells (left), and a model using MMD as the reconstruction loss for control cells (right).

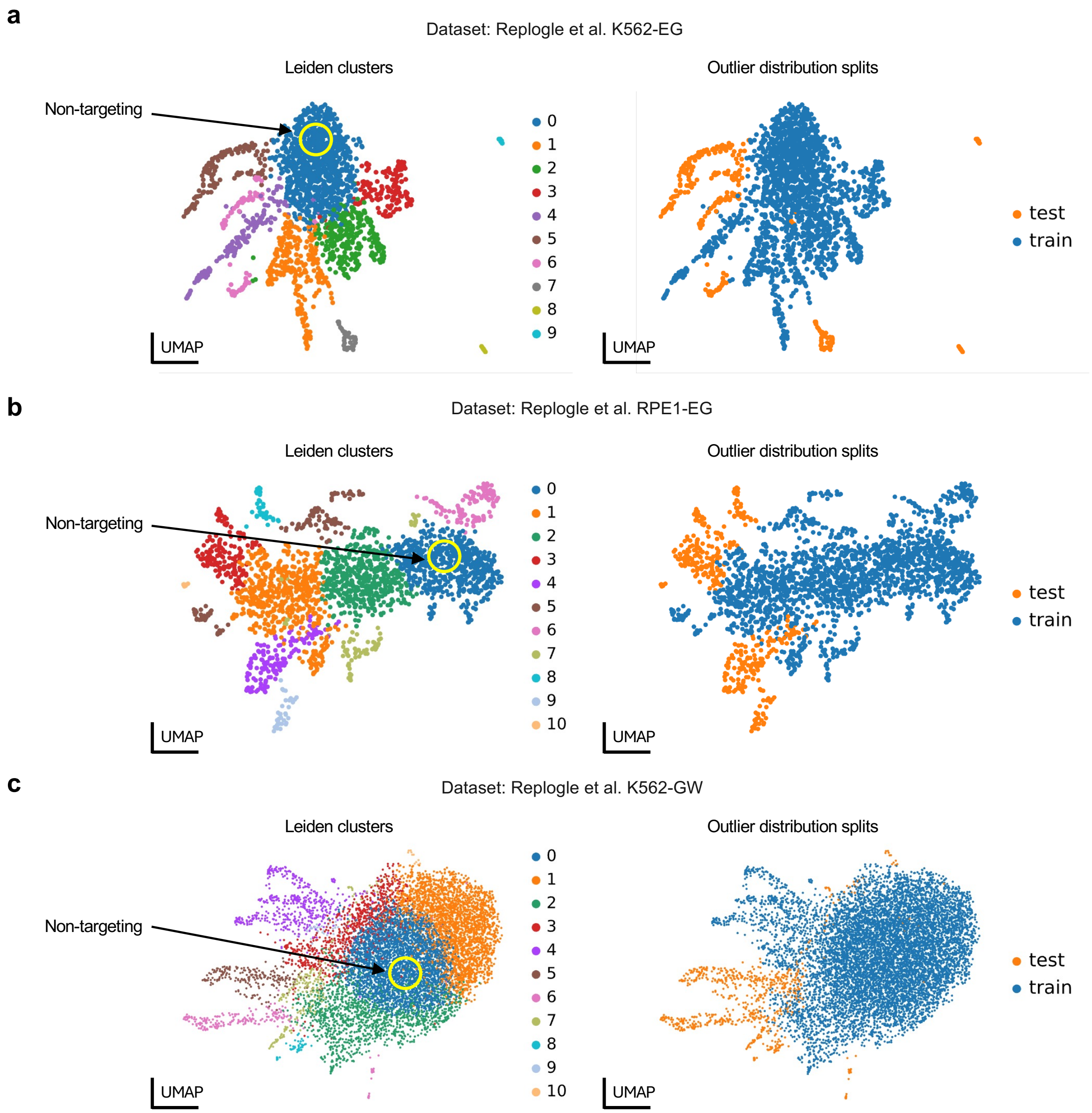


**Supplemental Figure 2**: **Illustration of outlier distribution splits.**

**a-c,** UMAP plots illustrating the outlier distribution splits for single-cell Perturb-seq data from [5] across different cell lines: K562 essential scale (ES) (**a**), RPE1 (**b**), and K562 genome-wide (GW) scale (**c**). Each point represents a perturbation, depicted by its pseudo-bulk profile. On the left, perturbations are colored by Leiden clusters. On the right, perturbations are colored based on whether they belong to the outlier distribution training or test set. The test set comprises perturbations from the five Leiden clusters that are farthest from the cluster containing non-targeting cells. Distances are calculated using the pseudo-bulk profiles to determine the highest Euclidean distance between cluster centers.

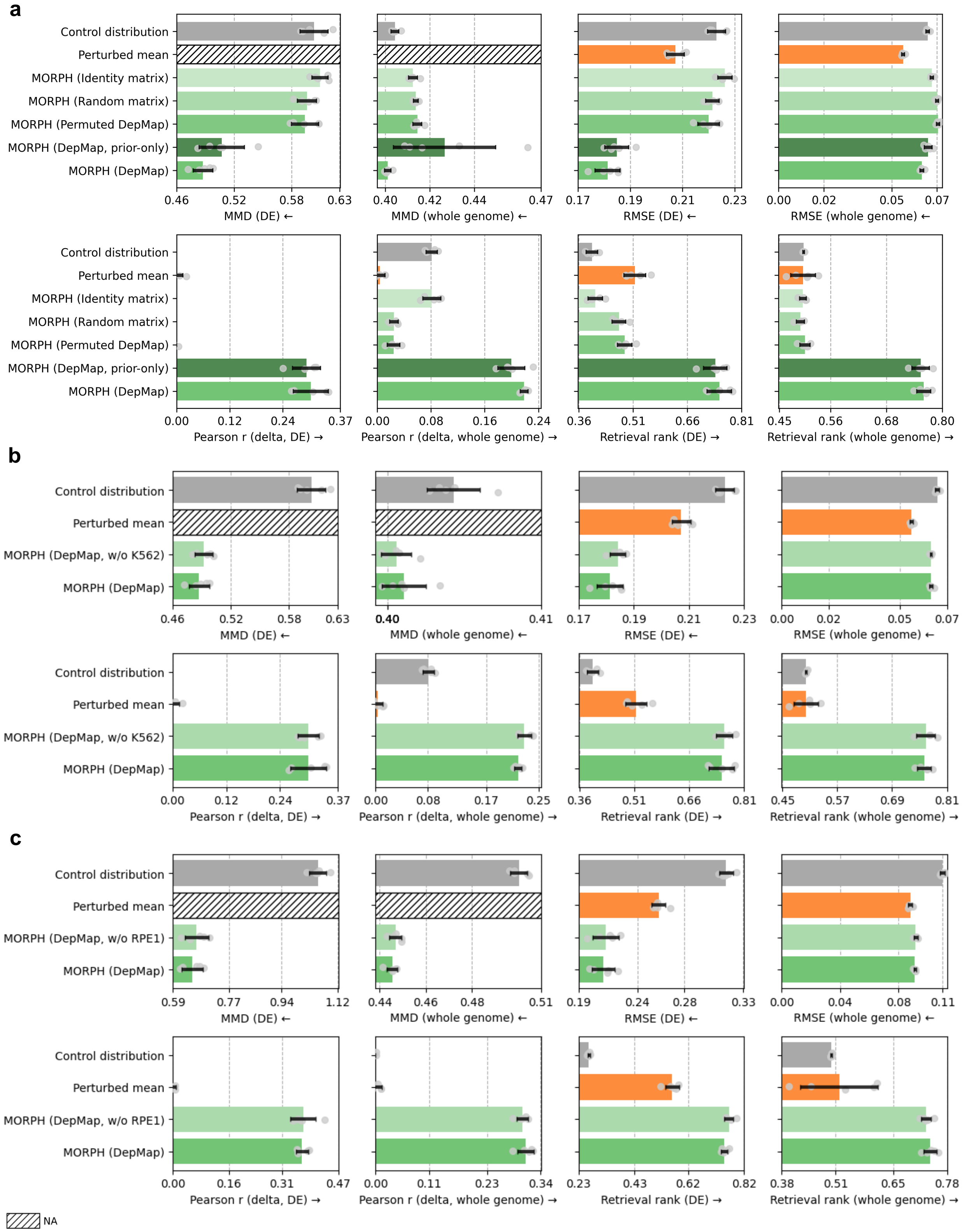


**Supplemental Figure 3**: **Ablation of gene embedding priors and control-cell inputs in MORPH. a**, MORPH's performance under ablations of the gene embedding prior or input control cells on the K562-EG dataset from [5]. Shown are results for models using informative DepMap gene embeddings, a prior-only variant with control-cell expression masked as zeros, and models using non-informative gene embeddings (identity, random, or permuted DepMap). **b,** Performance on the K562-ES dataset [5] after removing all K562 cell lines from DepMap prior to embedding construction. **c,** Performance on the RPE1-ES dataset [5] after removing all RPE1 lineage variants from DepMap prior to embedding construction. Two baseline predictors are included: randomly sampled control cells ("Control distribution") and the mean of perturbed cells from the training set ("Perturbed mean"). Performance is reported across multiple metrics—MMD, RMSE, Pearson r (delta), and retrieval rank—evaluated on the top 50 differentially expressed (DE) genes and across the whole genome. Pearson correlation values were clipped at 0 for visualization purposes. Hatched bars indicate metrics that are not applicable. MMD is not reported for Perturbed mean because it predicts only mean expression profiles and does not generate single-cell distributions. All methods were evaluated using five-fold cross-validation. Bars indicate mean performance across folds; points denote individual folds. Arrows indicate the direction of improvement ($\rightarrow$ higher is better; $\leftarrow$ lower is better).

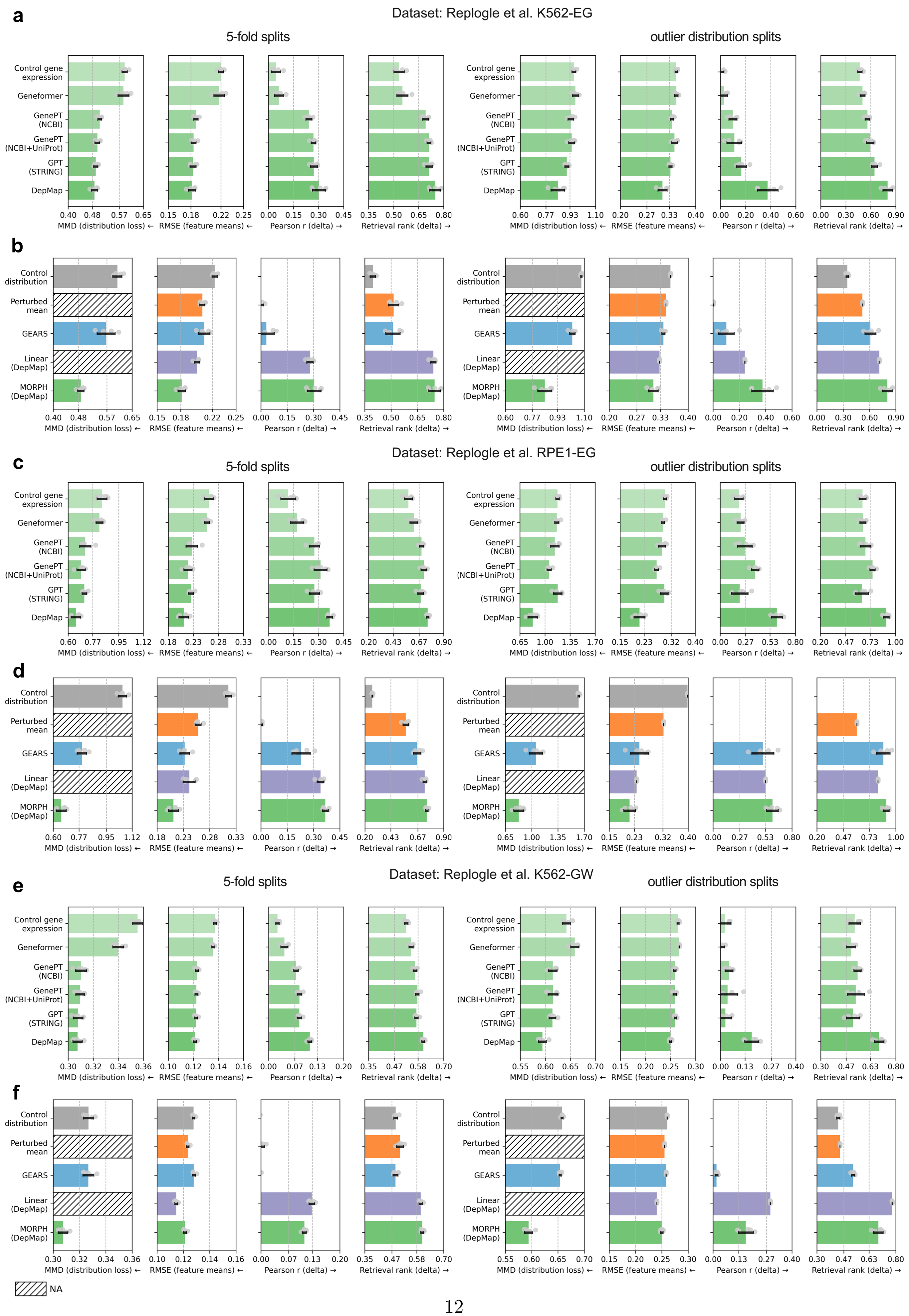
a
Dataset: Replogle et al. K562-EG
5-fold splits
outlier distribution splits
Control gene expression
Geneformer
GenePT (NCBI)
GenePT (NCBI+UniProt)
GPT (STRING)
DepMap
MMD (distribution loss) ←
RMSE (feature means) ←
Pearson r (delta) →
Retrieval rank (delta) →
b
Control distribution
Perturbed mean
GEARS
Linear (DepMap)
MORPH (DepMap)
c
Dataset: Replogle et al. RPE1-EG
d
e
Dataset: Replogle et al. K562-GW
f
NA

**Supplemental Figure 4**: **Evaluation of single-gene perturbation predictions on top 50 DE genes.**

**a-d**, Evaluation of single-gene perturbation predictions on the K562-EG dataset (**a-b**), the RPE1-EG dataset (**c-d**) and the K562-GW dataset (**e-f**) from [5]. Different sources of prior knowledge (**a, c, e**) and models (**b, d, f**) were evaluated using distributional distance (MMD), average RMSE between observed and predicted perturbed cells, Pearson correlation between the mean predicted and true post-perturbation gene expression changes relative to the training-set perturbed mean, and the retrieval rank. Pearson correlation values were clipped at 0 for visualization purposes. Hatched bars indicate metrics that are not applicable. MMD is not reported for Perturbed mean and the linear model because they predict only mean expression profiles and do not generate single-cell distributions. Metrics were calculated using the top 50 differentially expressed (DE) genes for each perturbation. Evaluations were performed across 5-fold cross-validation splits and outlier distribution splits. Arrows indicate the direction of improvement ($\rightarrow$ higher is better; $\leftarrow$ lower is better).

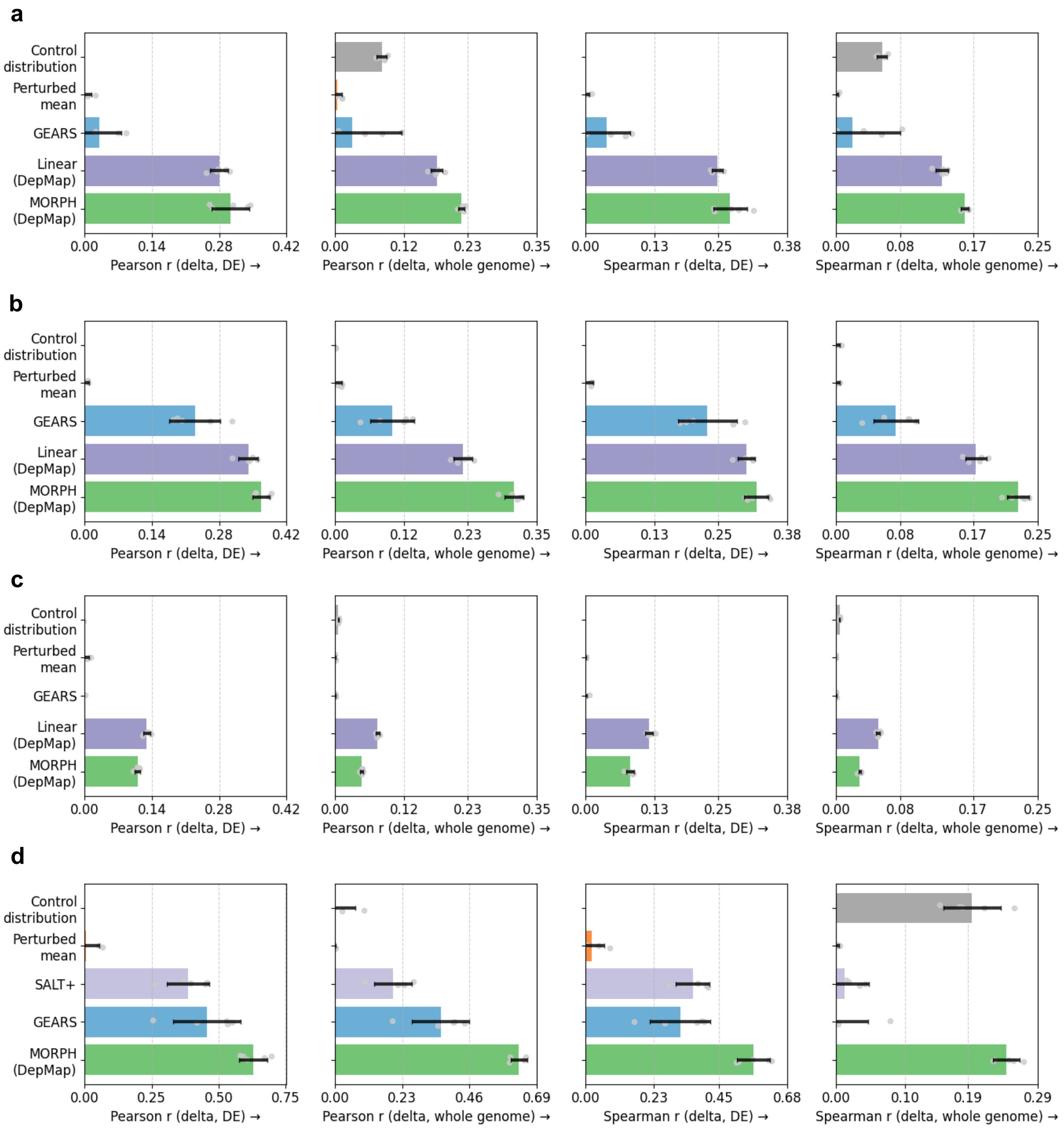


**Supplemental Figure 5**: **Evaluation of perturbation predictions on top 50 DE genes.**

**a-d,** Evaluation of single-gene perturbation predictions on the K562-EG (**a**), RPE1-EG (**b**), and K562-GW (**c**) datasets from [5], and of double-gene perturbation predictions on the Norman dataset (**d**) [1]. Models were evaluated using Pearson and Spearman correlation between the mean predicted and true post-perturbation gene expression changes relative to the training-set perturbed mean. Pearson and Spearman correlation values were clipped at 0 for visualization purposes. Metrics were calculated using the top 50 differentially expressed (DE) genes for each perturbation. Evaluations were performed across 5-fold cross-validation splits. Arrows indicate the direction of improvement (→ higher is better; ← lower is better).

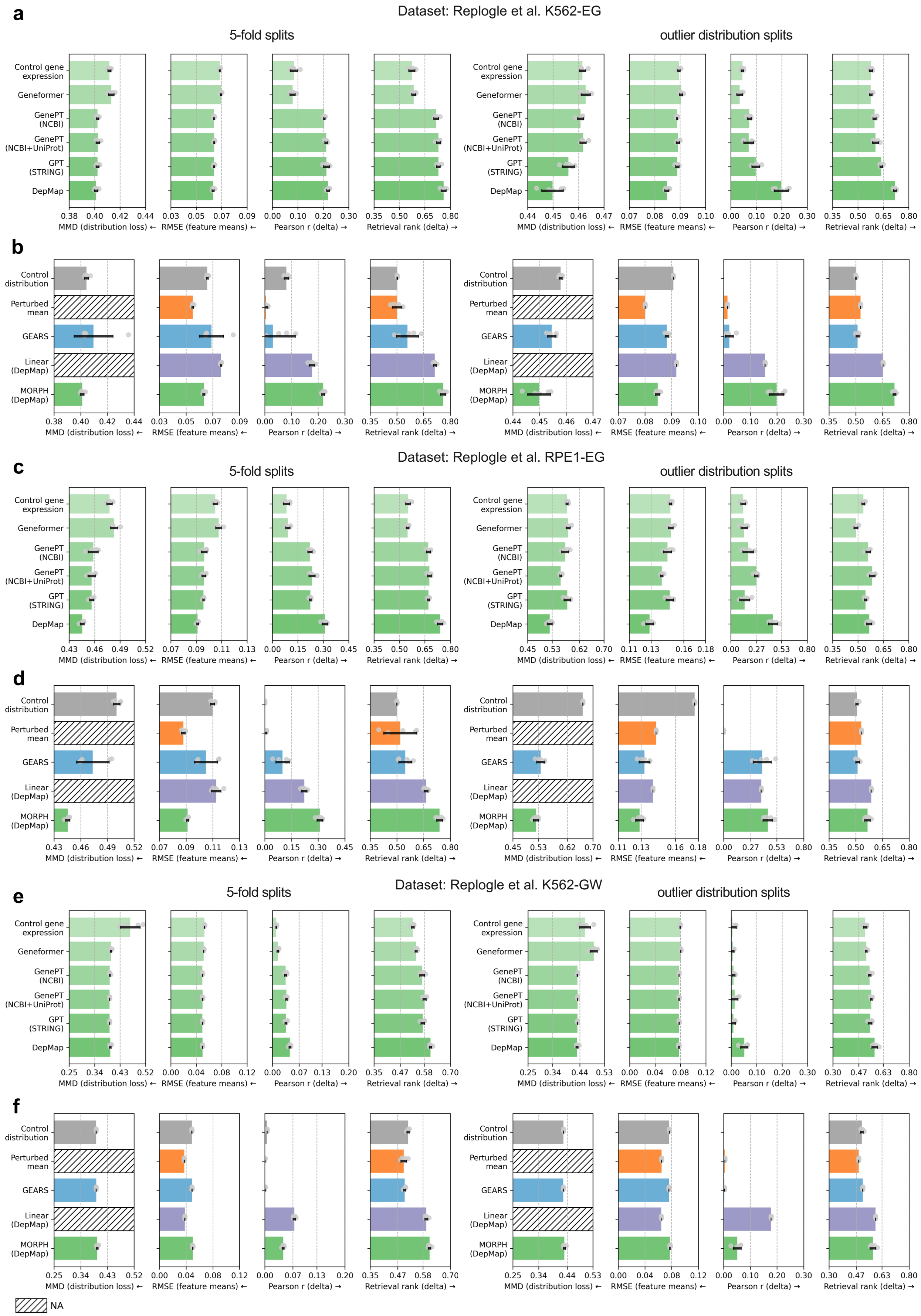
a
Dataset: Replogle et al. K562-EG
5-fold splits
outlier distribution splits
Control gene expression
Geneformer
GenePT (NCBI)
GenePT (NCBI+UniProt)
GPT (STRING)
DepMap
MMD (distribution loss) ←
RMSE (feature means) ←
Pearson r (delta) →
Retrieval rank (delta) →
b
Control distribution
Perturbed mean
GEARS
Linear (DepMap)
MORPH (DepMap)
c
Dataset: Replogle et al. RPE1-EG
d
e
Dataset: Replogle et al. K562-GW
f
NA

**Supplemental Figure 6**: **Evaluation of single-gene perturbation predictions on all genes.**

**a-f,** Evaluation of single-gene perturbation prediction performance across Perturb-seq datasets from [5]: K562-EG (**a-b**), RPE1-EG (**c-d**), and K562-GW (**e-f**). Different sources of prior knowledge (**a,c,e**) and models (**b,d,f**) were evaluated using distributional distance (MMD), average RMSE between observed and predicted perturbed cells, Pearson correlation between the mean predicted and true post-perturbation gene expression changes relative to the training-set perturbed mean, and the retrieval rank. Pearson correlation values were clipped at 0 for visualization purposes. Hatched bars indicate metrics that are not applicable. MMD is not reported for Perturbed mean and the linear model because they predict only mean expression profiles and do not generate single-cell distributions. Metrics were calculated using all genes for each perturbation. Evaluations were performed across 5-fold cross-validation splits and outlier distribution splits. Arrows indicate the direction of improvement ($\rightarrow$ higher is better; $\leftarrow$ lower is better).

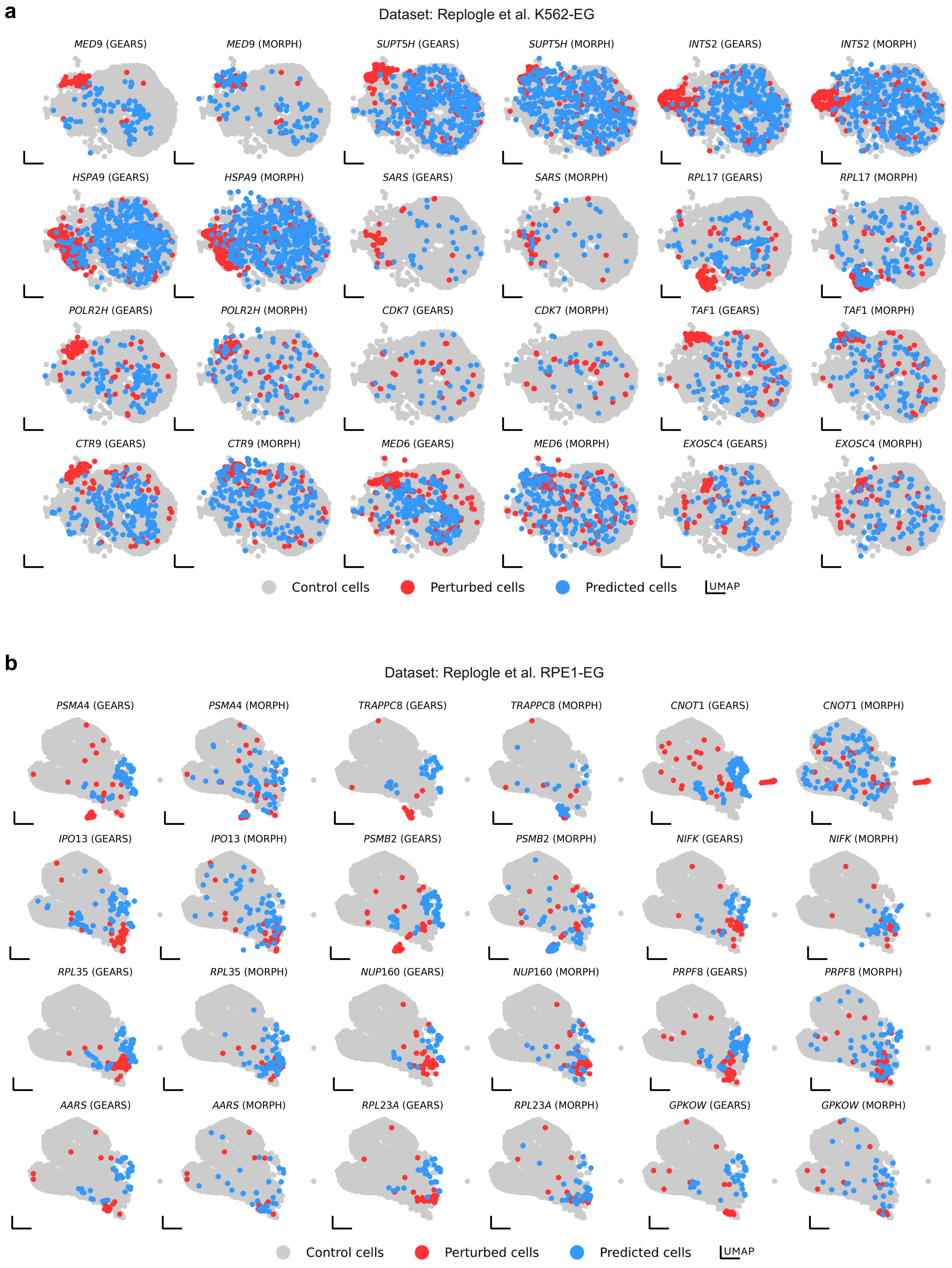


**Supplemental Figure 7: UMAP visualization of single-gene perturbation predictions.**

**a-b,** UMAP visualization of control cells, observed perturbed cells, and predicted perturbed cells from models trained on K562-EG dataset (**a**) and RPE1-EG dataset (**b**) from [5]. For both plots, the top 12 single-gene perturbations with the most pronounced effects (measured by the maximum mean discrepancy (MMD) between observed perturbed cells and control cells) in the test set are shown.

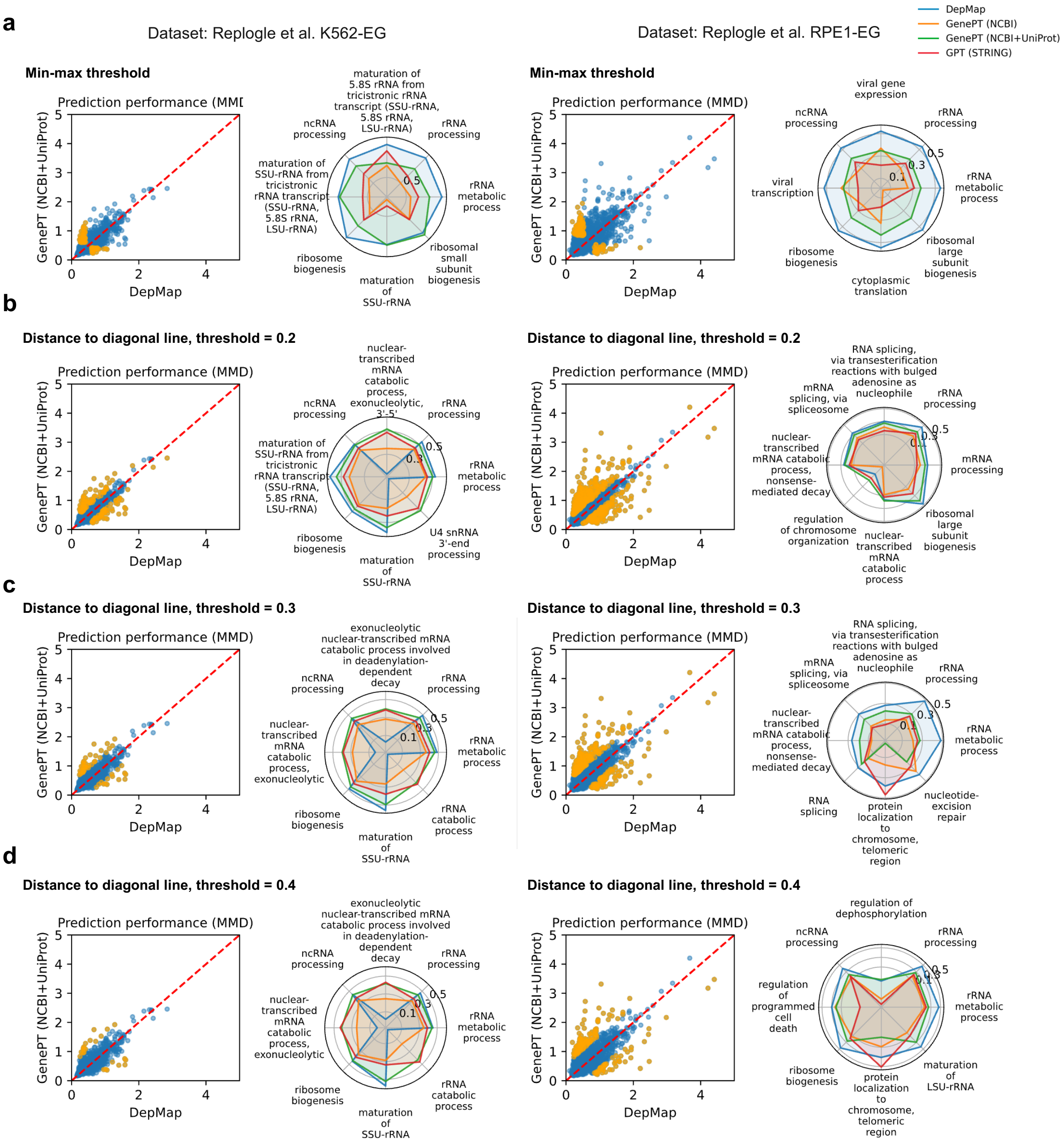


**Supplemental Figure 8**: **Sensitivity analysis of prior knowledge for perturbation embedding.**
**a,** Prior knowledge analysis performed on two datasets: K562-EG (left) and RPE1-EG (right) from [5]. Dot plots comparing the prediction performance of MORPH when trained with DepMap versus GenePT (the version that includes information from both NCBI and UniProt) as prior knowledge sources. Perturbations highlighted in yellow indicate cases where one prior significantly outperformed the other. Significant perturbations were identified using two thresholds: the lower bound of prediction loss, calculated as the mean of the minimum prediction loss across all priors, and a baseline defined by the mean prediction loss using only control gene expression. A perturbation was highlighted if one prior performed near the lower threshold while the other exceeded the baseline. This analysis was done for every pair of the top 5 strong priors that we found. Gene set enrichment analysis was performed on the union of these perturbations. The spider plot summarizes the mean prediction accuracy for each prior knowledge across the top 8 significantly enriched

gene sets. Prediction accuracy was calculated as $1 - \frac{MMD_{mean}}{max(MMD_{mean})}$, where $MMD_{mean}$ is the mean prediction loss, measured in MMD, of each gene set under a given prior, and $max(MMD_{mean})$ is the highest mean prediction loss across all gene sets. **b-d,** Similar analysis to **a** was performed, differing only in the method for identifying highlighted perturbations (cases where one prior outperformed the other). Here, perturbations were identified based on their distance to the diagonal line, which represents equal performance between the two priors. Different distance thresholds were used (0.2 in **b**, 0.3 in **c**, and 0.4 in **d**). All gene sets reported here were significantly enriched (adjusted p-value $< 0.05$), except for the RPE1 plots with threshold $= 0.3$ (**c**, right) and threshold $= 0.4$ (**d**, right). For these two plots, we reported the top 8 gene sets with the smallest adjusted p-values, which were 0.12 and 0.15, respectively.

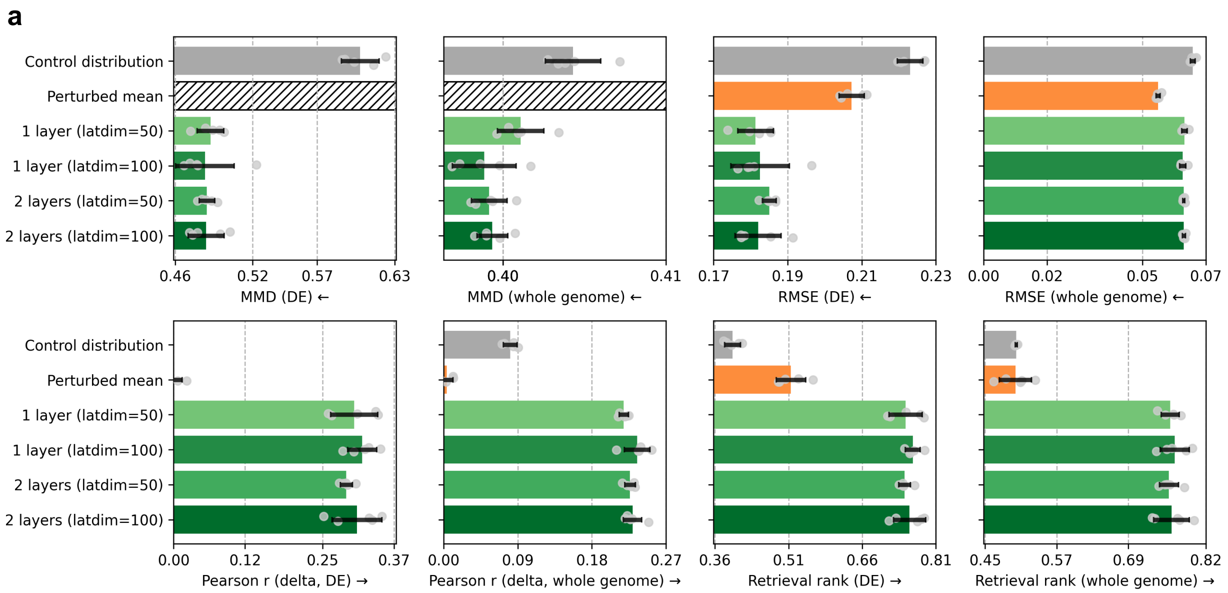


**Supplemental Figure 9**: **Robustness of MORPH to architectural choices and latent dimensionality. a,** MORPH performance under variations in model architecture, evaluated using five-fold cross-validation on the K562-EG dataset from [5]. We vary the perturbation and control-cell latent dimensionality (latent size = 50 or 100) and the number of decoder layers (one or two). Two baseline predictors are included for reference: randomly sampled control cells ("Control distribution") and mean of perturbed cells from the training set ("Perturbed mean"). Performance is reported across multiple metrics—MMD, RMSE, Pearson r (delta), and retrieval rank—evaluated both on the top 50 differentially expressed (DE) genes and across the whole genome. Pearson correlation values were clipped at 0 for visualization purposes. Hatched bars indicate metrics that are not applicable. MMD is not reported for Perturbed mean because it predicts only mean expression profiles and does not generate single-cell distributions. Bars indicate mean performance across cross-validation folds, and points indicate individual fold results. Arrows denote whether higher (→) or lower (←) values indicate better performance.

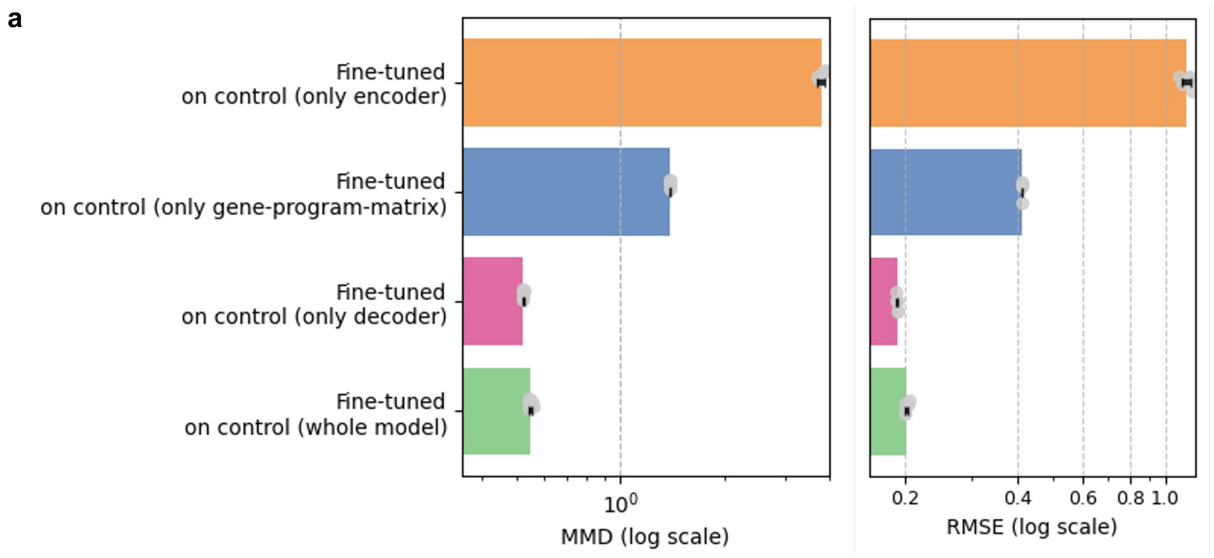


**Supplemental Figure 10**: **Ablation of fine-tuning strategies for transfer learning to a new cell line.**

**a,** Performance of different fine-tuning strategies for transfer from RPE1-EG to K562-EG [5], evaluated using Maximum Mean Discrepancy (MMD; left, log scale) and Root Mean Squared Error (RMSE; right, log scale) on perturbation predictions in the target cell line. Models were fine-tuned by updating only the encoder, only the decoder, only the gene-program matrix, or the entire model, while keeping all other components fixed. Bars represent the mean across five random seeds, with individual points indicating results from each seed.

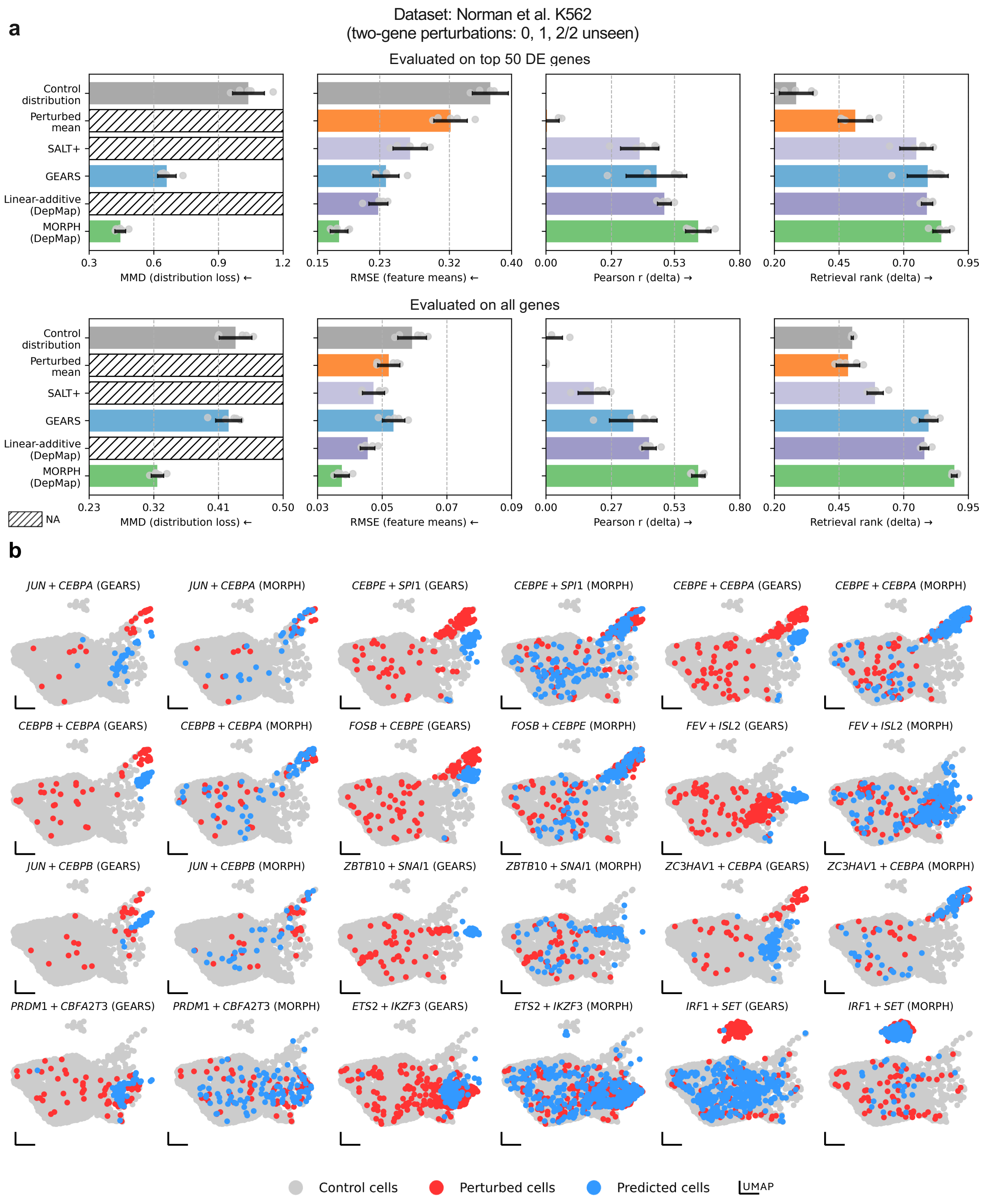


**Supplemental Figure 11: Evaluation of double-gene perturbation predictions.**

**a,** Evaluation of double-gene perturbation predictions on K562 data from [1]. Different models were assessed using distributional distance (MMD), average RMSE between observed and predicted perturbed cells,

Pearson correlation between the mean predicted and true post-perturbation gene expression changes relative to the training-set perturbed mean, and retrieval rank. Pearson correlation values were clipped at 0 for visualization purposes. Metrics were computed using either the top 50 differentially expressed (DE) genes per perturbation (left) or all genes (right). Evaluations were conducted using 5-fold cross-validation. Hatched bars indicate metrics that are not applicable. MMD is not reported for the Perturbed mean, SALT+ and linear-additive models, which predict only mean expression profiles. Arrows indicate the direction of improvement (→ higher is better; ← lower is better). **b,** UMAP visualization of control cells, observed perturbed cells, and predicted perturbed cells from models trained on data from [1]. The top 12 double-gene perturbations with the most pronounced effects (measured by the maximum mean discrepancy (MMD) between observed perturbed cells and control cells) in the test set are shown.

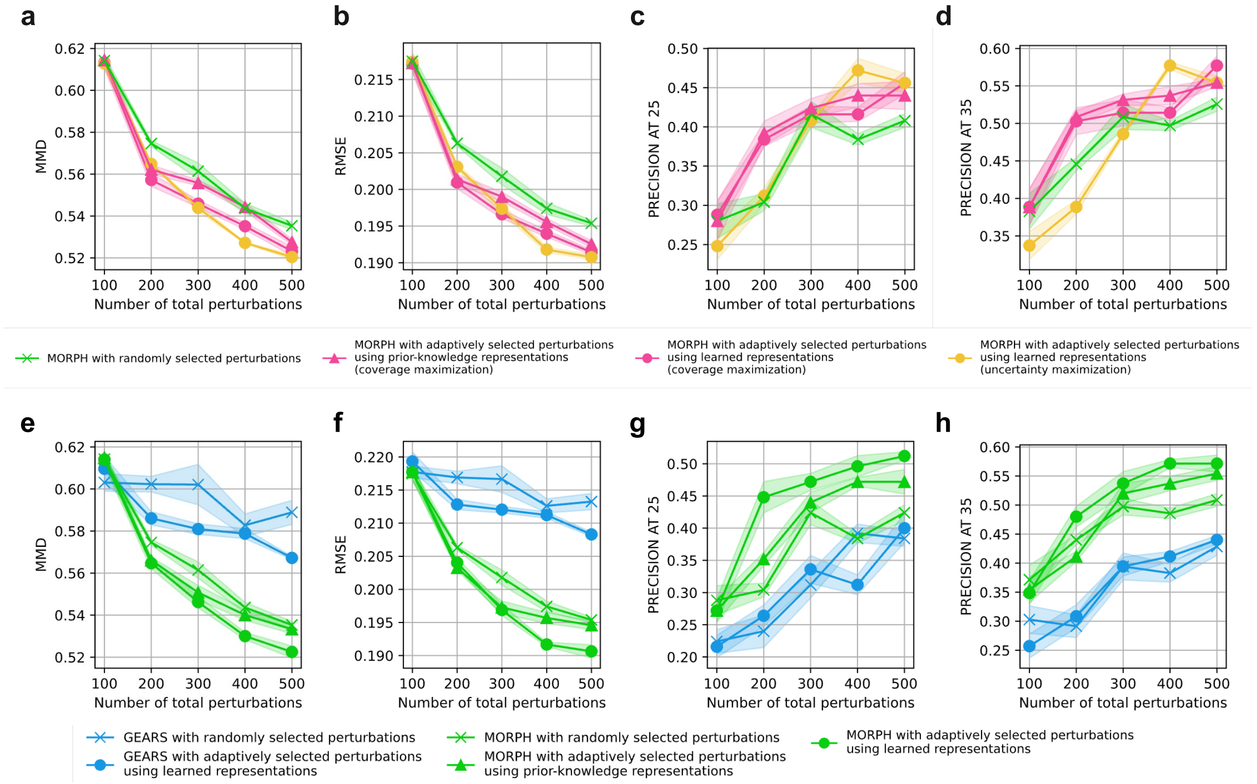


**Supplemental Figure 12**: **Perturbation experiment design with different strategies. a-d,** Line plots comparing acquisition strategies using MORPH for efficiently covering the perturbation space. "MORPH with adaptively selected perturbations using learned representations (coverage maximization)" (pink line with circles) selects perturbations closest to cluster centers in a model-learned latent space. For comparison, "MORPH with adaptively selected perturbations using prior-knowledge representations (coverage maximization)" (pink line with triangles) applies the same strategy in a fixed, prior-based latent space. "MORPH with adaptively selected perturbations using learned representations (uncertainty maximization)" (yellow line) prioritizes perturbations with the highest uncertainties measured in the model-learned latent space. Lines indicate mean performance and shaded regions show $\pm 0.2$ standard deviations across 5 runs with different random seeds. Evaluation metrics include MMD (**a**) and RMSE (**b**) calculated on the top 50 DE genes on a set of perturbations withheld for testing. **c-d** shows the precision of using the predictions to identify the top 25 and 35 perturbations that shifted cells farthest from the control state. **e-h**, Line plots comparing acquisition strategies and models for efficiently covering the perturbation space as measured by prediction over a held-out test set. The green line with 'X' markers represents MORPH with randomly selected perturbations. The green line with circle markers corresponds to MORPH using an adaptive strategy that selects perturbations with the highest predicted loss based on its learned latent representations. The green line with triangle markers shows a variant of MORPH that selects perturbations using a fixed, prior-based latent space. For comparison, the blue lines with 'X' and circle markers show GEARS with randomly selected and adaptively selected perturbations based on its learned latent representations, respectively. Lines indicate mean performance and shaded regions show $\pm 0.2$ standard deviations across 5 runs with different random seeds. Evaluation metrics include MMD (**a**) and RMSE (**b**) calculated on the top 50 DE genes on a set of perturbations withheld for testing. **c-d** shows the precision of using the predictions to identify the top 25 and 35 perturbations that shifted cells farthest from the control state.

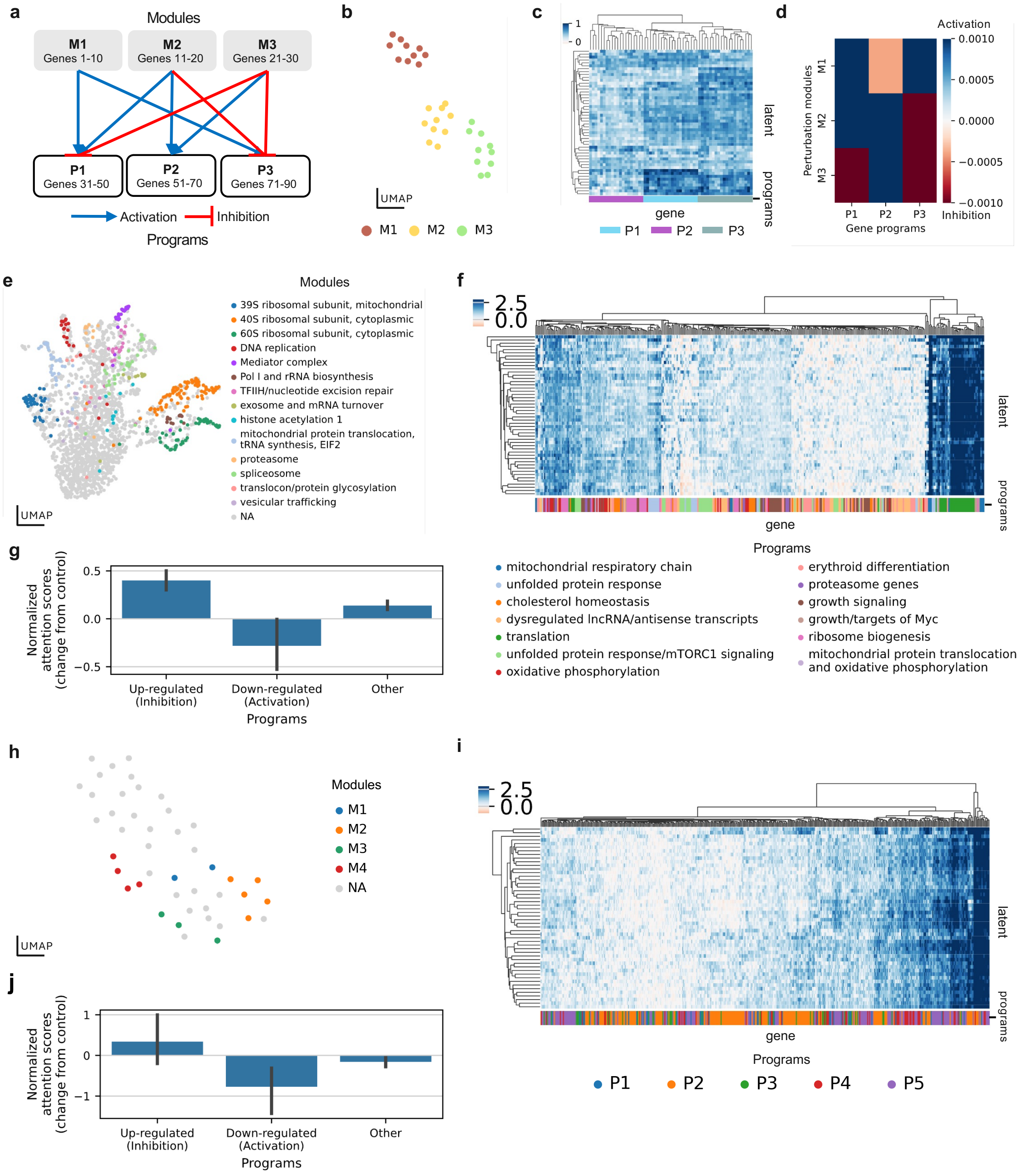


**Supplemental Figure 13**: **Gene regulatory network inference using model with residual connections on the first attention layer.**

**a-d**, Gene regulatory network analysis on simulated data. **a**, The gene regulatory network used to generate the data, where perturbation modules (M) are clusters of genes that induce similar effects when perturbed and gene programs (P) are genes that exhibit similar response to perturbations. **b**, UMAP visualization of the learned perturbation latent space colored by perturbation modules. **c**, Hierarchically clustered genes into programs using the learned mapping from programs to genes, colored by true gene programs. **d**, Recovered gene regulatory structures inferred from the learned attention maps. **e-g**, Same analysis for a model trained on K562-EG [5]: **e**, UMAP of the learned perturbation latent space colored by gene modules reported in [5]; **f**, Hierarchically clustered heatmap of a subset of the gene–latent attribution matrix **M**, in which rows represent latent programs and columns represent genes. For visualization, we display only genes annotated in curated gene programs from [5]; annotated gene programs are indicated by the color bar; **g**, box plots of rank-normalized attention score changes for up- and down-regulated programs post-perturbation. **h-j,** Same analysis for a model trained on the perturbed bone-marrow derived dendritic cells (BMDCs) from [6]. **h**, UMAP of the learned perturbation latent space colored by gene modules reported in [6]; **i**, Hierarchically clustered heatmap of a subset of the gene–latent attribution matrix **M**, in which rows represent latent programs and columns represent genes. For visualization, we display only genes annotated in curated gene programs from [6]; annotated gene programs are indicated by the color bar; **j**, box plots of rank-normalized attention score changes for up- and down-regulated programs post-perturbation.

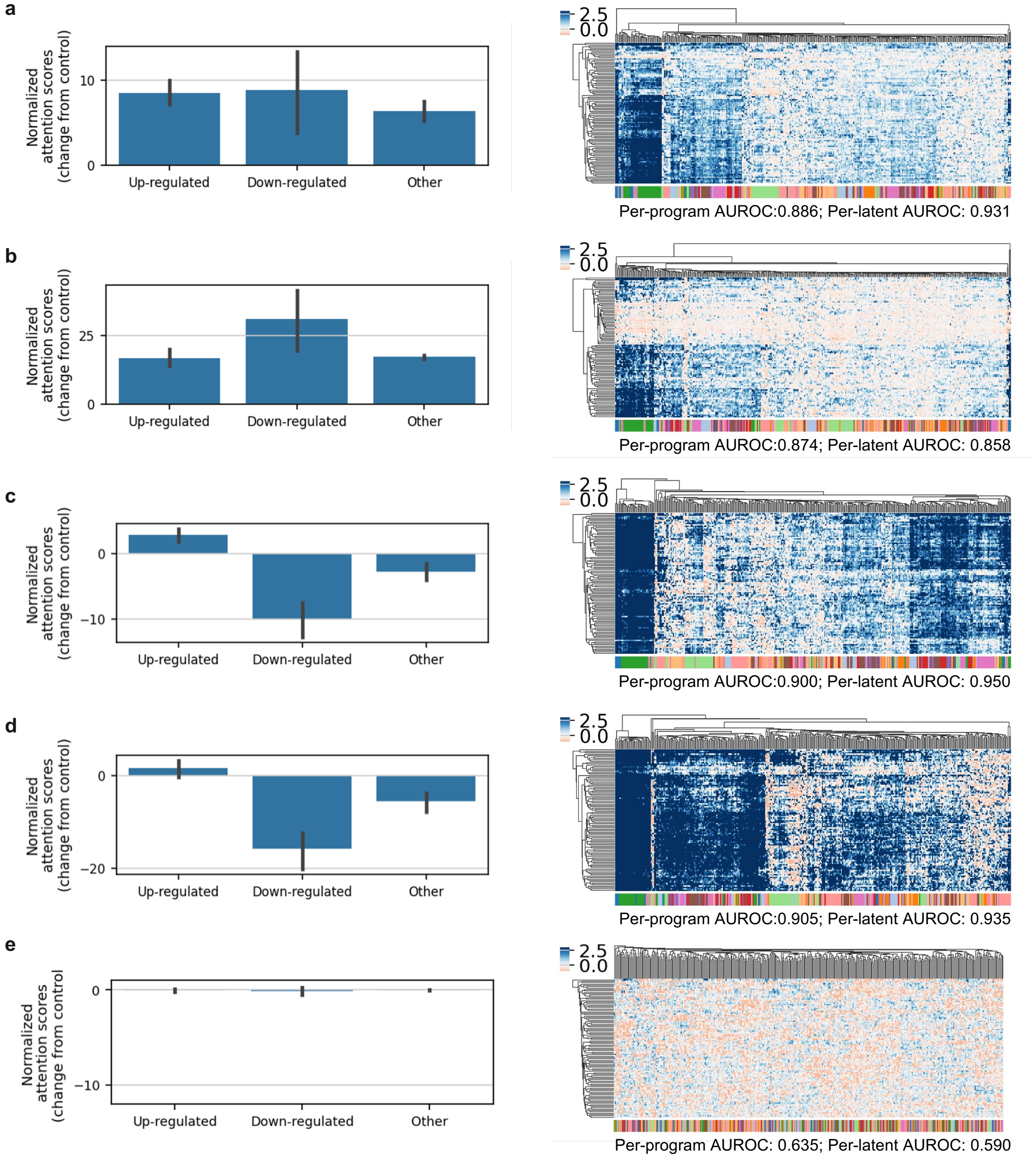


**Supplemental Figure 14**: **Evaluations of attention scores and gene program discovery under statistical perturbations of input data and model randomization.**

**a**–**d**, Evaluation of models retrained on the K562-EG dataset under four perturbation settings: random masking of 10% (**a**) or 90% (**b**) of genes in control cells; and library-size rescaling to total counts of 20,000 (**c**) or 100,000 (**d**). **e**, Evaluation of a model with randomized weights. **Left,** box plots of rank-normalized attention score changes for reported up- and down-regulated programs post-perturbation from [5]. **Right,** the gene–latent attribution matrix (**M**) inferred from each retrained model, displayed as hierarchically clustered heatmaps (rows, latent dimensions; columns, genes). For visualization, only genes from curated programs in [5] are shown (annotations indicated by the colour bar). Alignment between learned latent programs and annotated gene sets is quantified by an AUROC-based best-match assignment across all genes (see Methods), with numerical values indicated.

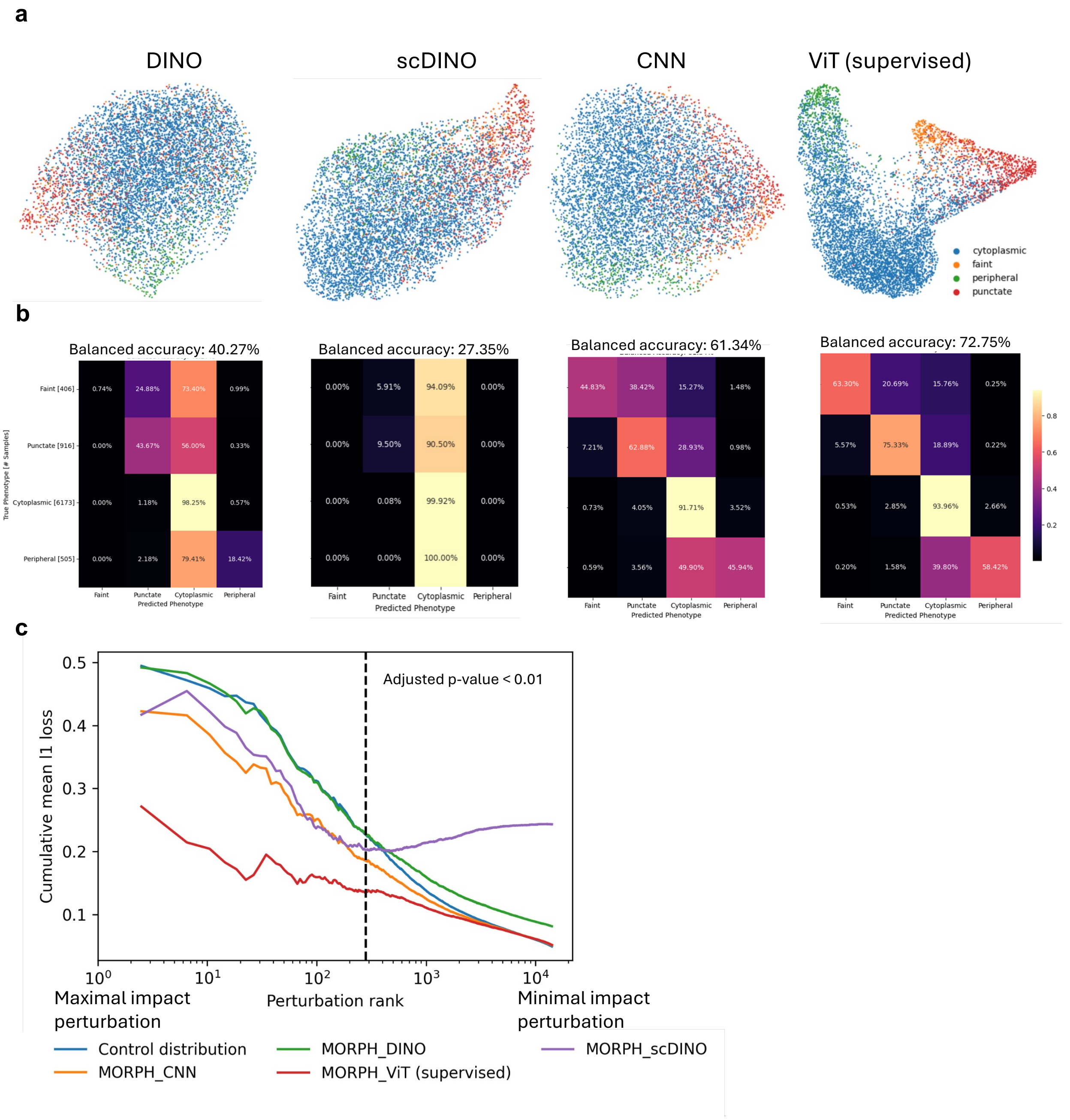


**Supplemental Figure 15**: **Comparison of image representations for downstream perturbation modeling.**

**a,** UMAP visualizations of extracted image feature representations colored by infection state (faint, peripheral, cytoplasmic, punctate) for different image encoders (DINO [7], scDINO [8], CNN, and supervised ViT). Each point represents a single cell. **b,** Confusion matrices and balanced accuracy of a simple classifier trained on top of each representation to predict infection state, using ground-truth labels. Balanced accuracy values are reported above each matrix. **c,** Model performance evaluations using normalized $\ell_1$ loss between predicted and true infection state distribution vectors. Perturbations on the x-axis are ranked by their impact on infection states based on chi-squared test, with genes having maximal impact on the left and minimal impact on the right. The y-axis shows the cumulative mean $\ell_1$ loss for each method. The dotted line indicates perturbations with a Bonferroni-corrected p-value $< 0.01$.

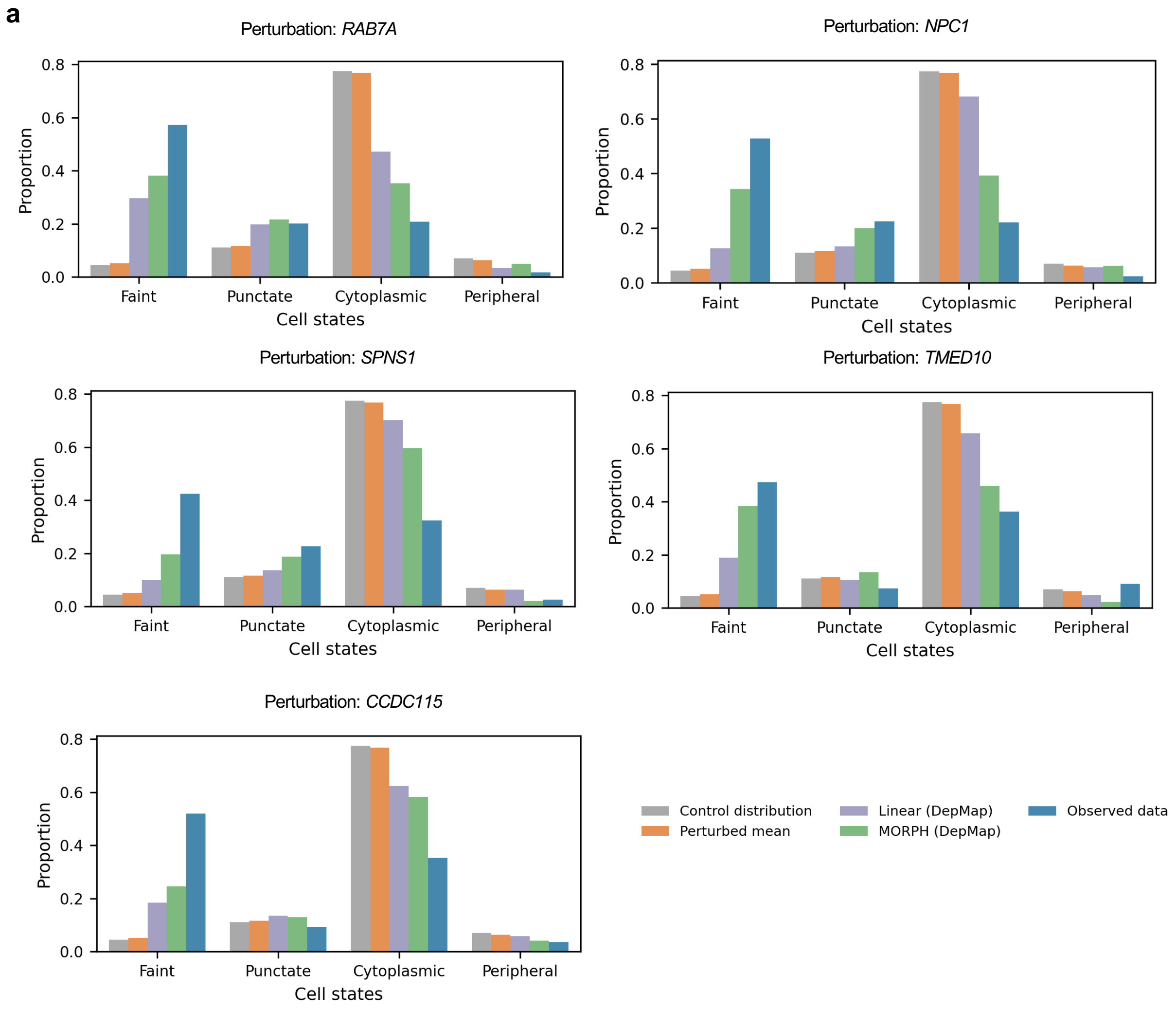


Scatter plot of chi-squared p-values versus L1 loss predicted by our model. Each dot represents a perturbation, colored by gene set. The dashed line indicates the Bonferroni-corrected significance threshold (p-value < 0.01).

**Supplemental Figure 16**: **Evaluation of prediction performance on imaging data.** **a**, Box plots comparing the predicted infection states of the top five perturbations with the greatest impact on cell state changes across different methods. The observed infection states (blue) represent the ground truth measurements.

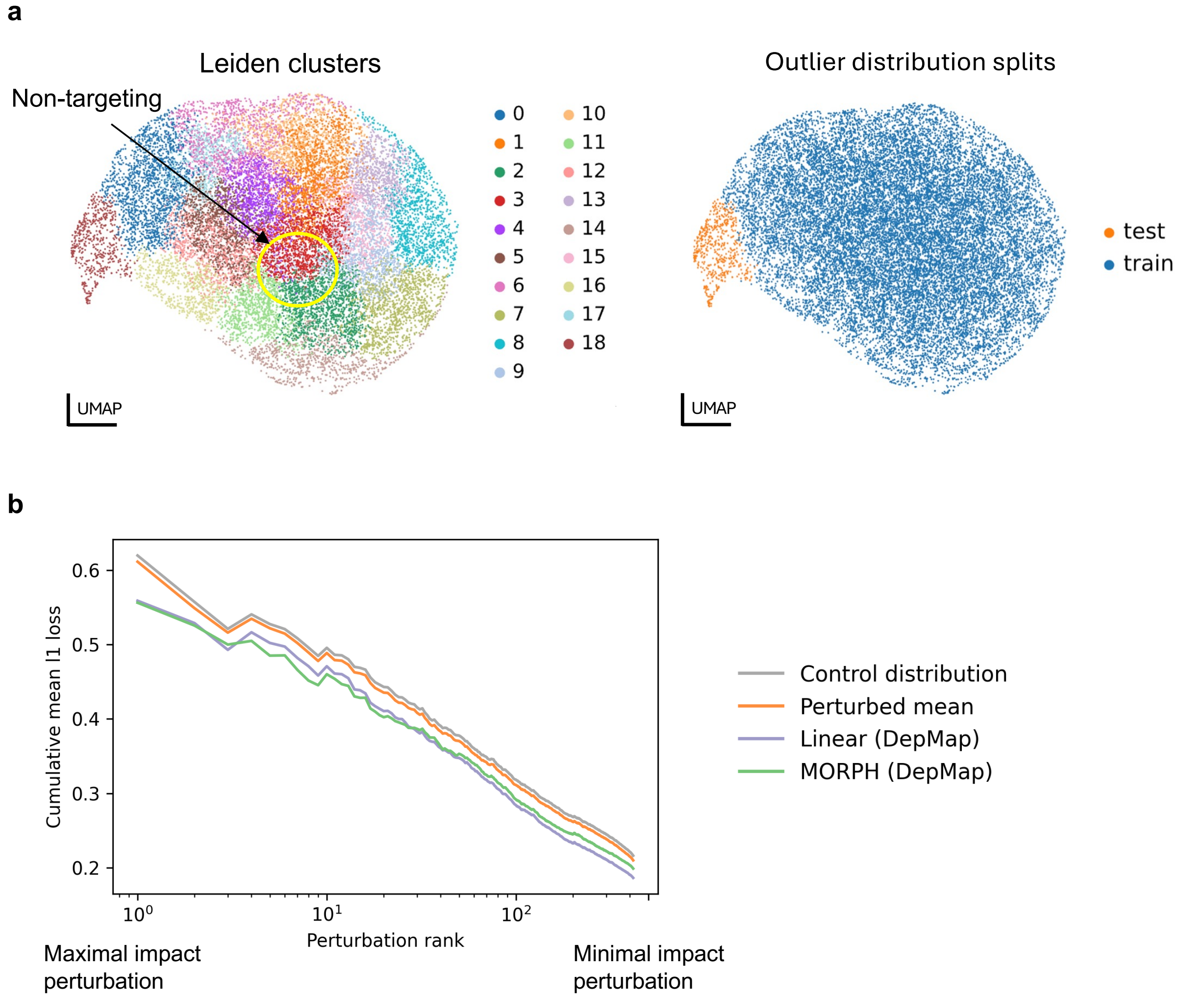


**Supplemental Figure 17**: **Model performance evaluation on optical pooled screening data using outlier distribution splits.**

**a,** UMAP visualization of perturbations in the optical pooled screening dataset, where each point represents a perturbation summarized by its pseudo-bulk imaging feature profile. Left: perturbations colored by Leiden clusters, with the cluster containing control cells receiving non-targeting guide RNAs indicated. Right: outlier distribution split, with perturbations colored by training or test assignment. The test set consists of perturbations belonging to the Leiden cluster whose center is farthest from the control-containing cluster, measured by Euclidean distance between cluster-level pseudo-bulk profiles. **b,** Cumulative mean normalized $\ell_1$ loss as a function of perturbation rank (ordered by impact), comparing MORPH, a linear baseline, and control-distribution and perturbed-mean baselines under the outlier distribution split.

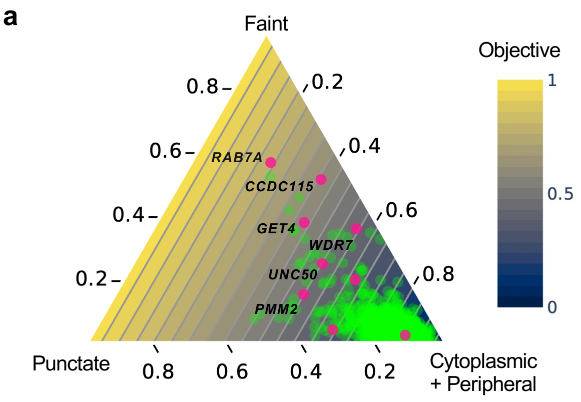


**Supplemental Figure 18**: **Evaluation of MORPH for target nomination.**

**a,** Ternary plot showing the ground-truth cell state distributions for genetic perturbations. The three axes represent the proportion of cells in the "Faint", "Punctate", and combined "Cytoplasmic" and "Peripheral" stages of infection. The background gradient visualizes the objective function, defined as the early-state fraction (the sum of "Faint" and "Punctate" states). Green points represent the 3,000 perturbations from the simulated expert-guided pilot screen (training set). Pink points indicate the top 10 candidate genes nominated by MORPH from the 14,973 held-out perturbations.
